# INFIMA leverages multi-omics model organism data to identify effector genes of human GWAS variants

**DOI:** 10.1101/2021.07.15.452422

**Authors:** Chenyang Dong, Shane P. Simonett, Sunyoung Shin, Donnie S. Stapleton, Kathryn L. Schueler, Gary A. Churchill, Leina Lu, Xiaoxiao Liu, Fulai Jin, Yan Li, Alan D. Attie, Mark P. Keller, Sündüz Keleş

## Abstract

Genome-wide association studies have revealed many non-coding variants associated with complex traits. However, model organism studies have largely remained as an untapped resource for unveiling the effector genes of non-coding variants. We develop INFIMA, **In**tegrative **Fi**ne-**Ma**pping, to pinpoint causal SNPs for Diversity Outbred (DO) mice eQTL by integrating founder mice multi-omics data including ATAC-seq, RNA-seq, footprinting, and *in silico* mutation analysis. We demonstrate INFIMA’s superior performance compared to alternatives with human and mouse chromatin conformation capture datasets. We apply INFIMA to identify novel effector genes for GWAS variants associated with diabetes. The results of the application are available at http://www.statlab.wisc.edu/shiny/INFIMA/

## 1 Introduction

Vast majority of disease and complex human trait-associated single nucleotide polymorphisms (SNPs) identified through genome-wide association studies (GWAS) are non-coding^[1]^. This creates two key challenges for translation of genetic discoveries into disease mechanisms. GWAS have capitalized on large-scale genomic and epigenomic data to address the first challenge of interpreting non-coding risk SNPs and assigning them potential regulatory roles^[2,3]^. In many cases, non-coding loci with risk SNPs span broad genomic regions that contain multiple genes^[4]^. This creates the second challenge of identifying the effector genes through which risk SNPs exert their impact on the phenotype, possibly via long-range chromatin interactions. With the advances in three-dimensional (3D) chromatin structure and interaction profiling, recent studies have successfully shown that a genetic variant is not necessarily causal for the nearest gene^[5,6]^. The consequence of this new perspective is a vast expanse of the set of candidate effector genes for a GWAS risk locus. In addition, the linkage disequilibrium (LD)^[7]^ further complicates the elucidation of effector genes for most GWAS risk SNPs because the causal variant may not be the SNP with the strongest association, but one that is in high LD. Collectively, these challenges hinder the delineation of effector genes for the majority of GWAS risk SNPs.

The recent transcriptome-wide association studies (TWAS) that leverage reference expression panels led to notable progress in identifying candidate disease-associated genes^[8,9]^. However, these approaches do not directly link the effector genes to SNPs. In addition, and perhaps more restrictively, they rely on reference transcriptomes which may not be readily available or are difficult to obtain for an array of disease-relevant tissues. Complementary to these, model organism studies continue to provide opportunities to unveil susceptibility genes and investigate findings from human GWAS. Specifically, progress during the last decade confirmed that evolutionary conservation can be used to discover regions of coding and non-coding DNA that are likely to have biological functions^[10–12]^, and thus may harbor functional SNPs. In this paper, we leverage model-organism multi-omics data, specifically, data from the Diversity Outbred (DO) mouse population^[13]^, to develop a framework for identifying candidate effector genes of non-coding human GWAS SNPs.

The DO mouse population^[13]^, a model organism resource derived from eight founder strains (129, AJ, B6, CAST, NOD, NZO, PWK, WSB), has been widely used to identify QTL for a variety of physiological and molecular phenotypes, including type 2 diabetes and gene expression in pancreatic islets^[14–18]^. These studies led to novel insights into the genetic architecture of islet gene regulation^[14]^ and insulin secretion^[19]^. However, a key impediment to maximizing the results of these types of eQTL studies is the lack of genomic resolution required to pinpoint the causal variants, and elucidate potential regulatory mechanisms. These inbred genomes harbor long stretches of genetic variants in high LD^[20]^. While this is advantageous for achieving gene-level mapping because, compared to a human GWAS, comparatively fewer markers (i.e., tag SNPs) are needed to genotype a larger group of SNPs, it results in groups of SNPs with similarly high LOD scores. Consequently, it hinders identifying enhancer-sized regions (i.e., in the order of hundreds of bases) underlying the detected associations. For example, an eQTL marker with the highest LOD score was identified for the gene *Abcc8* (Fig. 1a), where PWK has the lowest allelic effect (Fig. 1b, DO-eQTL allelic effects estimated by R/qtl2 ^[21]^). However, several SNPs within a 0.8 Mb sub-region are in high LD, i.e., at a level that greatly exceeds the applicability of existing GWAS fine-mapping methods^[22,23]^, and thus have similarly high LOD scores (Fig. 1c).

**Figure 1:**
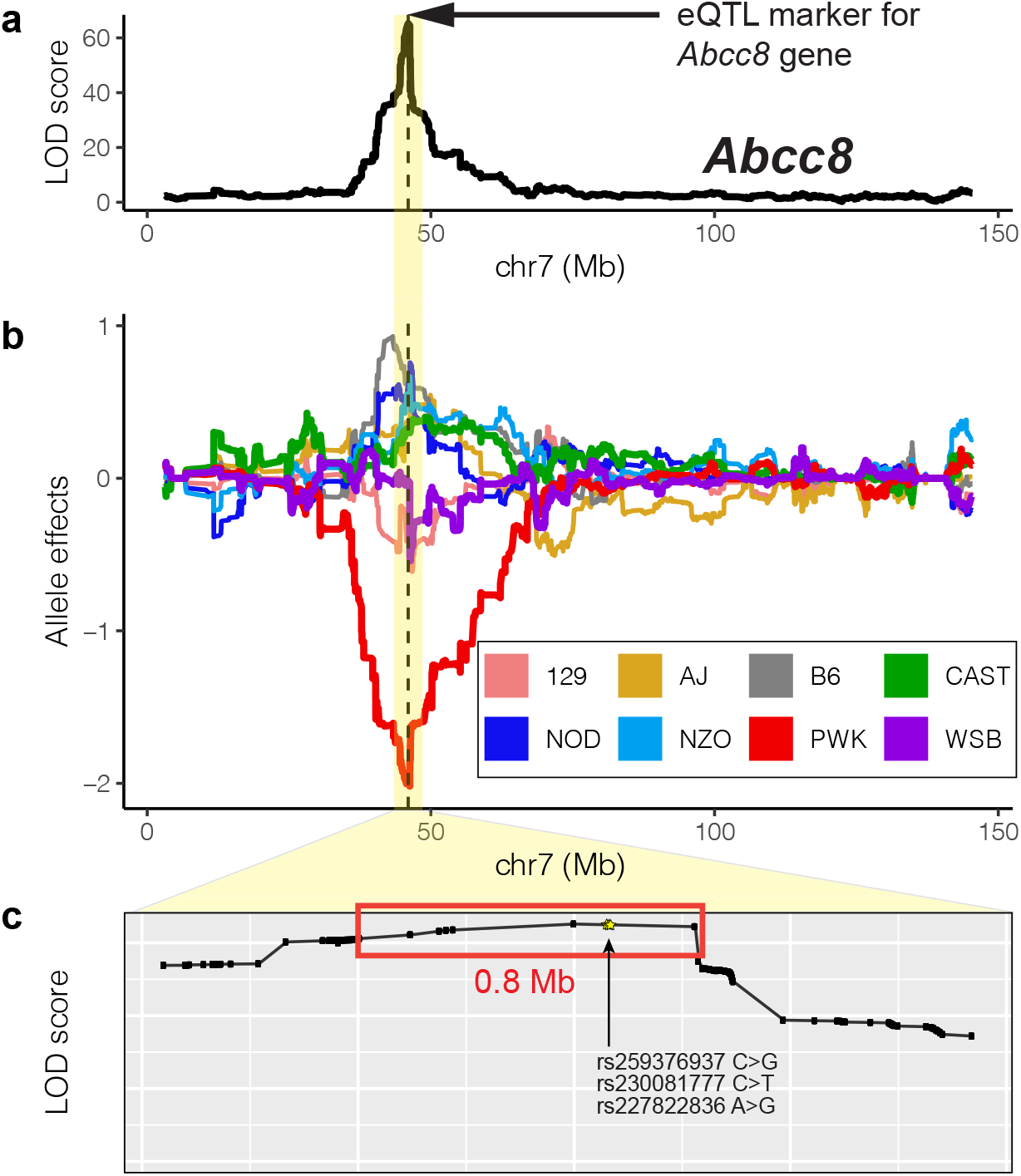
Diversity Outbred (DO) mice expression QTL (eQTL) analysis results at the *Abcc8* locus. **a.** LOD score profile for *Abcc8* is maximized at marker location chr7:46,000,542 (dashed line)^[14]^. **b.** Allele dependence for *Abcc8* local eQTL, where PWK harbors the low expression allele. **c.** Zoomed in version of the LOD score profile from (a) at the SNP level. SNPs tied for the same highest LOD score are marked with the red box. Fine-mapped SNPs by INFIMA are highlighted by yellow stars.

To facilitate fine-mapping of DO-islet eQTLs, we generated functional multi-omics data by assay for transposase-accessible chromatin using sequencing (ATAC-seq) ^[24]^ and transcriptome sequencing (RNA-seq)^[25]^ from the islets of founder DO strains. Analysis of these individual data sets established wide-spread variation in chromatin architecture and gene expression in the DO founder strains. Next, we developed an integrative statistical model named INFIMA (**In**tegrative **Fi**ne-**Ma**pping with Model Organism Multi-omics Data) that leverages multiple multi-omics data modalities to elucidate causal variants underpinning the DO islet eQTLs. INFIMA exploits differences of the candidate genetic variants in terms of their multi-omics data support such as the chromatin accessibility of the variant locations, correlations of chromatin accessibility and transcriptome with variant genotypes and DO mice allelic expression patterns. As a result, it maps genetic variants within the DO founder strains to eQTL genes by quantifying how robustly the multi-omics data explains the allelic patterns observed in the eQTL analysis. Application of INFIMA to islet eQTLs identified in DO mice^[14]^ revealed genetic variants that affect chromatin accessibility, and lead to strain-specific expression differences. Leveraging our INFIMA-based fine-mapping of DO islet eQTLs enabled us to nominate effector genes for ~3.5% of the ~15,000 human GWAS SNPs associated with diabetes. We validated INFIMA fine-mapping predictions with high throughput chromatin capture data from both mouse and human islets. Our results demonstrate that INFIMA provides a foundation for the critical task of capitalizing on model organism multi-omics data to elucidate target susceptibility genes of GWAS risk loci.

## 2 Results

### 2.1 ATAC-seq analysis reveals variable chromatin accessibility in islets of founder DO strains

We performed ATAC-seq to survey chromatin accessibility in pancreatic islets of both sexes of the eight founder DO strains (Fig. 2a; Methods). After quality control with transcription start site (TSS) enrichment analysis (Supplementary: Fig. S1) and data processing, we obtained 77.7 ± 4.1 million reads (excluding mitochondrial DNA) per sample which yielded a total of 51,014 accessible chromatin regions (Supplementary: Fig. S2). Specifically, ATAC-seq reads from 16 samples were aligned to the reference mouse genome (B6) assembly version mm10, yielding an average alignment rate of 92.3 ± 0.7 % (Supplementary: Table S1; Methods). To eliminate potential reference strain bias, we also aligned to individualized genomes, and observed, on average, only 0.86% difference (with a range of 0% and 3.66% across all alignments) between the two alignment strategies (Supplementary: Table S2). Since these differences were not above the level one would expect from slight variation in alignment parameters^[26]^, we used alignments to the reference mouse genome. We identified regions of accessible chromatin with MOSAiCS^[27,28]^ and applied irreproducible discovery rate (IDR) analysis^[29]^ to generate ATAC-seq peak sets of each strain (at IDR of 0.05; Supplementary: Supplementary Notes). The resulting peak sets were then merged to generate a combined peak list. Overall, we observed high concordance of chromatin accessibility (Pearson’s r ~ 0.95) between the sexes for each strain (Supplementary: Fig. S3).

**Figure 2:**
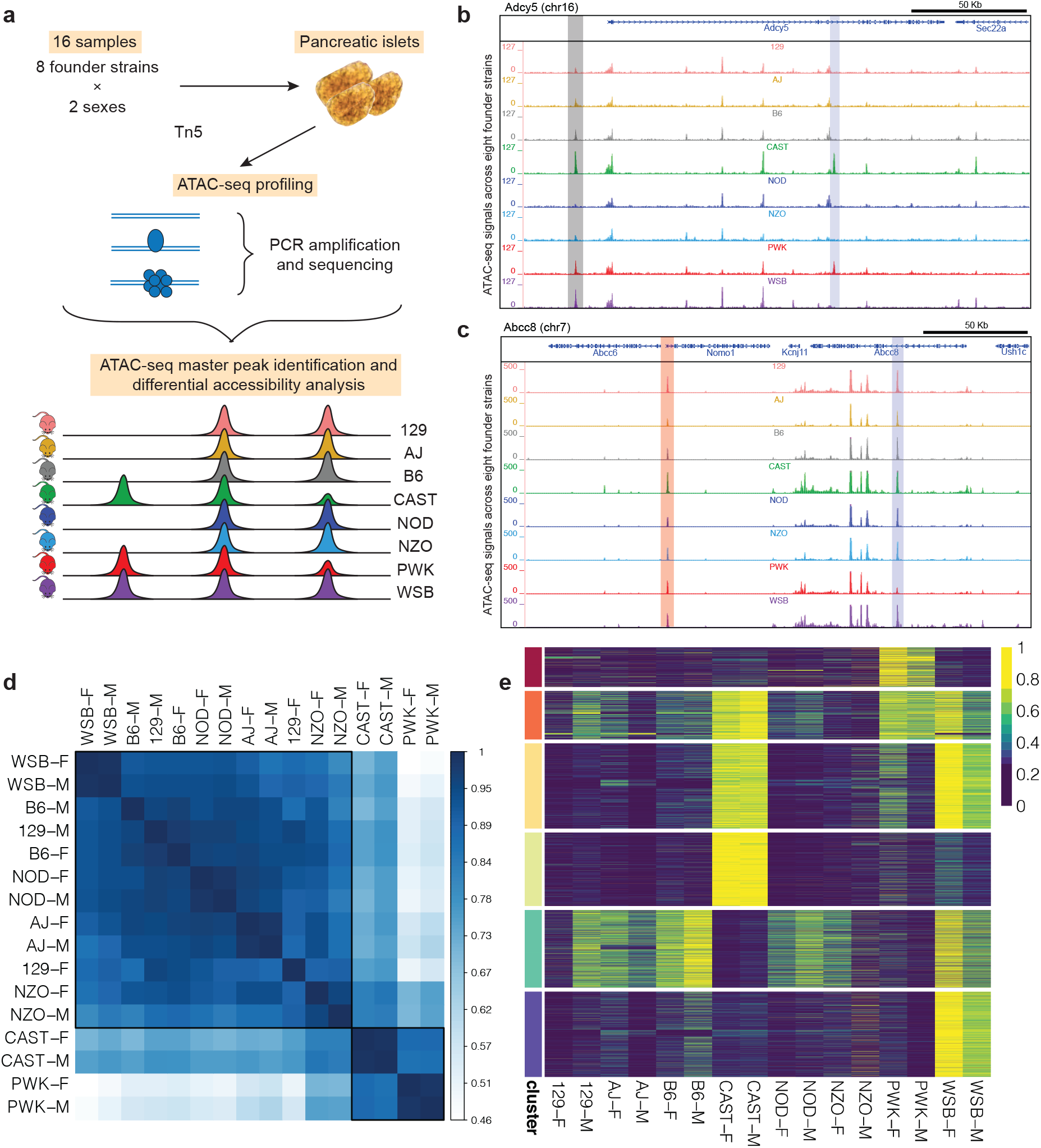
Variable chromatin accessibility across founder DO strains. **a.** Experimental overview and schematic of primary output for chromatin accessibility profiling of founder DO strains by ATAC-seq and differential accessibility analysis. **b - c.** Genome browser displays of differentially accessible ATAC-seq peaks. **b** shows a differentially accessible distal intergenic ATAC-seq peak (translucent gray) and a CAST-PWK specific ATAC-seq peak at the *Adcy5* intron (translucent blue). **c** displays a differentially accessible ATAC-seq peak at the *Nomo1* promoter (translucent red) and an ATAC-seq peak less accessible in PWK at the *Abcc8* intron (translucent gray). **d.** Heatmap of Pearson correlations between each pair of samples based on normalized chromatin accessibility cluster strains consistent with their genetic relatedness. Hierarchical clustering reveals the two clusters of strains outlined in black. **e.** Differentially accessible regions (rows) in 16 samples (columns) of eight founder DO strains across two sexes. ATAC-seq peak scores are standardized to the [0, 1] range. Rows are clustered by k-means (k = 10). The six wild-derived clusters from top to bottom are: PWK, CAST-PWK-WSB, CAST-WSB, CAST, absent in CAST-PWK, WSB. Supplementary: Fig. S2 is the full version of this figure.

More than 70% of the accessible chromatin regions shared by all the strains corresponded to promoters and/or enhancers according to H3K27ac and H3K4me3 ChIP-seq based classification of tissue-specific promoters and enhancers from ENCODE (see URLs; Supplementary: Supplementary Notes). In contrast, only 26.2% of the peaks that were specific to a single strain were annotated as promoters or enhancers (Supplementary: Figs. S8 and S9). These results suggest that most of the strain-specific ATAC-seq peaks occur in strain-specific enhancers that are not captured in the existing list of mouse enhancers from ENCODE.

Among the 51,014 islet ATAC-seq peaks identified, 76.0% showed strain-dependent differences (FDR of 0.05; Methods) in an additive model of strain and sex effect. In contrast, only 50 peaks, 39 of which are located on chromosome X, exhibited sex effects at the same FDR level. The small number of peaks with sex effect is largely driven by the use of strain-specific male and female data to define consistent peaks within a strain and enable irreproducible error rate calculations for robust peak calling. Therefore, our analysis does not reflect the overall chromatin accessibility differences between the sexes of strains. Figs. 2b and 2c display a variety of peaks with strain differences. Specifically, an intronic region of *Adcy5* is more accessible in CAST and PWK compared to other strains, while a distal intergenic region exhibits more accessibility in CAST, PWK, and WSB (Fig. 2b). An intronic region of *Abcc8* is less accessible in PWK compared to other strains, whereas the *Nomo1* promoter is more accessible in CAST (Fig. 2c). We observed that differentially accessible chromatin regions were, overall, over-represented in promoters and under-represented in distal intergenic regions; however, these differentially accessible regions were more likely to be located in distal intergenic regions compared to peaks that did not exhibit significant strain effect (34.5% versus 28.8%, Supplementary: Fig. S10, quantified by regioneR^[30]^ and ChIPseeker^[31]^). Clustering of the normalized ATAC-seq signals of the master peaks across the 16 samples (both sexes, eight strains) revealed a grouping structure largely consistent with the phylogenetic relationships among the founder strains (Fig. 2d). CAST, PWK and WSB are wild-derived subspecies of *M. musculus* ^[32]^, and represent ≥80% of the strain-specific peaks (Fig. 2e). These results suggest that the disproportionate amount of genetic variation contributed by these wild-derived strains mediate much of the differential chromatin accessibility we identified in islets.

Recent computational advances have enabled modeling of the magnitude and the shape of genome-wide chromatin accessibility profiles to infer putative transcription factor (TF) binding sites^[33,34]^. We leveraged PIQ^[33]^ to identify putative TF binding sites within the islet ATAC-seq peaks identified in the founder strains. Utilizing 744 known TF motifs in mouse and human, we identified high-confidence binding profiles for 12 TFs, Mzf1, Gata1, Yy1, Sox10, Nfic1, Ets1, Spib, Znf354c, Gata3, Spi1, Nfatc2, and the complex Arnt:Ahr (Fig. 3a). Nfatc2 is a well-established regulator of *β*-cell proliferation in mouse and human islets^[35]^ and Yy1 ^[36]^, Sox10 ^[37]^, Ets1 ^[38]^, Sbip1 ^[39]^ are TFs abundantly expressed in pancreatic islets. Recent work on a *β*-cell specific knockout of Arnt supports a key role in glucose-stimulated insulin release and islet gene expression^[40,41]^. While the standard footprint analysis considers both the sequence motifs and ATAC-seq signals of binding sites, it cannot discriminate footprints of TFs with similar binding sites. To improve the specificity of the footprint analysis, we integrated the expression levels of TFs in islets from the founder DO strains, with abundant footprints identified from ATAC-seq profiles (Supplementary: Fig. S11), and the sequence similarity between TF motifs (Supplementary: Figs. S12-S16). These additional criteria revealed that the binding motif of the transcriptional repressor Znf354c, which is not expressed in founder islets, is similar to that of Nkx2-2 (Supplementary: Fig. S17), a well-characterized TF that is abundantly expressed and plays a key role in islet development^[42]^. Thus, the Znf354c sites may be occupied by Nkx2-2. In addition, Gata1 and Gata3 are not expressed in founder islets, whereas Gata2, a closely related TF to Gata1 and Gata3 ^[43]^, is highly expressed (Supplementary: Fig. S11), suggesting that it may bind these sites. As expected, *α*-cell specific TFs such as Arx, Irx1, Irx2 showed a fewer number of footprints (≤ 100) within the ATAC-seq peaks than *β*-cell specific TFs^[44]^ (e.g., Pdx1, Mnx1, NFATC2 with an average of ~4,900 footprints). Additional *β*-cell specific TFs, (e.g., Mafk, Pax4, Nkx2-2, Foxa2, Pax6, Nkx6-1), were collectively enriched in ATAC-seq peaks (p-value = 1.66e-2; Supplementary: Supplementary Notes), albeit with fewer footprints (~1,900).

**Figure 3:**
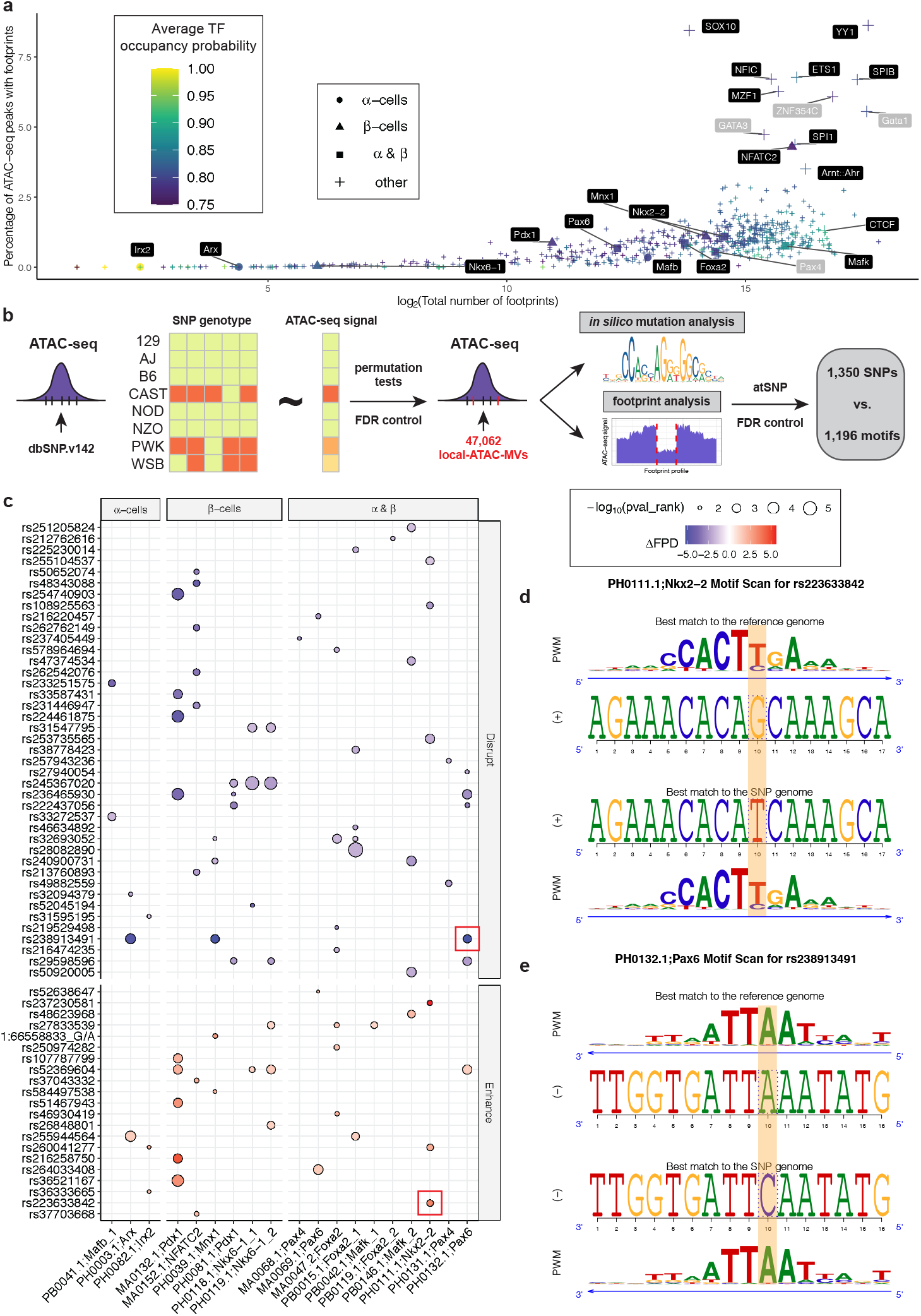
Genetic variants associate with differential chromatin accessibility in the islets of DO founder strains. **a.** Footprint analysis of ATAC-seq peaks. Transcription factors in black labels are expressed in founder islets (Supplementary: Fig. S11). **b.** Overview of local-ATAC-MV identification, footprint, and *in silico* mutation analysis with atSNP. **c.** A subset of SNPs (rows) that enhance/disrupt islet specific footprints (columns). The circles depict changes in the footprint depth with the SNP and reference alleles (ΔFPD = FPD_*SNP*_ − FPD_*REF*_). Enhancement and disruption based on comparative FPD are depicted by shades of red and blue, respectively. The circle size indicates the significance of the impacts of the SNP alleles to the motif matches as calculated by atSNP^[93]^. Larger circles correspond to more significant changes in the motif match. Examples in **d, e** are highlighted by red boxes in panel **c**. **d.** atSNP composite logo plot depicting Nkx2-2 binding site enhancement by SNP rs223633842 (G → T). **e.** atSNP composite logo plot depicting Pax6 binding site disruption by SNP rs238913491 (A → C).

### 2.2 Genetic variants associate with differential chromatin accessibility in islets of founder DO strains

We next evaluated the contribution of genetic variability present in the eight founder DO strains to differential chromatin accessibility within their islets. We associated the signal of 22,200 ATAC-seq peaks with at least one SNP, with the genotypes of the SNPs that they harbor. Although chromatin accessibility of a genomic region demarcated by an ATAC-seq peak can be modulated by SNPs in proximal and distal ATAC-seq peaks or genomic regions, we considered only the local SNPs to alleviate the multiple testing problem.

As a result, we identified 47,062 local ATAC-seq signal modulating variants (local-ATAC-MVs) within these 22,200 ATAC-seq peaks at FDR of 0.05 (Fig. 3b; Methods). The distribution of the number of local-ATAC-MVs within ATAC-seq peaks is right-skewed (Supplementary: Fig. S18) indicating that most peaks have one to three local-ATAC-MVs. Overall, 16,549 (42.7%) of the 38,749 differential peaks do not harbor any local-ATAC-MVs, suggesting that SNPs, or other factors, outside the ATAC-seq peaks contribute to their variable accessibility among the strains. The vast majority (95.6%) of the local-ATAC-MVs are associated with SNPs present in the three wild-derived strains (Supplementary: Fig. S19). Furthermore, a large percentage of the local-ATAC-MVs (77.3%) reside in distal intergenic or intronic regions, while 18.7% occur within promoters (Supplementary: Fig. S20).

Genetic variants can affect gene regulation by changing TF binding affinities to genomic sequences^[45]^. To assess whether local-ATAC-MVs influence TF binding, we first performed an *in silico* mutation analysis of TF binding using atSNP^[46]^. In addition, for each SNP-motif pair, we computed the relative change in footprint depth (FPD), a measure of TF activity within ATAC-seq peaks^[47]^, at the motif location across strains with the reference and alternative alleles (Supplementary: Fig. S21). Overall, we identified 8,029 loci where local-ATAC-MVs significantly influenced the footprint at TF binding sites after multiplicity adjustment at FDR level of 0.05 (see Fig. 3b for the overall pipeline and Supplementary: Fig. S22 and S23 for evaluation of all the SNP-motif combinations; Methods). Despite the stringent multiplicity adjustment, we identified 62 local-ATAC-MVs that impact binding sites of TFs that are highly expressed in *α*, *β* or other islet cell types^[44]^ (Fig. 3c). For example, the SNP rs223633842 enhances a Nkx2-2 motif (Figs. 3d), whereas the SNP rs238913491 disrupts a Pax6 motif (Figs. 3e). Together, these results suggest that strain-specific differences in chromatin accessibility are affected by local-ATAC-MVs residing within ATAC-seq peaks and distrupting or enhancing TF binding.

### 2.3 RNA-seq analysis in islets of founder DO strains reveals variable transcriptome

After establishing widespread association of SNP genotypes with differential chromatin accessibility in the founder DO strains, we sequenced the islet transcriptome of the same eight strains. This enabled us to link local-ATAC-MVs with strain-dependent differences of nearby gene expression. We quantified the expression of 13,568 protein-coding genes with RSEM^[48]^ (Supplementary: Fig. S24; Methods) which appropriately clustered the samples based on strain (Fig. 4a, Supplementary: Fig. S25). To maximize statistical power, we associated only the founder local-ATAC-MVs, instead of all the founder SNPs, with gene expression and identified 34,711 (73.8%) local-ATAC-MVs as associating with *cis* (as defined by 1 Mb neighbourhood of genes) gene expression variation (Methods). The expression patterns of the genes associated with the local-ATAC-MVs are largely driven by alleles of wild-derived strains CAST, PWK, and WSB. Specifically, alleles of these three strains exert the most significant associations of the genes, i.e., the top 6 genotypes driven by these strains compromise 50.3% of the top associations of the 6,418 local-ATAC-MV-associated genes (Fig. 4b). Next, we evaluated the distance between these genes and the proximal associated local-ATAC-MV loci. We found widespread contribution of promoters to expression variation across strains by harboring associated local-ATAC-MVs, i.e., 58% of the genes with at least one local-ATAC-MV association had associated local-ATAC-MV loci in their promoters (Supplementary: Fig. S26). We further investigated how well the differential ATAC-seq peaks within promoters explained the variation in gene expression across the strains. A pairwise differential expression analysis (Methods; FDR of 0.05) for the eight founder strains identified eGenes that were selective for one strain, i.e., B6 eGenes (expressed more in B6) and CAST eGenes (expressed more in CAST). As expected, B6 eGenes have higher promoter accessibility in B6, whereas CAST eGenes have higher promoter accessibility in CAST (Supplementary: Fig. S27). This concordance between strain-selective promoter accessibility and gene expression was observed, on average, for 67% of the eGenes (Supplementary: Fig. S28), suggesting a strong contribution of genetic variance of chromatin architecture within promoters to proximal gene regulation as also observed by others^[49–52]^.

**Figure 4:**
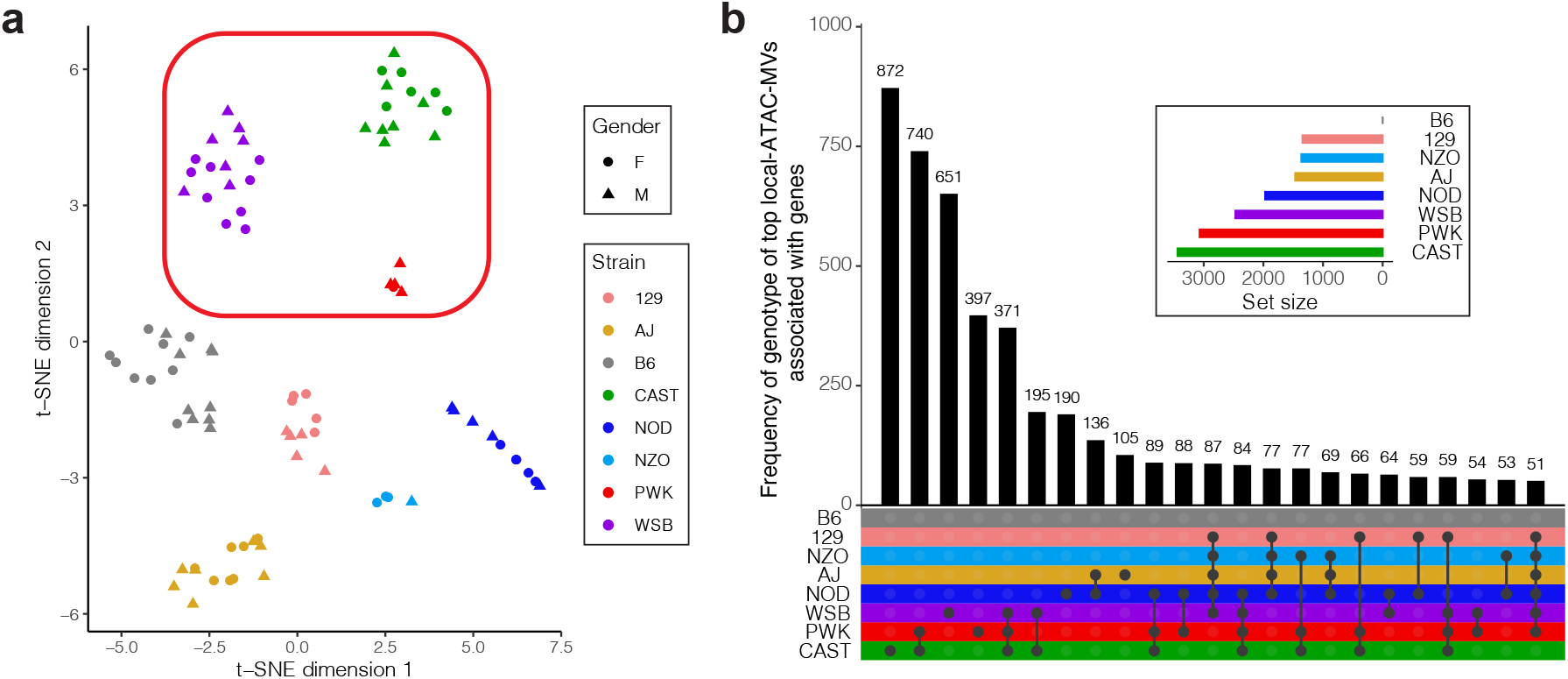
Variable transcriptome across islets of founder DO strains. **a.** Two-dimensional projection of the 91 founder RNA-seq samples with tSNE. Samples from wild-derived strains are boxed in with the red rectangle. **b.** UpSet plot^[102]^ for the frequencies of genotypes of local-ATAC-MVs associated with founder islet gene expression. Each gene with at least one significant association contributed its most significant local-ATAC-MV. Genotypes with frequencies less than 50 are not displayed.

### 2.4 INFIMA model for fine-mapping DO mouse islet eQTLs by leveraging founder strain islet ATAC-seq and RNA-seq

The strain-dependent differences in accessible chromatin and transcriptome landscapes in islets of the DO founder strains allowed us to identify local-ATAC-MVs and their putative effector genes. Next, we leveraged this founder data to fine-map islet eQTLs from DO mice^[14]^ (DO-eQTL, Figure 1). We developed an integrative framework, named INFIMA, that exploits the high-resolution of the founder ATAC-seq profiles and gene expression data to delineate enhancer-sized loci as the most likely causal locus for individual DO-eQTLs.

INFIMA is an empirical Bayes model that estimates the linkages between founder local-ATAC-MVs and DO-eQTL genes for improving the resolution of DO-eQTL analysis. This is achieved by quantifying how well each non-coding SNP in high LD with the islet DO-eQTL marker explains the observed relationship between the allelic effect of the eQTL, islet ATAC-seq profile and gene expression among the founder strains proximal to the marker locus, and derived TF footprint results (Fig. 5a). This quantification enables inferring the likelihood of each candidate SNP, implied by the marker, to be causal. We summarize the INFIMA framework in Fig. 5 and provide the statistical details in this section.

**Figure 5:**
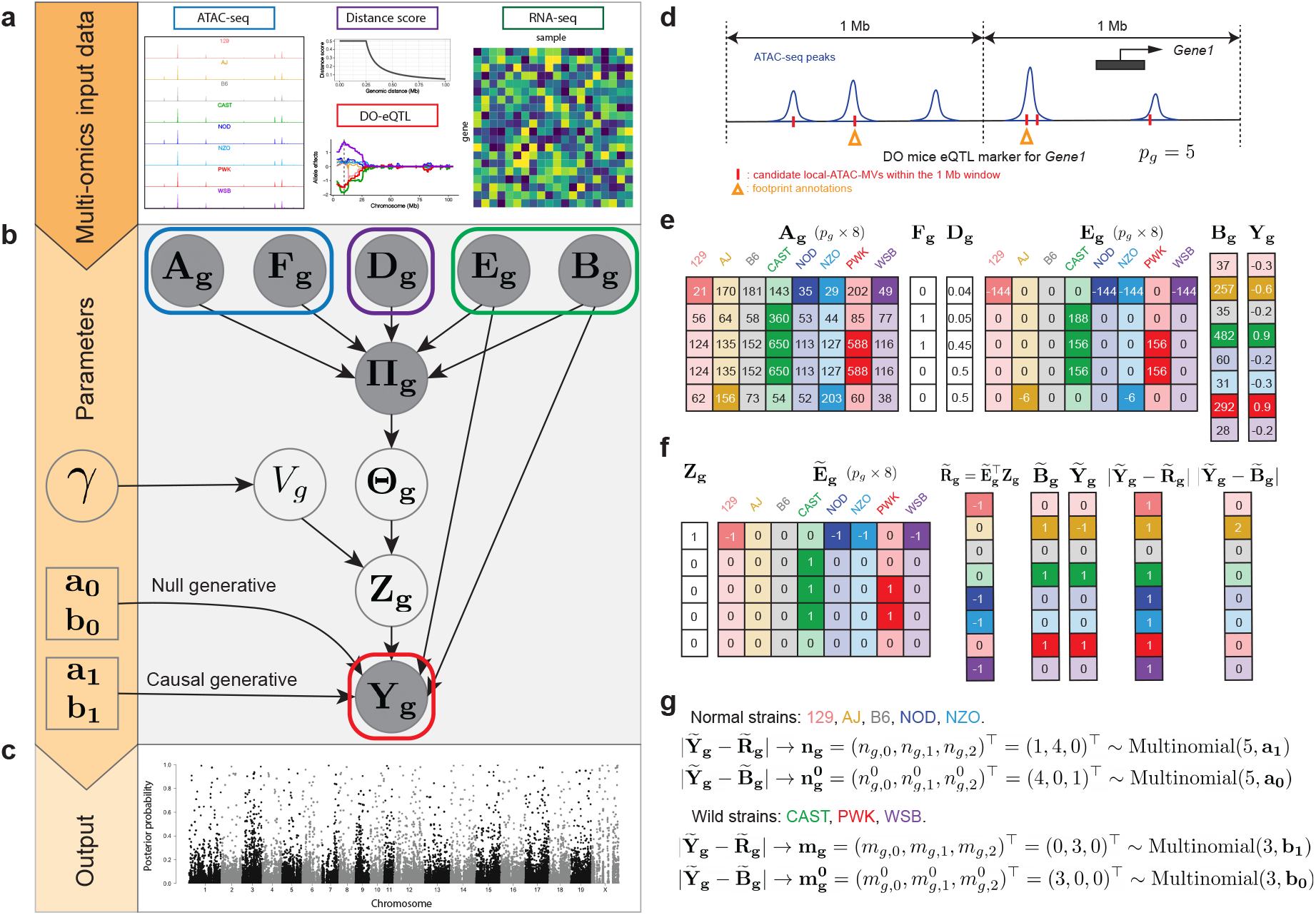
INFIMA model overview. **a.** Input data for the INFIMA model. INFIMA leverages summaries of model-organism multi-omics data to model the relationship between allelic expression patterns (**Y**_**g**_) of DO-eQTL genes (i) and founder expression patterns (**B**_**g**_) under a null model of no causal SNPs (*V*_*g*_ = 0); (ii) and founder genotype expression patterns **R**_**g**_ = **E**_**g**_^⊤^**Z**_**g**_, where **E**_**g**_ represents genotype effects of candidate SNPs on founder expression and **Z**_**g**_ encodes the causal SNP for gene *g*, under an alternative model with causal SNPs (*V*_*g*_ = 1), across all the genes indexed by *g*. **b.** Plate representation of the INFIMA model summarizing data and the parameters. Blank circles: latent variables and parameters to be inferred; Filled circles: observed variables. **c.** INFIMA infers SNP-level posterior probabilities of association for fine-mapping across all the candidate local-ATAC-MVs. **d-g.** An example input of the INFIMA model. **d.** An overview of a *W* = 1 Mb window around a DO-eQTL marker (centered dashed line) associated with *Gene1*. Two out of five candidate local-ATAC-MVs (red short lines) are decorated with comparative footprint effects (orange triangles). **e.** Example input data for the five candidate local-ATAC-MVs. **f.** An illustration of data trinarization and edit distance with multinomial distributions. The trinarization details can be found in Methods. **g.** The edit distance variables quantify how many strains have 0, 1, or 2 absolute distances between 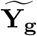 and 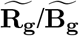 and are modeled by multinomial distributions.

A key step in the INFIMA framework is featurization of the DO-eQTL and founder data. We let 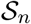 and 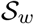 denote the index set for the classical in-bred (129, AJ, B6, NOD, NZO) and wild-derived strains (CAST, PWK, WSB), respectively, and let *s* be the index for the strains. Let *G* denote the total number of instances of DO-eQTL data, i.e., total number of gene-marker associations, *g* = 1 *... G* the index for the DO-eQTL gene of the *g*^*th*^ instance, *p*_*g*_ the number of candidate local-ATAC-MVs within a window size *W* of the eQTL marker for gene *g*, and *k* = 1 *... p*_*g*_ the index for the local-ATAC-MVs within this window (Fig. 5d). In our application, we have *G* = 10,936 contributed by 8,046 eQTL markers versus 10,393 genes. A given DO-eQTL marker can have multiple DO genes that it associates with (Supplementary: Fig. S29). Let **Y**_**g**_ be an 8 × 1 vector of DO-eQTL allelic expression effects estimated with R/qtl2 ^[21]^ at marker location with the highest LOD score. We denote the features extracted from founder ATAC-seq and RNA-seq by **X**_**g**_ = (**A**_**g**_, **F**_**g**_, **D**_**g**_, **E**_**g**_, **B**_**g**_), where **A**_**g**_ is a *p*_*g*_ × 8 matrix of the normalized ATAC-seq signal of the peak each candidate local-ATAC-MV resides in; **F**_**g**_ is the indicator vector (*p*_*g*_ × 1) of whether or not the candidate local-ATAC-MV is affecting a footprint significantly, i.e., it is among the set of 8,029 SNP-motif combinations identified in the aforementioned comparative footprint analysis; **D**_**g**_ is a *p*_*g*_×1 vector of distance scores computed from the distances of local-ATAC-MV to the promoter of gene *g*; **E**_**g**_ is a *p*_*g*_×8 matrix of founder RNA-seq genotype effects of these candidate SNPs for gene *g* (i.e., marginal regression of gene expression with respect to genotype); **B**_**g**_ denotes an 8 × 1 vector of the normalized founder expression of gene *g*. Fig. 5e illustrates an example of the extracted features.

INFIMA model assumes at most one causal local-ATAC-MV per gene for a single marker-gene association. This is encoded by an unobserved random variable *V*_*g*_ ∈ {0, 1} representing the number of causal local-ATAC-MVs for eQTL gene *g*. While this assumption can be relaxed at the expense of computational cost, it already enables multiple causal loci per gene when the gene is associated with multiple markers. Next, we define an additional unobserved *p*_*g*_ × 1 random variable 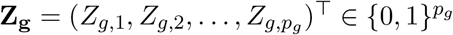 to denote the causal local-ATAC-MV. It immediately follows that **1**^⊤^**Z**_**g**_ = *V*_*g*_. Finally, in the presence of a local-ATAC-MV, i.e., *V*_*g*_ = 1, we define **R**_**g**_ = **E**_**g**_^⊤^**Z**_**g**_ as an 8 × 1 vector of the genotype effects of the causal SNP estimated from founder RNA-seq data for gene *g*.

For causal SNPs, we expect the allelic effects from DO mice (**Y**_**g**_ from the eQTL study) to be in agreement with the genotype effect of the causal SNP on the founder expression (**R**_**g**_). We quantify this relationship with a causal generative model of **Y**_**g**_ conditional on **R**_**g**_. To avoid parametric assumptions needed for modeling continuous allelic effects **Y**_**g**_ and **R**_**g**_, in addition to supporting potential differences in distributions for the classical in-bred and wild-derived strains, we consider an edit distance model. Specifically, we convert **Y**_**g**_, **R**_**g**_, and **B**_**g**_ to trinary indicators encoding three levels of signal strengths: lower, the same, and higher than the reference strain B6 (Fig. 5f; Methods). After trinarizing the effects 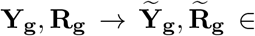 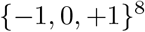, we compute absolute values of the differences between their trinarized values 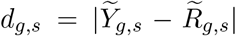 for each strain *s*. Then, we define the edit distance random variables 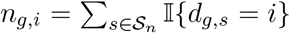 and 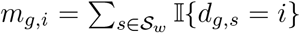 for *i* = 0, 1, 2. The set of edit distances (*n*_*g*, 0_, *n*_*g*, 1_, *n*_*g*, 2_) represent numbers of 0’s, 1’s, and 2’s in an experiment that corresponds to rolling a 3-sided dice 5 times. Hence, it follows that **n**_**g**_ = (*n*_*g*,0_, *n*_*g*,1_, *n*_*g*,2_)^⊤^ ~ Multinomial(5, **a**_**1**_) and, similarly, **m**_**g**_ = (*m*_*g*,0_, *m*_*g*,1_, *m*_*g*,2_)^⊤^ ~ Multinomial(3, **b**_**1**_). Here, **n**_**g**_ = (5, 0, 0) and **m**_**g**_ = (3, 0, 0) indicate that the allelic expression pattern in the DO mice completely matches the genotype effect estimated from the founders for gene *g* and the causal SNP specified by **Z**_**g**_. In this model, the lack of a candidate causal SNP is encoded by *V*_*g*_ = 0. However, some concordance between DO mice allelic expression **Y**_**g**_ and founder gene expression **B**_**g**_ is still warranted. Leveraging this intuition, we develop a null generative model for **Y**_**g**_ conditional on **B**_**g**_ with a similar trinarization approach as above. The trinarized data 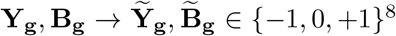, with absolute differences 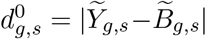 can be defined similarly as in *V*_*g*_ = 1. We define edit distance random variables 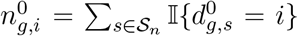 and 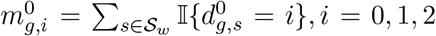 and assume individual multinomial distributions 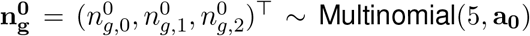, 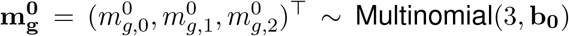, parametrized by parameters **a**_**0**_ and **b**_**0**_, respectively. Fig. 5g illustrates an example of the trinarized data and the corresponding edit distances.

Next, we combine the two settings, namely *V*_*g*_ = 1 and *V*_*g*_ = 0, as a mixture over the two generative models. Specifically, we assume that the latent causal indicators are random draws, i.e., 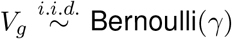, with the prior probability, *γ* ∈ (0, 1), for the causal generative model. Let 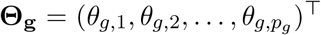 denote the probabilities that each candidate SNP is causal for gene *g*; then, **Z**_**g**_ is a mixture distribution over a Multinomial distribution and a point mass at vector of 0’s as

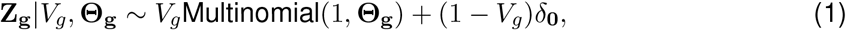

where *δ*_**0**_ is a size *p*_*g*_ vector of 0’s. To leverage the multiomic data further, we assume a Dirichlet prior for the probability vector **Θ**_**g**_|**Π**_**g**_ ~ Dirichlet(**Π**_**g**_), where 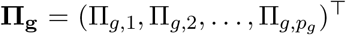 is defined as

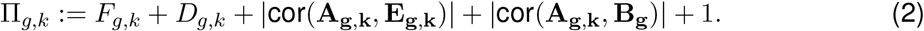

Here, each component of Π_*g,k*_ provides prior multi-omics information that contributes to the likelihood of SNP *k* to be causal for gene *g*. Specifically, *F*_*g,k*_ ∈ {0, 1} indicates impact on a TF binding site; *D*_*g,k*_ ∈ (0, 0.5] is a function of the distance between the DO-eQTL marker and the candidate SNP to utilize genomic distance; |cor(**A**_**g,k**_, **E**_**g,k**_)| ∈ [0, 1] measures the correlation between ATAC-seq signal of the peak harboring SNP *k* and the genotype effect of SNP *k* on founder expression; |cor(**A**_**g,k**_, **B**_**g**_)| ∈ [0, 1] similarly quantifies the correlation between ATAC-seq signal and gene expression in the founder strains.

The combined generative model for DO-eQTL effect size **Y**_**g**_ is then given by

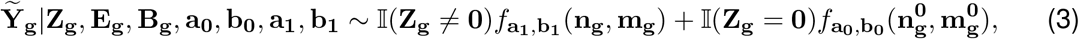

where 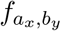 denotes the product of Multinomial probability distribution functions parametrized by *a*_*x*_ and *b*_*y*_. In summary, INFIMA model takes as input DO-eQTL results, summarized functional data from RNA-seq and ATAC-seq analysis of founder strains, as well as ATAC-seq-based comparative footprint and *in silico* mutation analysis of SNPs and outputs SNP-level quantifications (Fig. 5c).

### 2.5 Simulations reveal improved statistical power and fine-mapping with INFIMA

We first evaluated INFIMA for its ability to improve statistical power of fine-mapping and identification of credible sets of SNPs in marker eQTL applications. We designed data-driven simulations where the parameters of the generative model are set based on the actual DO-eQTL and summarized founder strain multi-omics data from ATAC-seq, RNA-seq, and comparative footprint and *in silico* mutation analysis. We varied the prior information extracted from the multi-omics data to be non-informative (NI), moderately informative (MI), and highly informative (HI) by varying the information contributed by the comparative footprint analysis (Methods).

This allowed modulation of the informativeness of the prior parameters without considering generative models for summaries extracted from ATAC-seq and RNA-seq data. INFIMA model has two key inference variables: *V*_*g*_ ∈ {0, 1} which encodes whether or not a gene has a causal SNP, and 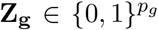 which encodes the causal SNP. Although the prior parameter *γ* for *V*_*g*_ does not depend on the summarized multi-omics data (i.e., is expected to be insensitive to the prior information), varying levels of informativeness in the multi-omics data yield improved area under receiving operating characteristics and precision recall curves, with an average of 0.61 ± 0.079% improvement in power from moderately to highly informative setting (Supplementary: Fig. S30). Since INFIMA leverages the multi-omics data to specifically infer **Z**_**g**_ by informing the prior probabilities of causal SNPs, we assessed the impact of levels of informativeness of the priors on fine-mapping. Specifically, we considered the most and least likely causal associations inferred by INFIMA for each gene as “Most Likely”: local-ATAC-MV with the highest posterior probability of being causal; “Least Likely”: local-ATAC-MV with the lowest posterior probability of being causal. We compared these INFIMA strategies with three intuitive and model-free baseline strategies of selecting causal SNPs as “Random”: a randomly selected local-ATAC-MV; “Closest to Marker”: local-ATAC-MV closest to the DO-eQTL marker in genomic distance; “Closest to Gene”: local-ATAC-MV closest to the gene promoter in genomic distance. This comparison revealed that INFIMA predictions provide markedly better fine-mapping compared to baseline strategies regardless of the level of informativeness of the priors. Specifically, the “Most Likely” selection by INFIMA provided the smallest credible proportion (the minimum proportion of ranked candidate local-ATAC-MVs required to encompass the causal variant). The NI, MI, and HI settings yielded 33.90%, 22.22%, and 14.04% credible proportions, respectively (Fig. 6), compared to the minimum of 52.48%, 48.32% and 50.00% achievable with the baseline strategies. Interestingly, even when the priors are non-informative (NI setting), the INFIMA-produced credible set is, on average, 29.1% smaller than the smallest set that can be achieved by the baseline strategies (33.90% by NI vs. 48.32% by MI). As expected, the least likely predictions with INFIMA performed worse than baseline strategies, confirming INFIMA’s ability to rank local-ATAC-MVs with respect to their causal potential. Overall, these simulations highlighted the significance of integrating multi-omics data into fine-mapping.

**Figure 6:**
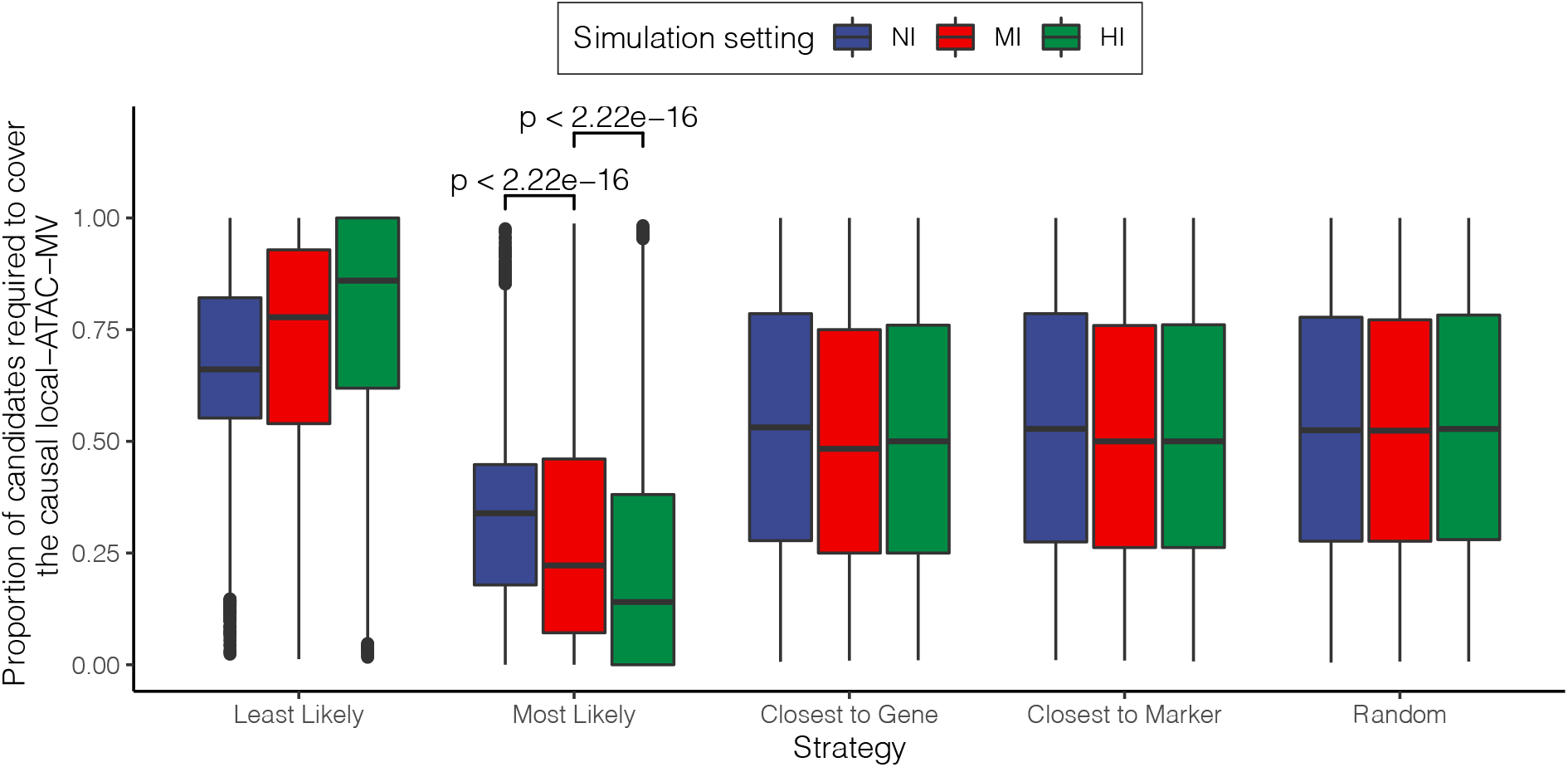
Simulations reveal improved statistical power and fine-mapping with INFIMA. Comparison of the fine-mapping performances of five strategies across three simulation settings: NI: Non-informative; MI: Moderately informative; HI: Highly informative. The y-axis reports, as the performance metric, the proportion of candidate local-ATAC-MVs required in the credible set to cover the causal SNPs.

### 2.6 INFIMA outperforms alternatives for fine-mapping DO mouse eQTLs

We fit INFIMA model with a 1 Mb window size (*W*) around DO-eQTL markers across all the *G*=10,936 gene-marker associations (8,046 eQTL markers and 10,393 genes). This resulted in a right-skewed distribution for the number of candidate local-ATAC-MVs within a window (Fig. 7a, median = 36.0, sd = 26.9). Fig. 7b summarizes the estimated posterior probabilities of having a causal local-ATAC-MV, i.e., 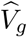, across the genes. It indicates that INFIMA infers a causal local-ATAC-MV for 3,846 (38.0%) DO-eQTL genes at FDR of 0.05.

**Figure 7:**
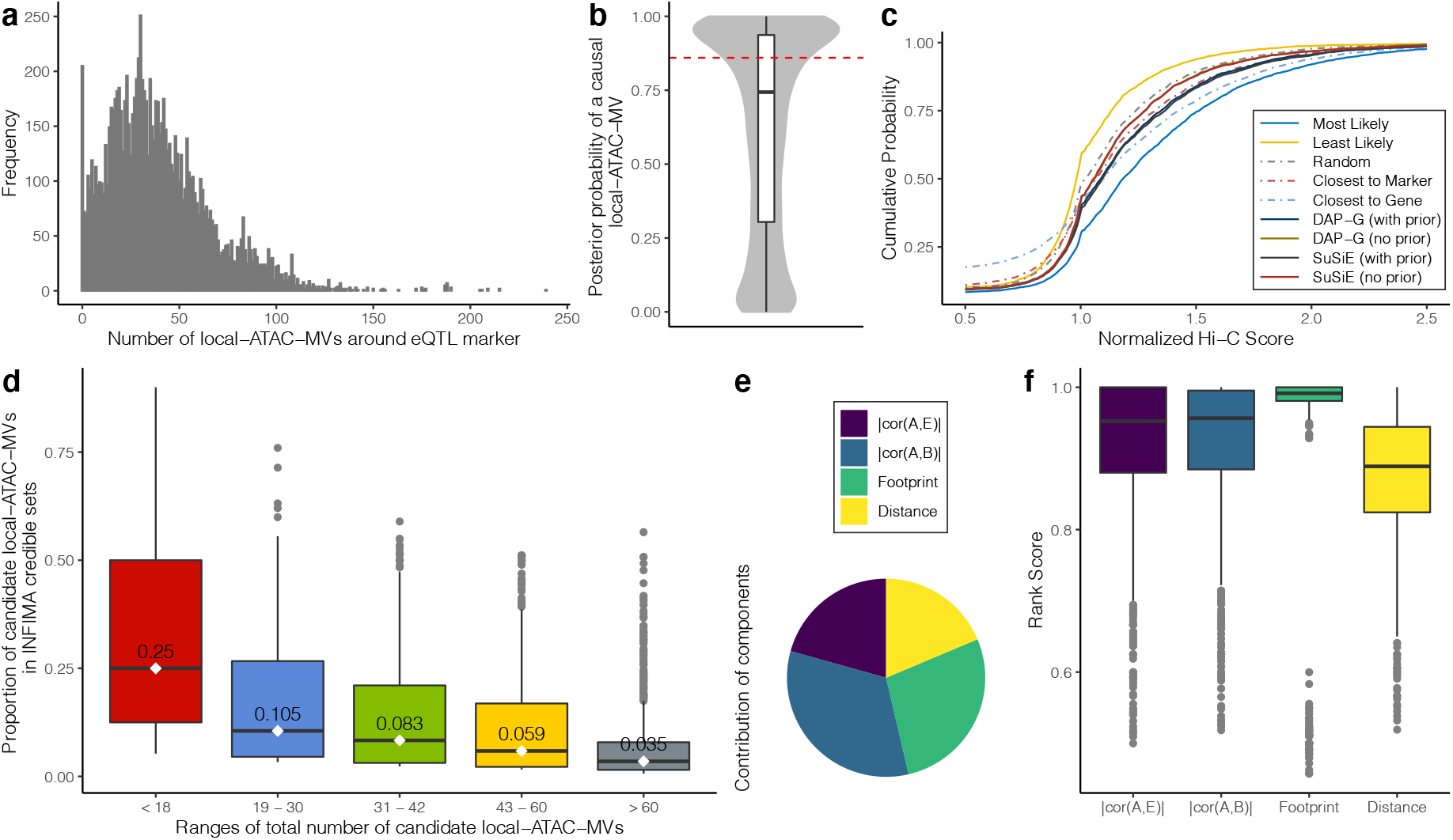
INFIMA outperforms alternatives for fine-mapping DO mouse eQTLs. **a.** Histogram of numbers of local-ATAC-MVs around DO-eQTL markers with window size of *W* = 1 Mb. **b.** Boxplot of INFIMA posterior probabilities of association, 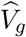, across all genes. Red dashed line depicts the posterior probability cutoff for FDR of 0.05. **c.** Evaluation of fine-mapping strategies with empirical cumulative distribution of normalized easy Hi-C scores. “Most Likely” and “Least Likely” refer to most and least likely predictions from INFIMA, respectively. **d.** Boxplots depict the proportion of the candidate causal local-ATAC-MVs that are included in the credible set by INFIMA stratified by the size of candidate sets. The intervals on x-axis are from quantile bins (20%, 40%, 60%, 80%, 100% percentiles) of number of local-ATAC-MVs around the eQTL marker, *p*_*g*_. Median values are displayed on each boxplot. **e.** Proportion of times each of the individual multi-omic components are the leading contributors to the INFIMA prior probability of causality: Correlation between ATAC-seq signal and founder eQTL effect sizes |*cor*(*A, E*)| : 0.207; Correlation between ATAC-seq signal and founder gene expression |*cor*(*A, B*)| : 0.330; Footprint: 0.277; Distance: 0.186. **f.** The rank scores of the inferred causal local-ATAC-MVs when individual components are the top ranking contributors. The higher the rank scores are, the more INFIMA weights in the component when inferring causal local-ATAC-MVs.

We further summarized INFIMA results as we have done for the simulations by identifying the most likely and least likely causal local-ATAC-MVs for genes with an inferred causal SNP, and compared these with the baseline strategies outlined in the simulations. In addition to these baseline methods, we also considered two recent human GWAS fine-mapping methods DAP-G^[53,54]^ and SuSiE^[55]^, both of which have demonstrated best performances in human GWAS fine-mapping studies. We initially considered applying DAP-G and SuSiE to all the SNPs tagged by the eQTL marker at the individual locus without restricting the set of SNPs to local-ATAC-MVs and by utilizing the multi-omics prior on the full set of SNPs. However, both methods failed to generate credible sets under this setting (Supplementary: Supplementary Notes) owing to the LD structure of the DO mice (Supplementary: Fig. S31). Therefore, we reduced the candidate SNP set to local-ATAC-MVs for fine mapping with DAP-G and SuSiE. We leveraged high resolution easy Hi-C data, processed with a recent computational pipeline^[56]^, from mouse islets and computed the empirical cumulative distribution curve of Hi-C signal between the DO-eQTL genes and their selected local-ATAC-MVs. We expect the local-ATAC-MVs that are likely to be true positives to interact with the gene promoters and, as a result, to exhibit high Hi-C signal compared to competing approaches. Fig. 7c depicts that the “Most Likely” selection by INFIMA outperforms the baseline predictions while the “Least Likely” selection by INFIMA performs worse than the baselines, highlighting an overall goodness-of-fit by INFIMA. The cumulative distribution curve of the “Most Likely” selection is significantly distinct from the baseline strategies (quantified by three different metrics: Kolmogorov-Smirov test, Kullback-Leibler (KL) divergence, and Chi-Squared test, Supplementary: Table S4-S6, Addition file 1: Fig. S32), confirming that INFIMA prediction of local-ATAC-MVs for DO-eQTL genes tend to be supported by higher Hi-C interaction signals. While the performances of DAP-G and SuSiE improve markedly with the INFIMA multi-omics data prior, they still perform worse than the baseline “Closest to Gene” and are significantly inferior to INFIMA. This is likely attributable to the large numbers of local-ATAC-MVs that are in perfect LD in DO mice compared to typical human GWAS fine mapping studies (Supplementary: Figs. S33). Hi-C contacts standardized to [0, 1] for each DO-eQTL gene to enable comparison across genes indicate that, concordant with the overall Hi-C score distribution comparison, the “Most Likely” and the “Least Likely” selections by INFIMA harbor the highest and lowest ranked Hi-C scores, respectively (Supplementary: Fig. S34).

After validating that INFIMA inferred causal local-ATAC-MVs are significantly better than those identified by the baseline and alternative strategies, we evaluated the impact on fine-mapping. INFIMA is able to reduce the size of the credible set of local-ATAC-MVs tagged by a marker by 96.5% when *p*_*g*_ > 60. When the set size, *p*_*g*_, is ≤ 18 (the lowest 20%), INFIMA reduces the size of the set of candidate local-ATAC-MVs by 75.0% (Fig. 7d). These are significant reductions at both the high and low ends of the size of the tagged local-ATAC-MV sets of a marker as it markedly reduces the number of loci for follow-up.

Since the multi-omics data INFIMA leverages to inform SNP prior probability of causality is multi-component, we asked whether the individual components contributed differently to the learned priors, i.e., **Π**_**g**_. Specifically, for each causal local-ATAC-MV of gene *g*, we ranked each of the individual components across the same category of components from all the competing *p*_*g*_ local-ATAC-MVs in ascending order, calculated a rank score^1^ by normalizing with *p*_*g*_, and reported the highest ranking contributor for the causal local-ATAC-MV as the component with the highest rank score. We found that, for only 20.1% of the causal local-ATAC-MVs, the Distance is the highest ranking contributor to the prior. The correlation between ATAC-seq signal and gene expression, i.e., |cor(*A, B*)|, contributes the most at 33.0% (Fig. 7e). Fig. 7f shows that when Distance is the leading contributor, the median rank scores of the causal local-ATAC-MV, at 0.889, is lower than other components. This further demonstrates that INFIMA is not biased towards the local-ATAC-MVs closest to the genes. Interestingly, the Footprint component, with the highest median rank score of 0.992 (Fig. 7f), exerts a salient impact on INFIMA’s ability to discriminate among the set of candidate causal local-ATAC-MVs.

### 2.7 INFIMA generates candidate susceptibility genes for human GWAS SNPs

The INFIMA model links ATAC-seq peaks and local-ATAC-MVs to candidate effector genes by fine-mapping DO-eQTLs. Next, we asked whether this approach can be leveraged to assign putative target genes in islets for non-coding human GWAS SNPs associated with diabetes.

Specifically, we considered 14,434 SNPs associated with 16 diabetes-related physiological traits from human GWAS^[57]^ (Supplementary: Fig. S35). We employed a two-step peak-based strategy to lift-over human GWAS SNPs to syntenic sequences in the mouse genome. We first lifted-over the GWAS SNPs directly using the UCSC lift-over tool (see URLs) and identified the nearest mouse ATAC-seq peak to the syntenic loci. The remaining GWAS SNPs (81.0%) that did not directly lift-over to the mouse genome were first linked to their nearest human islet ATAC-seq peaks^[58]^ and the peaks were lifted-over to mouse and linked to the nearest mouse islet ATAC-seq peak within 10 Kb (Supplementary: Fig. S36; Methods). This resulted in syntenic links between 4,268 GWAS SNPs (2,749 direct and 1,519 ATAC-seq peak-based) and 1,532 mouse ATAC-seq peaks. Several studies^[59–61]^ have proposed that genomic compartment annotations associated with promoters are largely conserved between human and mouse. Similarly, distal regulatory elements across species are more likely to reside in regions with similar genomic compartment annotations^[62,63]^. Therefore, we asked if these diabetes-associated syntenic regions had common genomic compartment annotations with their human counterparts. Overall, we observed a large degree of genomic annotation conservation for diabetes-associated GWAS SNPs (Fig. 8a). Specifically, ~70% of the local-ATAC-MVs syntenic to intronic/distal/promoter GWAS SNPs exhibited the same genomic compartment annotation in mouse. Furthermore, we found that mouse syntenic regions of GWAS SNPs associated with diabetes-linked traits, e.g., Type 1 Diabetes, Type 2 Diabetes, Body Mass Index, and Body Weight were enriched for local-ATAC-MVs (Fig. 8b; Bonferroni of 0.05, Methods). In contrast, mouse syntenic regions of a separate group of control SNPs associated with non-diabetic traits (e.g., Alzheimer’s disease, and white blood cell counts) were not enriched with local-ATAC-MVs. This enrichment analysis further confirmed the relevance of the local-ATAC-MVs discovered in the mouse for the diabetes-associated human GWAS SNPs.

**Figure 8:**
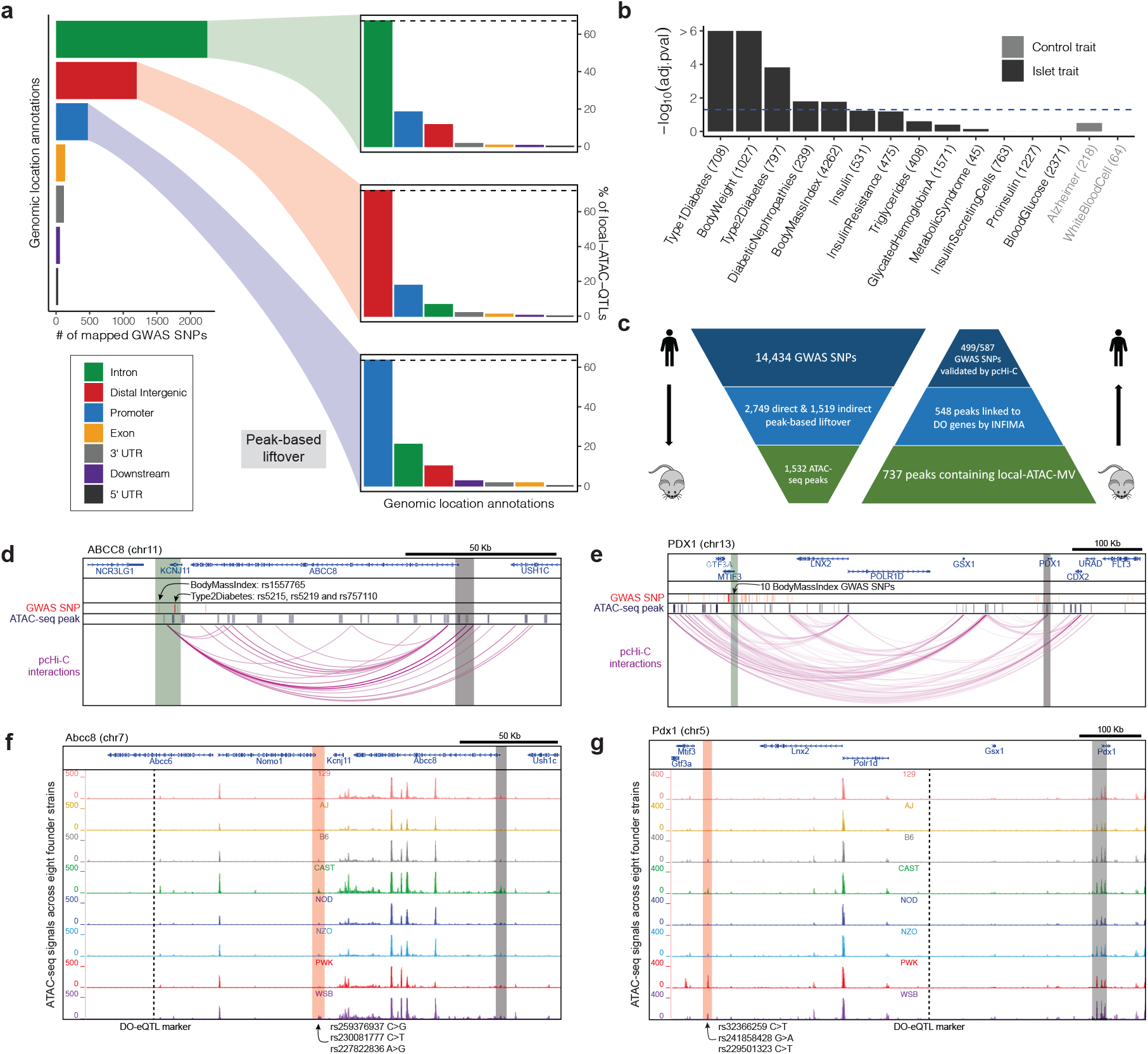
INFIMA generates candidate susceptibility genes for human GWAS SNPs. **a.** Comparison of genomic location annotations between human GWAS SNPs and the orthologous mouse genetic variants. Left: numbers of mapped GWAS SNPs within intronic, distal, promoter, exonic, UTR and downstream genomic locations. Right: Barplots of the local-ATAC-MVs mapping to the intronic, distal and promoter groups of GWAS SNPs, highlighting marked conservation of genomic location types. **b.** Mapped GWAS SNPs are enriched for local-ATAC-MVs. The numbers in parentheses depict the numbers of GWAS SNPs for each trait and the blue dashed line marks the threshold for the Bonferroni adjusted cutoff at 0.05. **c.** Summary of validation of INFIMA suggested susceptibility genes for human GWAS SNPs mapped to mouse with promoter capture Hi-C (pcHi-C). **d, f.** pcHi-C links *ABCC8* promoter to 4 GWAS SNPs which map to 3 mouse local-ATAC-MVs with INFIMA predicted effector gene *Abcc8*. **e, g.** Promoter capture Hi-C links *PDX1* promoter to 10 distal GWAS SNPs which map to 3 mouse local-ATAC-MVs with INFIMA predicted effector gene of *Pdx1*. **d, e.** Human genome depictions of interactions of distal GWAS SNPs (translucent green) with the *ABCC8* and *PDX1* promoters (translucent gray), together with the human ATAC-seq peaks. **f, g.** Mouse genome depiction of ATAC-seq signal for local-ATAC-MVs (translucent red) where INFIMA fine-maps DO-eQTL marker (dashed line) linked to genes *Abcc8* and *Pdx1* (promoters highlighted in translucent gray).

Next, we used INFIMA to predict effector genes of diabetes-associated GWAS SNPs. Among the 1,532 mouse ATAC-seq peaks syntenic to GWAS variants, 737 contained local-ATAC-MVs. Of these, 548 were causally linked to at least one DO-eQTL gene, with 18.1% linked to a single gene (Supplementary: Fig. S37). This generated a set of human gene orthologs as candidate effectors of GWAS SNPs. We next used human islet promoter capture Hi-C data (pcHi-C)^[64]^ and assessed whether pcHi-C interactions supported the inferred GWAS SNP-effector gene pairs (Supplementary: Fig. S38). First, we observed that the indirect peak-based lift-over strategy did not exhibit any discernible difference from the direct lift-over in terms of pcHi-C validation (Fisher’s exact test p-value = 0.848). Next, we compared INFIMA effector gene predictions for these human GWAS SNPs with two baseline strategies: (1) linking mouse ATAC-seq peaks syntenic to GWAS variants to their nearest genes instead of INFIMA predictions; (2) linking human GWAS SNPs to their nearest genes without going through model organism data and INFIMA predictions. We observed that INFIMA predictions were markedly better supported by the pcHi-C data (Fisher’s exact test p-values of 3.48e-96 and 1.20e-21 for comparisons of INFIMA predictions to strategy (1) and (2) respectively; Methods).

Overall, we identified putative effector genes for 587 GWAS SNPs, 499 of which were supported by the candidate effector gene promoter regions exhibiting significant Hi-C signal^[64]^ with either the corresponding GWAS SNPs or human ATAC-seq peaks at enhancer regions (Fig. 8c; Methods). Among these effector genes are *ABCC8*, *KCNJ11*, *PDX1*, *ADCY5*, and *KCNQ1*, which are recognized as pancreatic *β*-cell genes strongly associated with Type 2 Diabetes^[65,66]^. The *ABCC8* promoter is linked to a distal intergenic GWAS SNP rs1557765 (Body Mass Index) as well as three *KCNJ11* intronic GWAS SNPs rs5215, rs5219, and rs757110 (Type 2 Diabetes) by pcHi-C data. These three human SNPs are syntenic to rs25937937, rs230081777, and rs227822836 in mouse, and are identified by INFIMA as causal for a *Abcc8* DO-eQTL, the homologue to human *ABCC8* (Figs. 8d and f). In addition to nominating candidate effector genes, INFIMA analysis also facilitates comparison of potential impacts of human GWAS SNPs and their syntenic mouse local-ATAC-MVs on transcription factor binding. For example, atSNP search^[67]^ results on human SNPs rs5215 and rs1557765 indicate that both rs1557765 and rs5215 lead to better sequence motifs for TCF7L2 (atSNP p-values of 5.99e-3 and 6.33e-4 for motif enhancement) and, furthermore, rs5215 also results in a better sequence motif for YY1 (atSNP p-value of 1.68e-3, Supplementary: Fig. S39a). Similarly, their syntenic mouse local-ATAC-MVs rs227822836 and rs230081777 enhance the binding sites for orthologous Tcf712 and Yy1 (atSNP p-values of 2.03e-2 and 8.53e-3 for motif enhancement; Supplementary: Fig. S39b).

pcHi-C data supports a chromatin loop that links *PDX1*, deficiency of which associates with *β*-cell dysfunction^[68]^, to 10 GWAS SNPs rs1924074, rs9581853, rs9579083, rs9319366, rs9581854, rs4771122, rs12584061, rs12585587, rs9581856, and rs9579084 (also associates with Body Mass Index) at promoter and intronic regions of *MTIF3*. These GWAS SNPs are lifted-over to a mouse locus, with local-ATAC-MVs rs32366259, rs241858428, and rs229501323, and for which INFIMA identifies *Pdx1* as the potential effector (Figs. 8e and g). We further observe that TFAP2A, GABPA, and HIC1 motifs are disrupted while CREB1, NFYA, TP53, NKX3-2, and EGR1 motifs are enhanced by the aforementioned human GWAS SNPs and their syntenic mouse local-ATAC-MVs, suggesting orthologous TF bindings (Supplementary: Fig. S40 - S47).

In addition to these examples where the human GWAS SNPs with inferred effector genes are likely to enhance or disrupt TF binding sites, our results include cases where the SNPs exert their effects on expression through H3K27ac modification which is one of the enhancer defining histone modifications. An example of this is type 2 diabetes GWAS SNP rs11708067 for which INFIMA analysis identified *ADCY5* as the effector gene (Supplementary: Fig. S48). This SNP was shown to contribute to type 2 diabetes by disrupting an islet enhancer and, consequently, resulting in reduction of *ADCY5* expression^[69]^. In addition, *ADCY5* was also inferred as the effector gene for SNPs rs11708903, rs6438788, and rs4450740 associated with Blood Glucose & Insulin Secreting Cells and residing in the intronic region of *ADCY5*. Finally, supporting data for *KCNQ1*, a susceptibility gene for type 2 diabetes^[70]^, is provided in Supplementary: Fig. S49.

## 3 Discussion

While advances in genome sequencing improved the power of GWAS studies, elucidating which genes GWAS SNPs might be impacting is still a critical barrier for fully unleashing the power of GWAS. Recent large-scale and innovative efforts that leverage reference transciptome datasets to impute gene expression in GWAS cohorts and leverage co-localization with GWAS results have been successful in suggesting gene-level associations^[71–73]^. However, these studies are limited by the availability of reference transcriptomes in relevant tissues and accurate predictive models of gene expression. In a complementary approach, we leveraged model organism multi-omics data for this challenging task. Specifically, we developed INFIMA as a statistically grounded framework to capitalize on multi-omics functional data and fine-map model organism molecular quantitative trait loci. Application of INFIMA to DO mouse islet eQTLs fine-mapped previously identified eQTLs. Next, we asked whether INFIMA islet eQTL fine-mapping results could be transferred to human to infer effector genes of non-coding human GWAS SNPs. This reasoning is instigated by the observation that non-coding human GWAS SNPs associated with pancreatic islet functions are overwhelmingly enriched in synthenic accessible chromatin regions in islets of founder DO strains, suggesting potential functional relatedness among the two sets of non-coding regions. We utilized INFIMA resolved DO mouse SNP-effector gene linkages to infer effector genes for about fifteen thousand human GWAS SNPs. This application identified effector genes for 587 GWAS SNPs, linkages of 85% were supported by promoter capture Hi-C data of human islets. Notably, a limitation of pcHi-C data as the gold standard is the lack of specificity compared to, for example, large-scale CRISPR screening experiments. However, it currently serves as a widely used approach for identifying putative links^[74–76]^. The effector gene set included genes with well-established connections to islet functions (e.g., *ABCC8*, *KCNJ11*, *PDX1*, *ADCY5*, and *KCNQ1*) as well as novel candidates (e.g., *NFATC2IP*). While the ability to infer susceptibility genes for only 3.5% of the GWAS SNPs might appear low, this is due to several potential factors. First, by utilizing multi-omics data from islets, we are aiming to identify effector genes of diabetes associated GWAS variants in islets. This will inherently exclude SNPs that might be exerting their effects in other tissues. Second, the set of candidate regulatory regions (local-ATAC-MVs) that we have defined in founder strain islets excludes other known potential regulatory mechanisms (e.g., alternative transcriptional regulation and 3D interactions^[77,78]^) that the non-coding SNPs might be involved in. Third, only a subset of the trait-associated human GWAS SNPs are likely to be eQTLs^[79]^, and, furthermore, GWAS SNPs can mediate their effects through molecular mechanisms beyond expression modulation. These, in combination with potential organism-specific regulatory mechanisms, impact the extent of effector gene inference from human GWAS SNPs and fine-mapped model organism eQTL data. Despite these shortcomings, we showed with promoter capture Hi-C data validation that INFIMA, with the current lift-over strategies that we employed, can be a powerful transfer learning approach for exploring susceptibility genes of human GWAS loci. The lift-over strategies to identify syntenic non-coding regions between human and mouse are likely to benefit from recent analysis of cross-species enhancers^[80]^.

## 4 Conclusions

Model organism studies provide extensive resources for human GWAS; however, effective model organism data integration methods as well as reliable cross organism transfer learning frameworks are lagging behind. INFIMA provides a general framework for fine-mapping model organism molecular quantitative loci by integrating multiple functional data modalities. The availability of such fine-mapping results enables their transfer to the human genome to identify putative effector genes of GWAS variants. The current implementation of INFIMA excludes trans-eQTLs. As the ability to measure inter-chromosomal interactions matures, incorporating trans-eQTLs into INFIMA framework would be a natural extension. The INFIMA software is released at GitHub under the MIT license^[81]^, https://github.com/keleslab/INFIMA. The web application for INFIMA results are available at http://www.statlab.wisc.edu/shiny/INFIMA/.

## Supporting information

INFIMA Supplementary Information

## URLs

The INFIMA software https://github.com/keleslab/INFIMA; The web application for IN-FIMA results http://www.statlab.wisc.edu/shiny/INFIMA/; The processed data and results https://doi.org/10.5281/zenodo.4625293; The source code for reproducing the results https://github.com/ThomasDCY/INFIMA-paper; The DO mouse eQTL data https://churchilllab.jax.org/qtlviewer/attie/islets; ENCODE 15-state chromHMM data https://www.encodeproject.org/search/?type=Annotationannotation_type=chromatin+stateassembly=mm10files.file_type=bed+bed9; ENCODE H3K27ac and H3K4me3 ChIP-seq based classification of tissue-specific promoters/enhancers http://zlab-annotations.umassmed.edu/enhancers/ and http://zlab-annotations.umassmed.edu/promoters/; dbSNP142 ftp://ftp-mouse.sanger.ac.uk/current_snps/mgp.v5.merged.snps_all.dbSNP142.vcf.gz; The UCSC lift-over tool https://bioconductor.org/packages/liftOver/; The reciprocal chain file https://hgdownload-test.gi.ucsc.edu/goldenPath/hg19/vsMm10/reciprocalBest/.

### Authors’ contributions

S.K., M.K., and A.A. conceived the project. Sh.S., D.S., and K.S. performed ATAC-seq assays. G.C. contributed RNA-seq data. C.D. and S.K. analyzed ATAC-seq & RNA-seq datasets and performed footprint analysis. Su.S. and S.K. optimized the ATAC-seq data analysis pipeline. C.D., S.K. developed and evaluated the INFIMA model. L.L., X.L., F.J., and Y.L. contributed easy Hi-C data. C.D. and S.K. developed the initial version of the manuscript. C.D., S.K., and M.K. wrote the manuscript with input from other authors.

### 5 Methods

#### ATAC-seq sample preparation

The ATAC-seq samples were prepared using a selection of 50 average sized mouse islets. The islets were washed with 500 *μ*L of PBS at 4C and pelleted by centrifugation at 100 × g for 1 minute. 300 *μ*L of ATAC Lysis buffer (10 mM Tris-HCl pH 7.4, 10 mM NaCl, 3 mM MgCl2, 0.1% IGEPAL CA-630) was used to resuspend the islets. The islets were incubated for 20 minutes on ice. After incubating, the islets were lysed by trituration with a 25 gauge needle until intact islets were no longer visible, usually 6 triturations. The lysate was centrifuged at 500 x g for 10 minutes at 4C. This generated a crude nuclei pellet and a supernatant. The supernatant was discarded and the nuclei pellet was washed with 100 *μ*L of ATAC Lysis buffer in order to reduce cytoplasmic and mitochondrial contamination. This mixture was centrifuged at 500 × g for 10 minutes at 4C and the supernatant was removed. Per ATAC-seq sample, a mixture of 25 *μ*L 2x TDE buffer, 22.5 *μ*L nuclease-free water, and 2.5 *μ*L TDE1 transposase enzyme (Nextera DNA Library Prep kit, Illumina) was applied and incubated for 30 minutes in a 37C water bath. The samples were then purified using a MinElute Reaction cleanup kit (Qiagen) and eluted using two sequential aliquots of 10 *μ*L EB buffer. After purification all ATAC samples were kept at −80C. All ATAC-seq samples were transposed and frozen prior to preparing all libraries. Libraries were amplified using 20 *μ*L of ATAC sample, 2.5 *μ*L Primer-1 (Ad1 noMX, 25 *μ*M working stock), 2.5 *μ*L Primer-2 (Ad2.X, 25 *μ*M working stock), and 25 *μ*L of NEBNext High Fidelity 2x PCR Master Mix. Each ATAC sample was amplified by 12 cycles which was determined by qPCR to be saturating for the libraries. The PCR thermocycler was set to 72C for 5 minutes, 98C for 30 seconds, and then 12 total cycles of 98C for 10 seconds, 63C for 30 seconds, 72C for 1 minute. After amplification the libraries were purified using MinElute PCR purification cleanup kit (Qiagen). The libraries were sequenced to a depth of 134.8 ± 8.2 million reads using paired-end 125 bp reads on a HiSeq2500 (Illumina) at the University of Wisconsin Biotechnology Center DNA Sequencing Facility.

#### ATAC-seq data analysis

##### Alignment of ATAC-seq reads

Illumina Nextera adapters were trimmed with cutadapt (version 2.0)^[82]^ using the option “-q 30 –minimum-length 36”. Paired-end ATAC-seq reads were aligned to the mouse genome assembly (mm10) with bowtie2 (version 2.3.4.1)^[83]^ with option “-X 800 –no-mixed –no-discordant”. For each sample, unmapped reads were filtered out by SAMtools view (version 1.8)^[84]^ with option “-F 4” and mitochondrial reads were removed. Duplicated reads were removed with Picard tools (version 2.9.2)^[85]^. This resulted in an average of 77.7 ± 4.1 million reads per sample. TSS enrichment analysis was performed with ataqv^[86]^.

##### Generation of a master peak list from the ATAC-seq samples

Peaks from individual and pooled samples across sexes of each strain were identified using MOSAiCS^[27,28]^ at FDR of 0.05. Blacklisted regions (see URLs) and Chr Y regions were filtered. We employed IDR analysis^[29]^ to obtain reproducible sets of peaks between male and female samples at IDR of 0.05 and leveraged “SignalValue” and “p-value” outputs from IDR analysis as measures of peak-level signal to noise. The “SignalValue” output was normalized across strains by multiplying 10^8^/ (# of reads) to adjust for differences in the sample sequencing depths. IDR identified peaks from the pooled peak sets were trimmed to exclude peaks with the lowest 10% “SignalValue” for each strain and then merged to form the master peak list across all strains. “SignalValue” and “− log_10_(p-value)” columns were aggregated as “MeanSignal” and “MeanP” in the master peak list.

Strain-specific ATAC-seq peaks tended to have lower ATAC-seq signals compared to peaks present in multiple strains (Supplementary: Fig. S4). We mitigated the potential for this bias by trimming the combined peak list to maximize the overlap of the trimmed set with the ENCODE chromHMM annotations depicting non-quiescent regions of the genome (See URLs; Supplementary: Figs. S4, S5, and S6; Supplementary: Supplementary Notes). We reasoned that ATAC-seq peaks across the strains should largely be within non-quiescent chromatin states. We utilized 15-state chromHMM data for mm10 across 12 tissues from the ENCODE portal^[87]^ and annotated the master peak list according to the pooled set of the non-quiescent chromHMM regions across the 12 tissues. For each level of “Total”, i.e., the number of strains a master peak is identified in, we varied two tuning parameters: percentile of “MeanSignal” and percentile of “MeanP”, both of which varied in {0, 1, ..., 50}. Supplementary: Fig. S5 depicts the heatmaps for the percentage of non-quiescent peaks and the percentage of remaining peaks as a function of these two trimming parameters. In order to maximize these two quantities, we chose tuning parameters for each level of the “Total” and generated the trimmed master peak list. Finally, the reference strain B6 did not have more strain-specific peaks compared to other strains regardless of the trimming procedure, further demonstrating that alignments to the reference mouse genome did not amplify B6 ATAC-seq peak signals (Supplementary: Table S3, Supplementary: Fig. S7).

##### Differential accessibility analysis

The ATAC-seq count matrix for the set of master peaks was computed by the R package ChromVAR^[88]^. We used DESeq2 ^[89]^ to identify strain effects (the model “~ strain” vs. the null model) and sex effect (the model “~ sex + strain” vs. “~ strain”) by corresponding likelihood ratio tests at FDR of 0.05.

##### Footprint analysis of ATAC-seq peaks

We utilized PIQ^[33]^ to identify footprints of the 1,316 curated JASPAR motifs^[90]^ in B6 ATAC-seq samples with purity score cutoff 0.75, i.e., TF occupancy probability. To investigate whether ATAC-peaks were enriched for footprints of TFs highly expressed in islets, we first quantified the ATAC-seq signal genome-wide at base pair resolution by counting the 5’ end Tn5 cut sites for each strain and normalized the cut sites by the sequencing depths. Then, for each potential transcription factor binding site along the genome, we computed the average Tn5 cuts at (1) the binding site, (2) 25 bp flanking regions of the binding site, and (3) 26 - 50 bp flanking regions of the binding site. We adapted the footprint depth (FPD) metric^[47]^ as the proportional decrease in cut sites at the binding site compared to flanking regions (Supplementary: Fig. S21). The footprint profiles for the individual binding sites were computed from the base pair level ATAC-seq signal in B6 ATAC-seq samples and aggregated for each individual motif. We evaluated the significance of average FPD of each islet TF by comparing it to average FPDs of motifs that are similar in width (width within ± 1 of the islet TF motif width) and information content (information content within ± 0.2 of the islet TF motif information content). A randomization test was performed to evaluate the collective enrichment of islet TFs (Supplementary: Supplementary Notes).

##### Identification of local-ATAC-MVs

In order to evaluate the impact of SNPs on ATAC-seq signal, we first extracted genetic variants within differential ATAC-seq peaks for the eight founder strains from the dbSNP (v142) database (see URLs,^[91]^) with the R package VariantAnnotation (version 1.34.0)^[92]^. Retaining only the SNPs with “FILTER = PASS” and “QUAL = 999” resulted in 630,349 SNPs. In order to identify genetic variants genotypes of which are associated with the ATAC-seq signal, we conducted a permutation test and retained for each differential ATAC-seq peak only the SNP which associated the best with the local ATAC-seq signal while including all the SNPs with the same exact best association statistics. This resulted in 22,200 ATAC-seq peaks harboring a total of 47,062 local-ATAC-MVs at FDR of 0.05, with an average (median) of 2.1 (1.0) local-ATAC-MVs per peak.

#### *In silico* mutation and footprint analysis

##### Variant-level comparative footprint analysis

We applied atSNP^[93]^ to 47,062 local-ATAC-MVs with the 1,316 curated JASPAR motifs^[90]^ and quantified the *in silico* effect of SNPs on TF binding by labeling SNP-motif combinations with atSNP pval_rank < 0.05 as significant.

Next, to quantify the impact of SNPs on the realized ATAC-seq footprints, for each SNP × motif interaction, FPD with/without SNP were computed by aggregating the results for strains with/without the alternative allele. This ensured the disruption/enhancement of motif by a SNP to be consistent with a decrease/increase in FPD. In order to evaluate whether the change in FPD (ΔFPD) due to the SNP is significant, we generated motif-specific empirical null distributions of ΔFPD by treating insiginificant results from atSNP as the null set since this approximated the distribution of ΔFPD when the SNP is not affecting the motif. Only the SNP-motif combinations with pval_fpd < 0.05 were retained for the downstream analysis.

Accounting for both the *in silico* effect of SNP on TF binding and change in ATAC-seq FPD, resulted in 1,211,807 candidate SNP-motif interactions with consistent changes across the two metrics (640,038 Gain of function combinations: pval_ref > 0.05, pval_snp ≤ 0.05, ΔFPD > 0; 571,769 Loss of function combinations: pval_ref ≤ 0.05, pval_snp > 0.05, ΔFPD < 0). Finally, for each SNP, we recorded the minimum pval_fpd as the p-value for the null hypothesis that the SNP is not affecting any TF binding. Collectively, we identified 8,029 significant SNP × motif interactions comprising 1,350 SNPs and 1,196 motifs (FDR of 0.05).

#### RNA-seq sample preparation

Islet RNA profiling methods are described in detail in^[14]^.

#### RNA-seq data analysis

##### Quantification of transcript abundance

We used RSEM^[48]^ with GENCODE vm18 ^[94]^ gene annotation and obtained the gene expression count matrix across protein coding genes on Chromosomes 1-19, and X. Genes with the lowest 10% variance across the samples were removed from the downstream analysis. Upper quartile normalization^[95]^ and retaining the genes with non-zero counts in at least 85% of the samples resulted in 13,568 protein-coding genes.

##### Association analysis of founder local-ATAC-MVs and gene expression

We applied Matrix-EQTL^[96]^ with default settings to all local-ATAC-MVs and obtained 96,309 associated local-ATAC-MV and gene pairs (34,711 distinct local-ATAC-MVs, only *cis* regulatory local-ATAC-MVs were considered, 1 Mb window) at FDR of 1e-5.

#### INFIMA implementation details

##### INFIMA model fitting with an Expectation-Maximization algorithm

We estimated the INFIMA parameters with maximum likelihood using an Expectation-Maximization (EM) algorithm. We provide below the detailed derivations. Let **Γ**_**g**_ = (**Θ**_**g**_, **a**_**0**_, **b**_**0**_, **a**_**1**_, **b**_**1**_, *γ*) denote the full set of model parameters and **1**_**g,k**_ be a *p*_*g*_×1 vector with the *k*^*th*^ entry equal to 1 and 0 elsewhere. The joint likelihood of the data 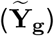 and the latent variables, conditional on features **X**_**g**_ extracted from founder RNA-seq and ATAC-seq, for **Z**_**g**_ = **1**_**g,k**_ is given by

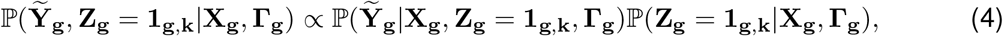

where the first term is given by

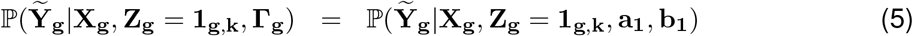

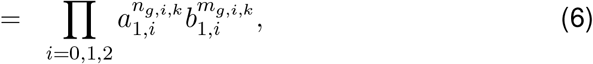

and the second term is given by

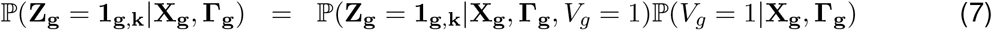

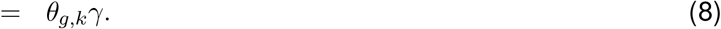

Similarly, the joint likelihood when **Z**_**g**_ = **0** is then

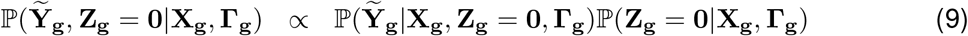

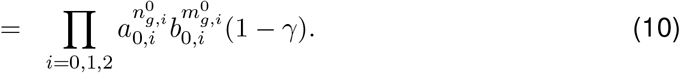

We next derive the full parameter joint posterior distribution given the latent variables **Z**_**g**_, *V*_*g*_ as

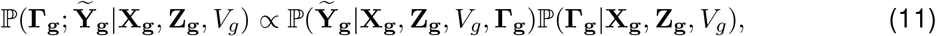

where

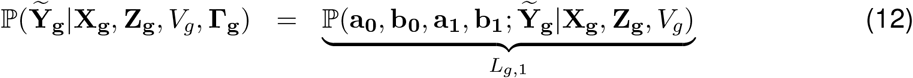

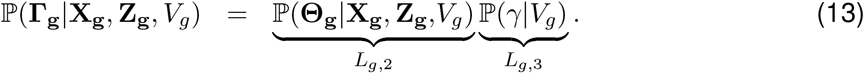

With the combined generative model, we have

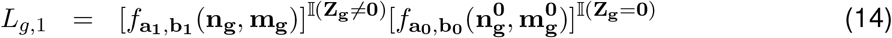

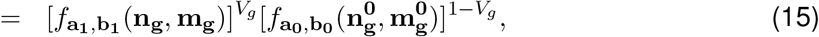

where *f*_.,._ denotes the product of Multinomial probability mass functions with appropriate parameters. The log likelihood aggregated over *g* ∈ {1, 2, ..., *G*} is given by

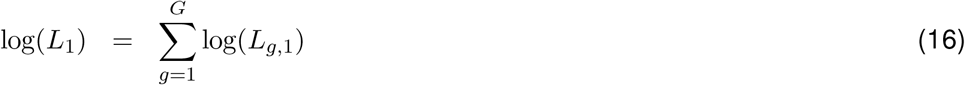

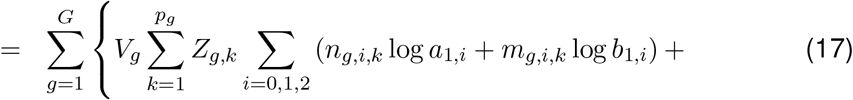

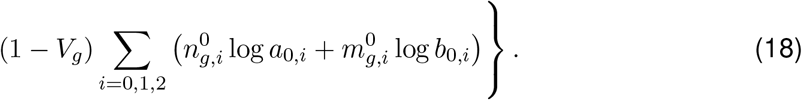

We define the weighted sums of the edit distance random variables **n**_**g**_, 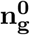, **m**_**g**_, 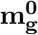 as

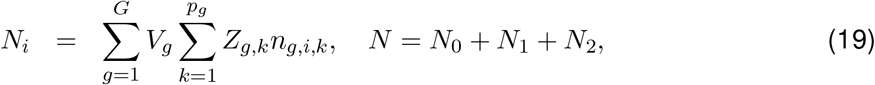

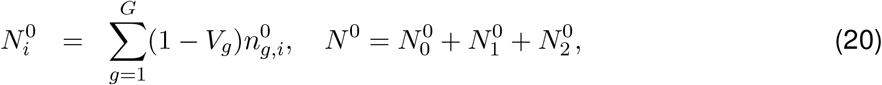

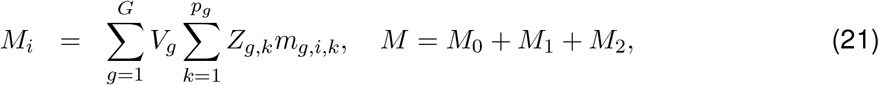

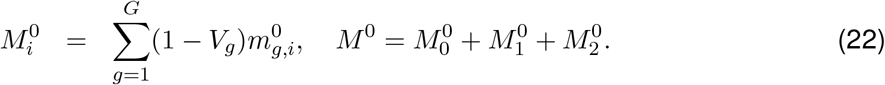

Then, the Maximum Likelihood Estimators (MLEs) of the parameters are given by:

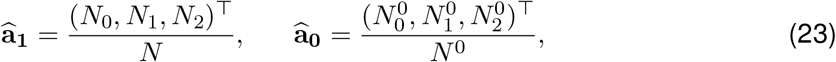

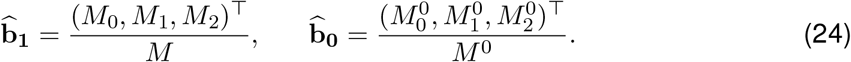

We note that *L*_*g*,2_ is the posterior distribution of **Θ**_**g**_. By the Dirichlet-Multinomial conjugacy, we have

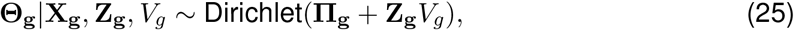

and the maximum a posteriori (MAP) estimator can be computed as

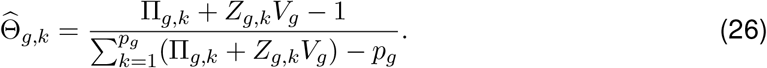

Maximizing *L*_*g*,3_ = ℙ(*γ*|*V*_*g*_) with respect to the prior probability *γ* that an association is driven by causal SNP, we get 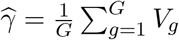.

In the DO-eQTL application, the INFIMA model was fit with the EM algorithm described in Algorithm 1, where *V*_*g*_ and *Z*_*g,k*_ values in the above equations were imputed in the E-step. Multiple initial values of parameters were employed to avoid local optima.

##### Trinarization of allelic expression effect sizes into allelic patterns

For the DO-eQTL data **Y**_**g**_, we first standardized the 8 × 1 vector to [0,1] and subtracted the allelic expression effect of the reference strain B6. We then trinarized the entries with values > 0.2, < −0.2 to 1, −1 respectively, and set other entries to 0 to obtain 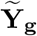. The cutoffs were selected by balancing the number of entries with the 3 values. The same trinarization scheme was applied to the normalized founder gene expression vector 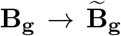 as well. For each row of the founder RNA-seq genotype effect matrix **E**_**g**_, if the effect size from the marginal regression of gene expression on the genotype was significant at level 0.05, we replaced the effect size with 1 or −1 depending on the sign of the effect size; otherwise, the effect size was replaced by 0. Therefore, we obtained 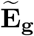 and 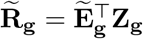. Fig. 5f illustrates a specific example in detail. *Distance prior.* A well known bias of Hi-C data is that Hi-C signal decreases exponentially as the distance between promoters and enhancers increases^[97]^. In order to avoid the bias towards the local-ATAC-MVs closest to the gene promoter, we chose not to penalize the distance until 250 Kb. When distance is above 250 Kb, the score function has a decreasing trend in order to slightly favor closer local-ATAC-MVs. We set the window size *W* equal to 1 Mb and defined the distance score function as *D*(*x*) = 0.5 if *x* ≤ 0.25 Mb; 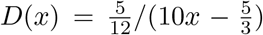 if *x >* 0.25 Mb, where *x* is the distance between local-ATAC-MV and DO gene promoter. As a component of the prior **Π**_**g**_, the maximum value of distance score **D**_**g**_ is 0.5, which serves as a “tie-breaker” rather than overwhelming the other three components (Fig. 5a).

###### Algorithm 1

INFIMA Model Fitting with Expectation-Maximization

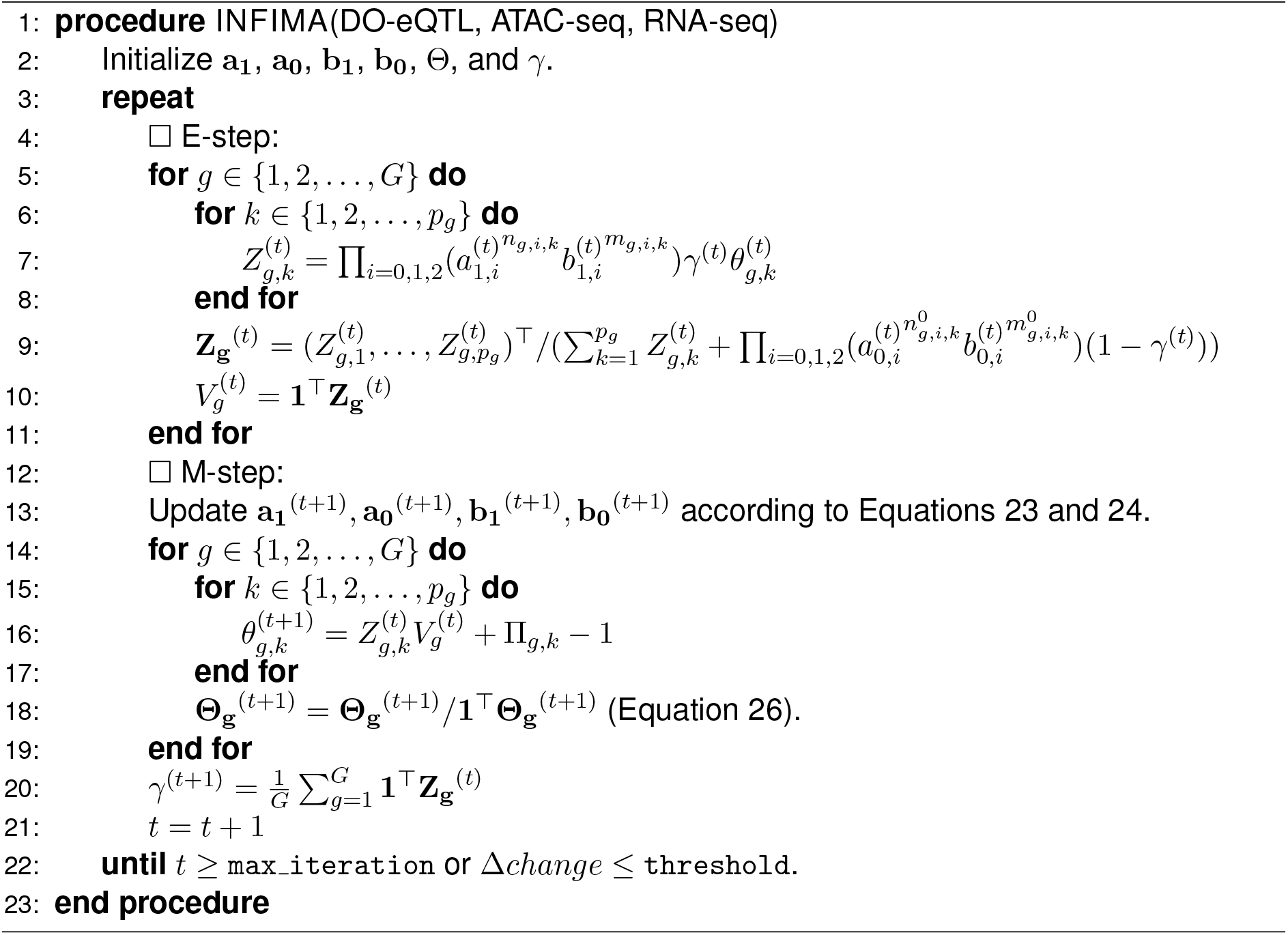

##### Pseudocounts for the edit distance random variables

To promote the consistency between the trinarized DO-eQTL data and founder data, i.e., to tilt the edit distance random variables to favor lower values, we utilized pseudocount parameters *λ*_0_ = 0.1, *λ*_1_ = 0.01, and *λ*_2_ = 0 for the multinomial edit distance random variables. Specifically, pseudocounts *λ*_*i*_*p*_*g*_ were added to the weighted sums of edit distance variables (Eq. 19 to 22) in estimation of **a**_**0**_, **b**_**0**_, **a**_**1**_, and **b**_**1**_ as:

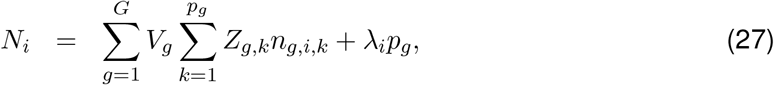

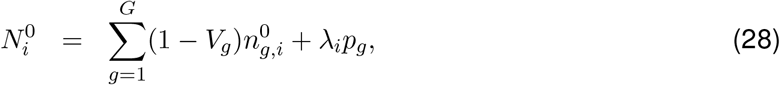

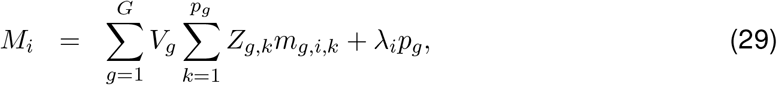

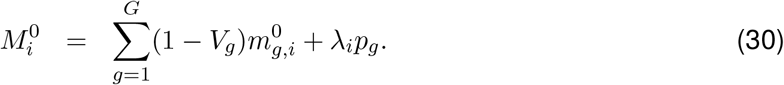

Under *λ*_0_ >> *λ*_1_ >> *λ*_2_, INFIMA formulation promotes the resulting causal SNPs to have consistent relationships between the founder data and the DO-eQTL data; therefore, the ordering of the SNPs is relatively insensitive to the actual values of these pseudocount parameters.

#### Data-driven simulations

In order to simulate realistic data for our evaluations, we leveraged the parameters estimated by the INFIMA on the DO-eQTL data fit with all the summarized data from ATAC-seq, local-ATAC-MVs, and RNA-seq data. We used these parameter values as well as the actual summarized ATAC-seq, local-ATAC-MVs, and RNA-seq to simulate *V*_*g*_, **Z**_**g**_, and **Y**_**g**_ from the plate model in Fig. 5b. We varied the informativeness level of the summarized data by varying the prior parameter Π_*g,k*_ := *F*_*g,k*_ + *D*_*g,k*_ +|cor(**A**_**g,k**_, **E**_**g,k**_)| + |cor(**A**_**g,k**_, **B**_**g**_)| + 1 according to the following three settings:

NI: The prior parameter Π_*g,k*_ is set to be 1 for all candidate SNPs, corresponding to an uninformative prior.
MI: The prior parameter Π_*g,k*_ set to its observed value in the actual data and accommodates multiple SNPs with *F*_*g,k*_ = 1. Multiple SNPs are affecting footprints under this setting. Causal SNPs are distinguished by other components of the prior parameter.
HI: *F*_*g,k*_ is set to 10 for a randomly selected SNP *k* and 0 for other SNPs. Under this setting, the SNPs that affect footprints are more likely to be chosen as causal due to the dominant contribution of the footprint component.

Statistical power for *V*_*g*_ was calculated at FDR of 0.05 by using a direct posterior probability approach^[98]^.

#### Linking human GWAS SNPs to mouse islet ATAC-seq peaks

The peak-based lift-over consisted of two steps: (1) direct (2) indirect (Supplementary: Fig. S36b). After removing black-listed and chr Y human ATAC-seq peaks^[58]^, we obtained 156,861 human islet ATAC-seq peaks. For indirect mapping, we used “nearest()” function in “GenomicAlignments” R package^[99]^ to link GWAS SNPs to their nearest human ATAC-seq peaks within 10 Kb distance. We used ‘liftOver()’ function in “rtracklayer” R package^[100]^ and the hg19 to mm10 reciprocal chain file (see URLs). For each human genomic region, we merged gaps less than 10 bp among its mapped regions in the mouse genome and selected the one with maximum width as the syntenic region. We then linked these syntenic regions to their nearest mouse ATAC-seq peaks within 10 Kb distance. The distance constraints aided to remove potential false positives to preserve conservation of genomic compartments between the syntenic regions of the two organisms. We observed a decline in level of conservation without imposing the distance constraints (Supplementary: Fig. S50).

#### Enrichment analysis of human GWAS SNPs associated with islet function traits

We carried out an enrichment analysis for the associated SNPs of islet function related GWAS traits with more than 40 SNPs. Enrichment p-values were calculated based on a resampling based null distribution that matched the phylogenic conservation score, width, and chromosomal distribution of the syntenic regions of each GWAS trait. Specifically, for each trait, we sampled the same number of random syntenic regions as the size of the set lifted over to mouse genome by matching the phylogenic conservation score, width, and chromosomal distribution of the sampled regions to those of the actual syntenic regions. The random syntenic regions were mapped to mouse ATAC peaks within 10 Kb distance, and the overlap with the local-ATAC-MVs were recorded. Repeating this procedure one million times generated a null distribution for the actual observed number of local-ATAC-MVs that mapped to GWAS. The resulting enrichment p-values were corrected for multiple testing with the Bonferroni procedure at the significance level of 0.05.

#### Validation of INFIMA predicted SNP-effector gene linkages with promoter capture Hi-C

For validation purposes, we filtered out 8 out of 1,540 mouse ATAC-seq peaks because the human ortholog of the genes that they were linked to resided in different chromosomes than the corresponding GWAS SNPs that they mapped to. Then, we processed the INFIMA results that fine-mapped 737 local-ATAC-MV containing peaks that corresponded to syntenic regions of human GWAS SNPs. INFIMA resulted in mappings for 587 GWAS SNPs (548 local-ATAC-MV containing peaks) by considering the local-ATAC-MVs with aggregated posterior probability of being causal larger than 0.80 and with a credible set less than 50% of the all the candidate SNPs. We leveraged 175,784 significant promoter capture Hi-C contacts from^[64]^ for validation of the inferred links. With a median bin size ~4 Kb, the median interaction distance of the pcHi-C data is ~300 Kb. We required one end of pcHi-C interaction to be within 10 Kb upstream and 2 Kb downstream around TSS of human orthologous genes while the other end of pcHi-C to reside within 10 Kb distance of GWAS SNPs and human ATAC-seq peaks. We identified 346 GWAS SNPs that were supported by pcHi-C through at least one effector gene. Furthermore, at least one LD partner (R^2^ > 0.8, 1000 Genomes Phase 3 v5 European population, SNiPA v3.3 ^[101]^) of the 153 GWAS SNPs were in contact with the inferred effector genes. Comparison of INFIMA predictions to the baseline strategies was carried out with a Fisher’s exact test.

## Footnotes

1 rank() function in R was used with ties.method = “average”, and then normalized the resulting score by *p*_*g*_. The rank score is ∈ [0, 1] and larger magnitudes correspond to higher ranks.

