## Supplementary material for "INFIMA leverages multi-omics model organism data to identify effector genes of human GWAS variants": INFIMA Supplementary Information

**2 Supplementary Tables** 5

**3 Supplementary Figures** 7

### 1 Supplementary Notes

#### 1.1 ATAC-seq library amplification primers

Ad1_noMX
AATGATACGGCGACCACCGAGATCTACACTCGTCGGCAGCGTCAGATGTG
Ad2.1_TAAGGCGA
CAAGCAGAAGACGGCATACGAGATTCGCCTTAGTCTCGTGGGCTCGGAGATGT
Ad2.2_CGTACTAG
CAAGCAGAAGACGGCATACGAGATCTAGTACGGTCTCGTGGGCTCGGAGATGT
Ad2.3_AGGCAGAA
CAAGCAGAAGACGGCATACGAGATTTCTGCCTGTCTCGTGGGCTCGGAGATGT
Ad2.4_TCCTGAGC
CAAGCAGAAGACGGCATACGAGATGCTCAGGAGTCTCGTGGGCTCGGAGATGT
Ad2.5_GGACTCCT
CAAGCAGAAGACGGCATACGAGATAGGAGTCCGTCTCGTGGGCTCGGAGATGT
Ad2.6_TAGGCATG
CAAGCAGAAGACGGCATACGAGATCATGCCTAGTCTCGTGGGCTCGGAGATGT
Ad2.7_CTCTCTAC
CAAGCAGAAGACGGCATACGAGATGTAGAGAGGTCTCGTGGGCTCGGAGATGT
Ad2.8_CAGAGAGG
CAAGCAGAAGACGGCATACGAGATCCTCTCTGGTCTCGTGGGCTCGGAGATGT
Ad2.9_GCTACGCT
CAAGCAGAAGACGGCATACGAGATAGCGTAGCGTCTCGTGGGCTCGGAGATGT
Ad2.10_CGAGGCTG
CAAGCAGAAGACGGCATACGAGATCAGCCTCGGTCTCGTGGGCTCGGAGATGT
Ad2.11_AAGAGGCA
CAAGCAGAAGACGGCATACGAGATTGCCTCTTGTCTCGTGGGCTCGGAGATGT
Ad2.12_GTAGAGGA
CAAGCAGAAGACGGCATACGAGATTCCTCTACGTCTCGTGGGCTCGGAGATGT
Ad2.13_GTCGTGAT
CAAGCAGAAGACGGCATACGAGATATCACGACGTCTCGTGGGCTCGGAGATGT
Ad2.14_ACCACTGT
CAAGCAGAAGACGGCATACGAGATACAGTGGTGTCTCGTGGGCTCGGAGATGT
Ad2.15_TGGATCTG
CAAGCAGAAGACGGCATACGAGATCAGATCCAGTCTCGTGGGCTCGGAGATGT
Ad2.16_CCGTTTGT
CAAGCAGAAGACGGCATACGAGATACAAACGGGTCTCGTGGGCTCGGAGATGT
Ad2.17_TGCTGGGT
CAAGCAGAAGACGGCATACGAGATACCCAGCAGTCTCGTGGGCTCGGAGATGT
Ad2.18_GAGGGGTT
CAAGCAGAAGACGGCATACGAGATAACCCCTCGTCTCGTGGGCTCGGAGATGT
Ad2.19_AGGTTGGG
CAAGCAGAAGACGGCATACGAGATCCCAACCTGTCTCGTGGGCTCGGAGATGT
Ad2.20_GTGTGGTG
CAAGCAGAAGACGGCATACGAGATCACCACACGTCTCGTGGGCTCGGAGATGT
Ad2.21_TGGGTTTC
CAAGCAGAAGACGGCATACGAGATGAAACCCAGTCTCGTGGGCTCGGAGATGT
Ad2.22_TGGTCACA
CAAGCAGAAGACGGCATACGAGATTGTGACCAGTCTCGTGGGCTCGGAGATGT
Ad2.23_TTGACCCT

CAAGCAGAAGACGGCATACGAGATAGGGTCAAGTCTCGTGGGCTCGGAGATGT
Ad2.24_CCACTCCT
CAAGCAGAAGACGGCATACGAGATAGGAGTGGGTCTCGTGGGCTCGGAGATGT

#### 1.2 Irreproducible discovery rate (IDR) analysis

We identified regions of accessible chromatin with MOSAiCS for both separate and pooled samples of each founder strain at false discovery rate (FDR) of 0.05 and applied irreproducible discovery rate (IDR) analysis at IDR of 0.05 to generate peak sets of each strain.

#### 1.3 Quality trimming of ATAC-seq master peaks

Quality trimming addressed a potential bias revealed by the positive correlation of the number of strains shared by an ATAC-seq peak with the ATAC-seq signal. Specifically, we trimmed the master peak list to maximize the overlap of the peaks with 15-state chromHMM annotations across an existing large collection of mouse tissues from the ENCODE project. In total, 12 tissues from mouse were available: liver e16.5, heart e16.5, lung e16.5, kidney e16.5, forebrain e16.5, midbrain e16.5, stomach e16.5, intestine e16.5, neural tube e15.5, hindbrain e13.5, limb e11.5, embryonic facial e13.5 (Supplementary Fig. S5 and S4).

#### 1.4 Promoter/Enhancer annotation of ATAC-seq master peaks

We overlapped ATAC-seq master peaks with ChIP-seq based promoter/enhancer lists from ENCODE and aggregated the results over existing mouse tissues (pancreatic islet was not available). The following datasets were included: embryofacial (H3K4me3 e14.5; H3K27ac e14.5), forebrain (H3K4me3 e14.5; H3K27ac e14.5), hindbrain (H3K4me3 e14.5; H3K27ac e14.5), lung (H3K4me3 e14.5; H3K27ac e14.5), midbrain (H3K4me3 e14.5; H3K27ac e14.5), heart (H3K4me3 e16.5; H3K27ac e16.5), intestine (H3K4me3 e16.5; H3K27ac e16.5), kidney (H3K4me3 e16.5; H3K27ac e16.5), limb (H3K4me3 e15.5; H3K27ac e15.5), liver (H3K4me3 e16.5; H3K27ac e16.5), stomach (H3K4me3 e16.5; H3K27ac e16.5).

#### 1.5 Collective enrichment of islet $\beta$-cell TFs

For each of the 744 human or mouse JASPAR 2020 motifs, we computed the footprint depth (FPD, Supplementary Fig. S21) for aggregated footprint profiles in B6 ATAC-seq samples. In order to evaluate the collective enrichment of the known $\beta$-cell TFs (PB0042.1;Mafk_1, PB0146.1;Mafk_2, PH0131.1;Pax4, PH0111.1;Nkx2-2, PB0015.1;Foxa2_1, PB0119.1;Foxa2_2, PH0132.1;Pax6, MA0132.1;Pdx1, PH0118.1;Nkx6-1_1, PH0119.1;Nkx6-1_2), we first treated their averaged FPD as observed statistic. We then applied a randomization test by uniformly drawing similar TFs for each $\beta$-cell TF (width difference $\leq 1$ and information content difference $\leq 0.2$). We repeated the randomization for 100,000 times and computed the average FPD during each iteration. A p-value was obtained by comparing the observed averaged FPD to those obtained from the randomized samples.

#### 1.6 Application of DAP-G and SuSiE for fine-mapping DO mice eQTLs

A standard application of human GWAS fine-mapping methods to DO-eQTL results considers all the SNPs around the eQTL marker as candidates. As a showcase, we consider the *Adcy5* locus with 27,307 SNPs within the 1 Mb of the eQTL marker of *Adcy5* (representative LD structure for 4, 616 SNPs within the 200Kb of the marker is provided in Fig. S31). The top signal cluster estimated by DAP-G contained 3,590 SNPs with cluster Posterior Inclusive Probability (PIP) of 0.434 and an average LD of 0.956. SuSiE did not output any credible sets for any coverage level. The maximum estimated SNP PIPs for both approaches were less than 0.003. Next, we utilized the multi-omics prior for the 27,307 SNPs by classifying them as: (i) not within an ATAC-seq peak; (ii) within an ATAC-seq peak but is not a local-ATAC-MV; (iii) local-ATAC-MV. For (iii), we utilized the multi-omics priors ($\mathbf{\Pi_g}$) that we have used for INFIMA and set the priors to 0.5 and 1, for (i) and (ii), respectively. However, the outputs from both DAP-G and SuSiE were almost identical with or without using a prior, further supporting that the level of LD in the DO mice hampers a standard application of fine-mapping methods designed for human genetic studies.

#### 2 Supplementary Tables

Table S1: Founder ATAC-seq data alignment rates. In this manuscript, 129, AJ, B6, CAST, NOD, NZO, PWK, and WSB are short for 129S1_SvImJ, A_J, C57BL/6J, CAST_EiJ, NOD_ShiLtJ, NZO_HlLtJ, PWK_PhJ, and WSB_EiJ respectively.

| | Sample | AlignOnce % | AlignMultiple % | chrM % | Duplicate % |
|---|---|---|---|---|---|
| 1 | 129-F | 66.42 | 29.74 | 28.71 | 11.80 |
| 2 | 129-M | 57.96 | 29.19 | 41.65 | 17.74 |
| 3 | AJ-F | 55.95 | 37.11 | 41.75 | 18.86 |
| 4 | AJ-M | 55.00 | 38.83 | 37.05 | 16.31 |
| 5 | B6-F | 55.47 | 32.22 | 35.35 | 15.68 |
| 6 | B6-M | 59.47 | 34.01 | 16.46 | 9.14 |
| 7 | CAST-F | 69.99 | 23.84 | 18.57 | 11.41 |
| 8 | CAST-M | 68.34 | 25.37 | 19.83 | 8.41 |
| 9 | NOD-F | 58.39 | 32.55 | 33.39 | 17.79 |
| 10 | NOD-M | 63.50 | 32.21 | 29.16 | 11.51 |
| 11 | NZO-F | 66.15 | 26.68 | 19.33 | 8.94 |
| 12 | NZO-M | 66.60 | 24.45 | 12.72 | 7.75 |
| 13 | PWK-F | 55.80 | 35.56 | 34.35 | 19.16 |
| 14 | PWK-M | 57.01 | 34.25 | 25.27 | 13.49 |
| 15 | WSB-F | 71.38 | 24.97 | 21.23 | 7.81 |
| 16 | WSB-M | 62.91 | 24.77 | 20.93 | 7.60 |

Table S2: Comparison of rate of alignments to reference genome (mm10) and personalized genomes.

| | Sample | AlignReference % | AlignPersonalized % |
|---|---|---|---|
| 1 | 129-F | 96.16 | 96.38 |
| 2 | 129-M | 87.16 | 87.23 |
| 3 | AJ-F | 93.06 | 93.17 |
| 4 | AJ-M | 93.83 | 93.97 |
| 5 | B6-F | 87.64 | 87.64 |
| 6 | B6-M | 93.44 | 93.44 |
| 7 | CAST-F | 93.84 | 96.12 |
| 8 | CAST-M | 93.71 | 95.98 |
| 9 | NOD-F | 90.94 | 91.11 |
| 10 | NOD-M | 95.71 | 95.92 |
| 11 | NZO-F | 92.84 | 93.03 |
| 12 | NZO-M | 91.05 | 91.36 |
| 13 | PWK-F | 91.36 | 94.13 |
| 14 | PWK-M | 91.26 | 94.59 |
| 15 | WSB-F | 96.35 | 96.63 |
| 16 | WSB-M | 87.69 | 87.97 |

Table S3: Number of strain-specific ATAC-seq peaks before and after quality trimming.

| | Strain | BeforeTrim | AfterTrim |
|---|---|---|---|
| 1 | 129 | 368 | 105 |
| 2 | AJ | 242 | 19 |
| 3 | B6 | 1048 | 103 |
| 4 | CAST | 15894 | 1406 |
| 5 | NOD | 995 | 45 |
| 6 | NZO | 244 | 83 |
| 7 | PWK | 1924 | 430 |
| 8 | WSB | 5914 | 813 |

Table S4: P-values from pairwise Kolmogorov-Smirnov tests between cumulative distribution curves of the five fine-mapping strategies. "Most Likely" and "Least Likely" refer to INFIMA most and least likely predictions, respectively.

| **Strategy** | **Most Likely** | **Least Likely** | **Random** | **Closest to Marker** |
|---|---|---|---|---|
| **Least Likely** | $<$2.2e-16 | | | |
| **Random** | $<$2.2e-16 | $<$2.2e-16 | | |
| **Closest to Marker** | $<$2.2e-16 | $<$2.2e-16 | 1.0e-15 | |
| **Closest to Gene** | $<$2.2e-16 | $<$2.2e-16 | $<$2.2e-16 | $<$2.2e-16 |

Table S5: Kullback–Leibler (KL) divergence between density curves of the five fine-mapping strategies. "Most Likely" and "Least Likely" refer to INFIMA most and least likely predictions, respectively.

| Strategy | Most Likely | Least Likely | Random | Closest to Marker | Closest to Gene |
|---|---|---|---|---|---|
| **Most Likely** | 0 | 0.548 | 0.248 | 0.121 | 0.080 |
| **Least Likely** | 0.464 | 0 | 0.091 | 0.189 | 0.419 |
| **Random** | 0.236 | 0.098 | 0 | 0.069 | 0.215 |
| **Closest to Marker** | 0.126 | 0.216 | 0.072 | 0 | 0.085 |
| **Closest to Gene** | 0.091 | 0.478 | 0.220 | 0.082 | 0 |

Table S6: Bonferroni adjusted p-values from pairwise Chi-Squared tests comparing the distribution of Hi-C scores of the five fine-mapping strategies across ten quantile bins. "Most Likely" and "Least Likely" refer to INFIMA most and least likely predictions, respectively.

| Strategy | Most Likely | Least Likely | Random | Closest to Marker |
|---|---|---|---|---|
| **Least Likely** | 1.16e-201 | | | |
| **Random** | 1.19e-86 | 2.61e-23 | | |
| **Closest to Marker** | 1.31e-39 | 3.36e-72 | 2.92e-12 | |
| **Closest to Gene** | 3.08e-17 | 6.12e-160 | 1.71e-67 | 1.35e-17 |

### 3 Supplementary Figures

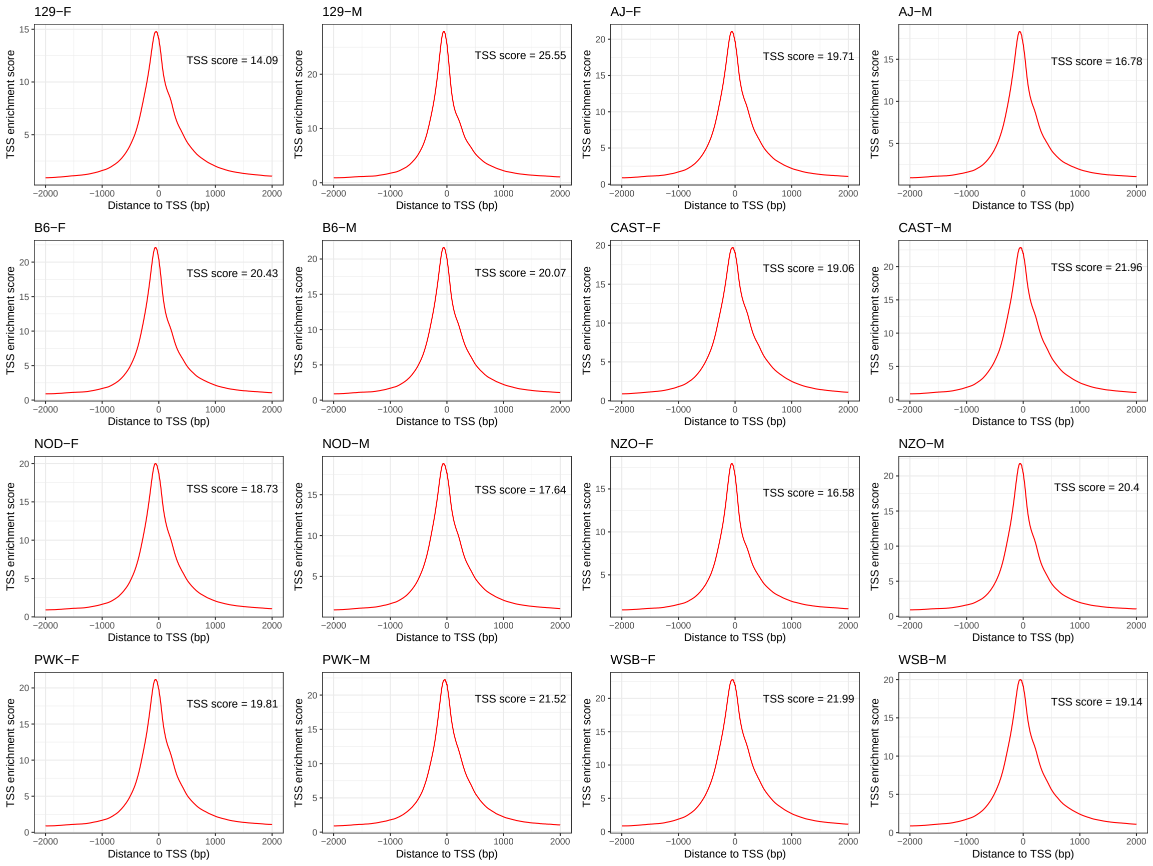

Fig. S1: Summary of transcription start site (TSS) enrichment analysis. TSS scores are calculated by ataqv (`https://github.com/ParkerLab/ataqv`) and represent an enrichment metric based on the transposition activity around the transcription start sites. Scores of 15 or larger are deemed as ideal quality.

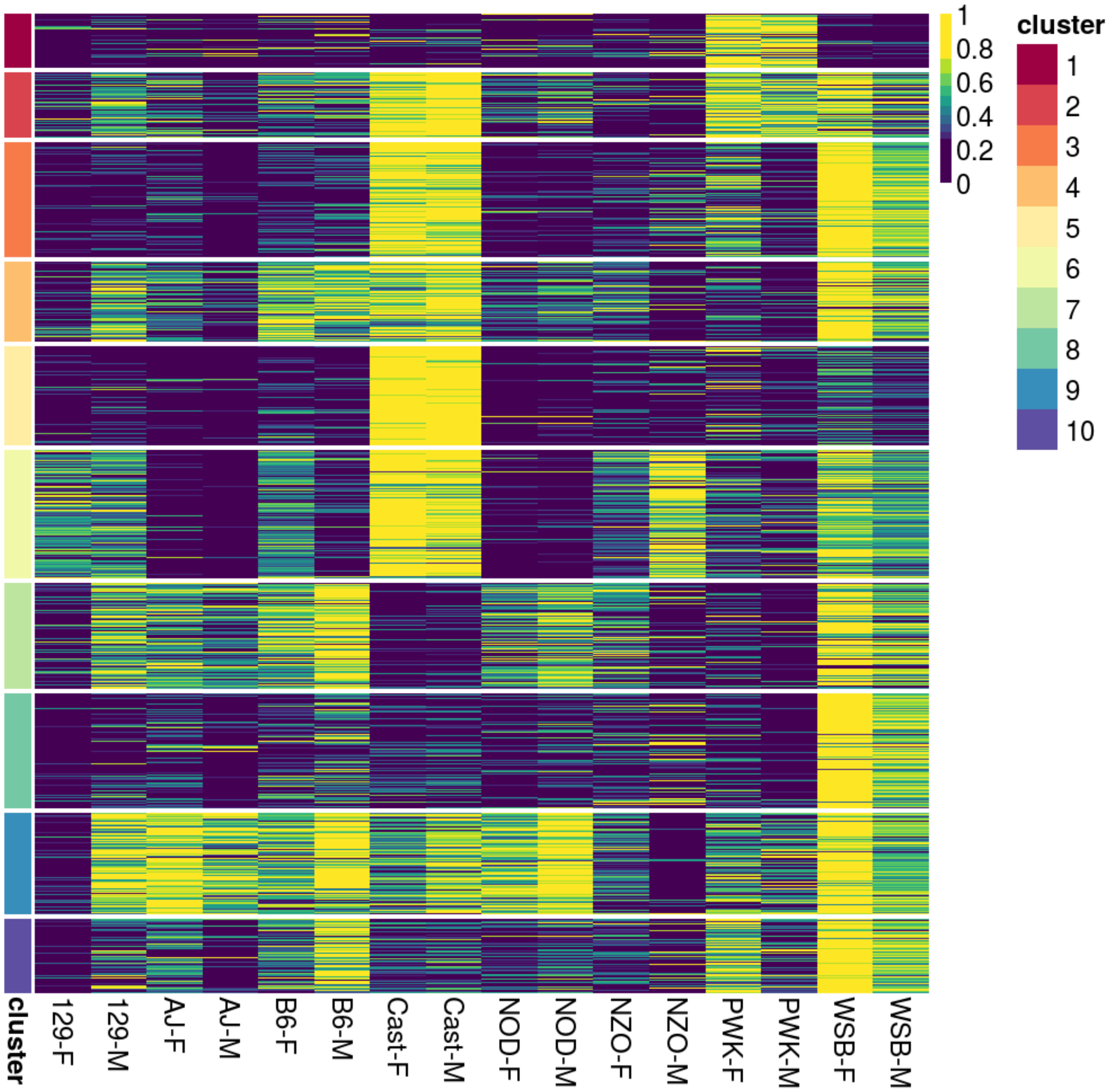

Fig. S2: The full version of Fig. 2e.

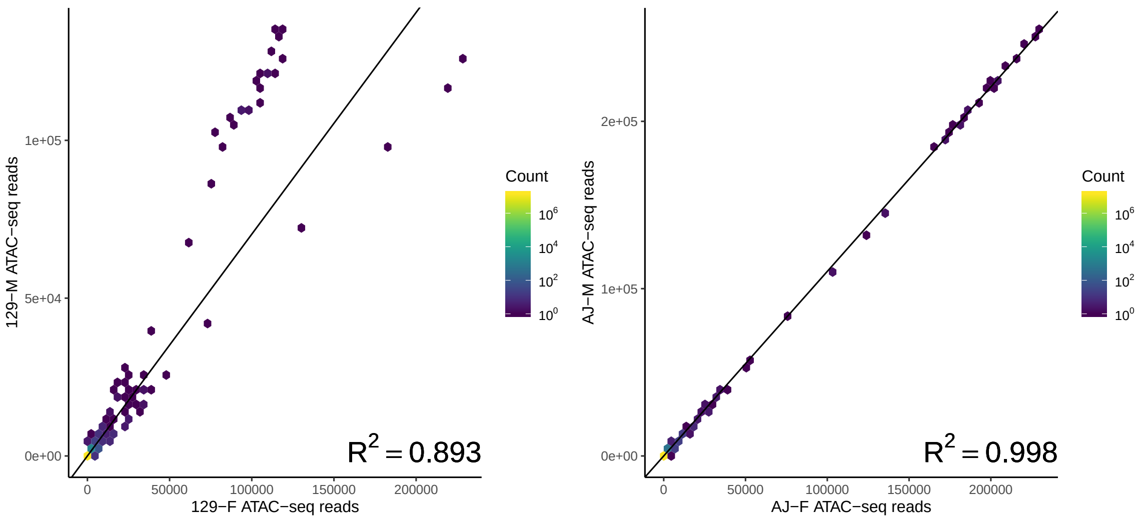

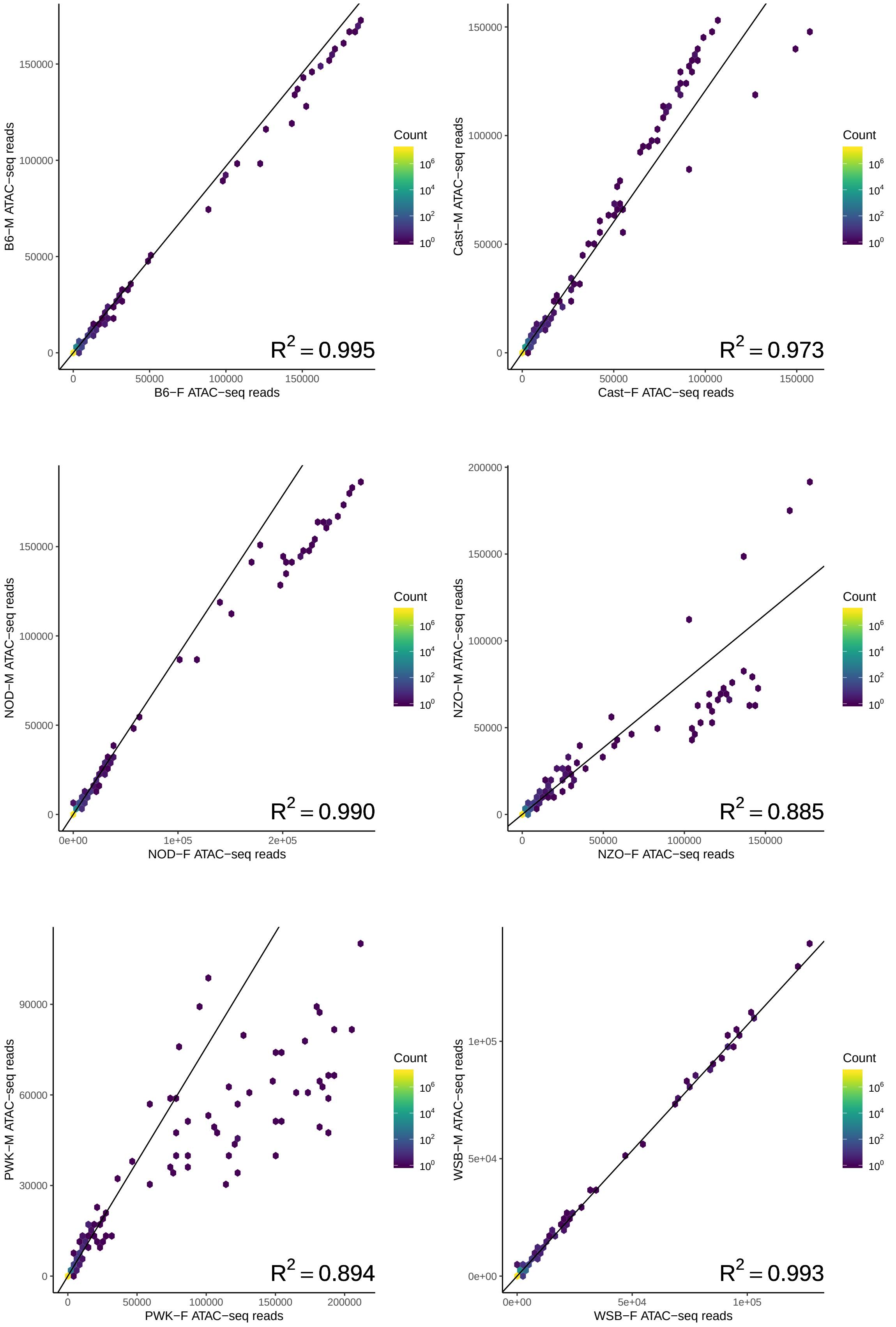

Fig. S3: Reproducibility of ATAC-seq samples. ATAC-seq read counts summarized over 200 bp bins along the genome across the male (M) and female (F) samples.

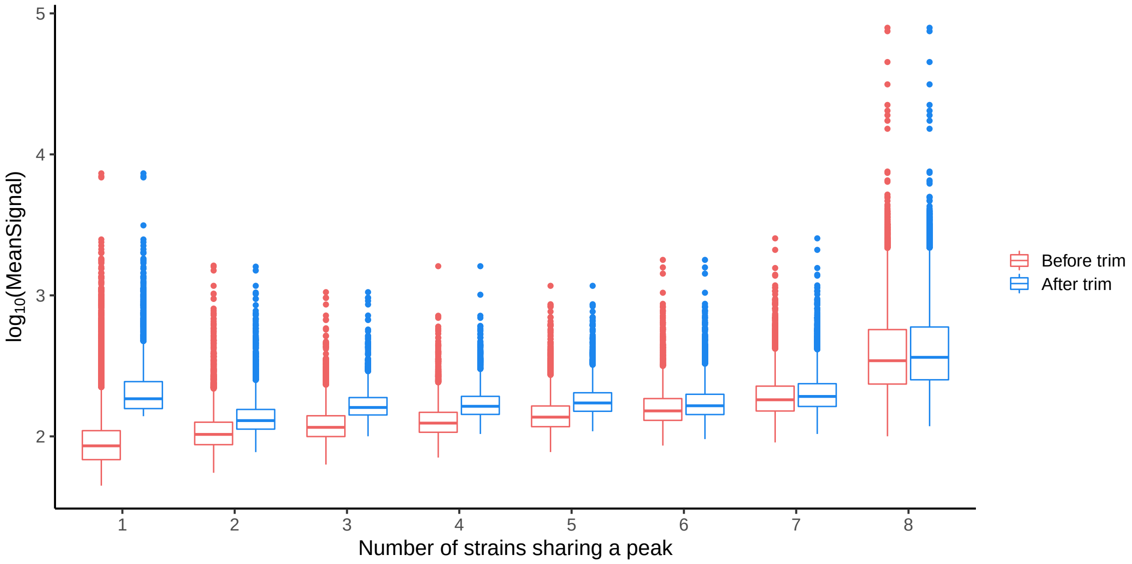

Fig. S4: ATAC-seq signal of peaks shared by different numbers of strains. ”Before trim” and ”After trim” denote ATAC-seq signal of the master peaks before and after trimming, respectively, as a function of the total number of strains a peak is shared by.

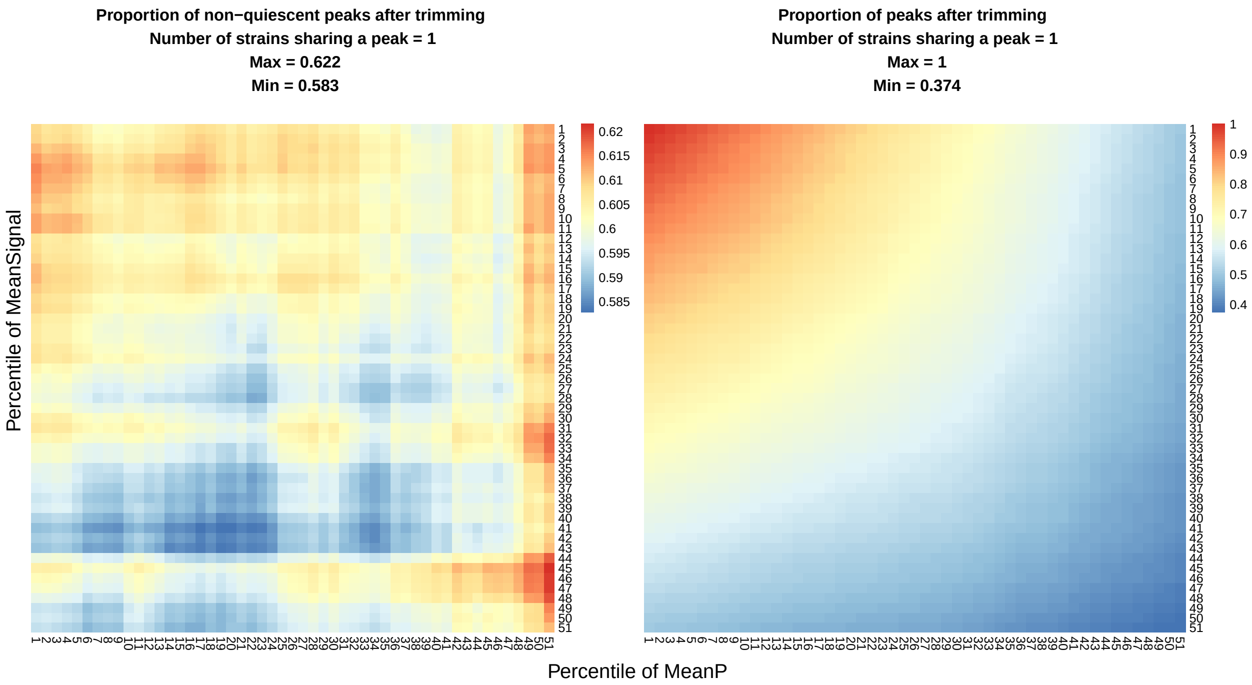

**Proportion of non−quiescent peaks after trimming**
**Number of strains sharing a peak = 2**
**Max = 0.735**
**Min = 0.692**

**Proportion of peaks after trimming**
**Number of strains sharing a peak = 2**
**Max = 1**
**Min = 0.403**

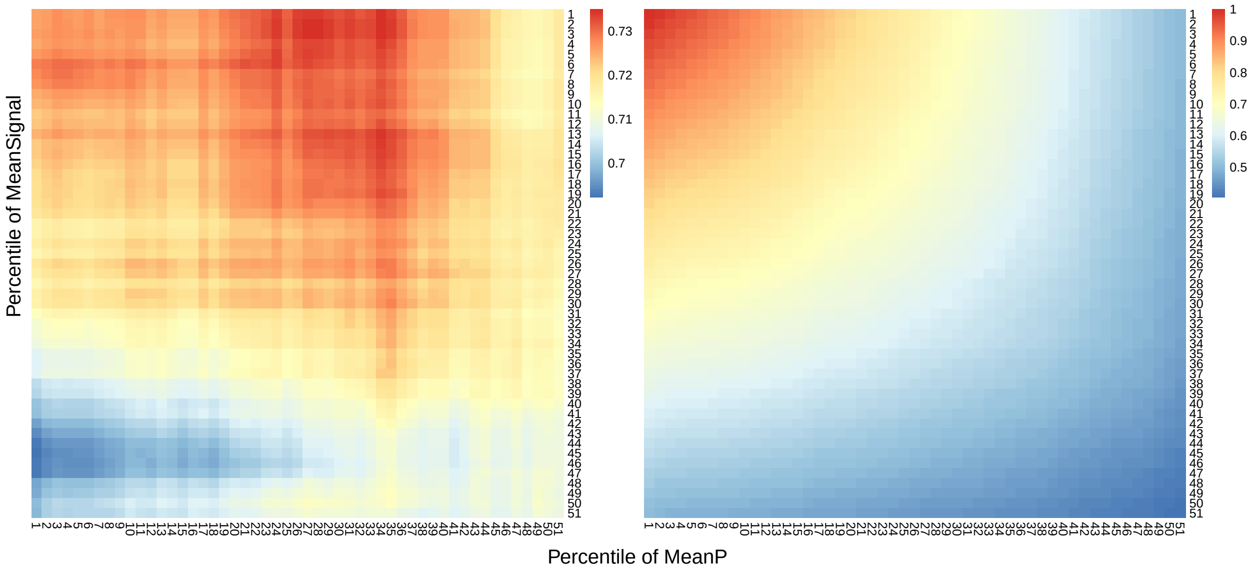

**Proportion of non−quiescent peaks after trimming**
**Number of strains sharing a peak = 3**
**Max = 0.778**
**Min = 0.752**

**Proportion of peaks after trimming**
**Number of strains sharing a peak = 3**
**Max = 1**
**Min = 0.402**

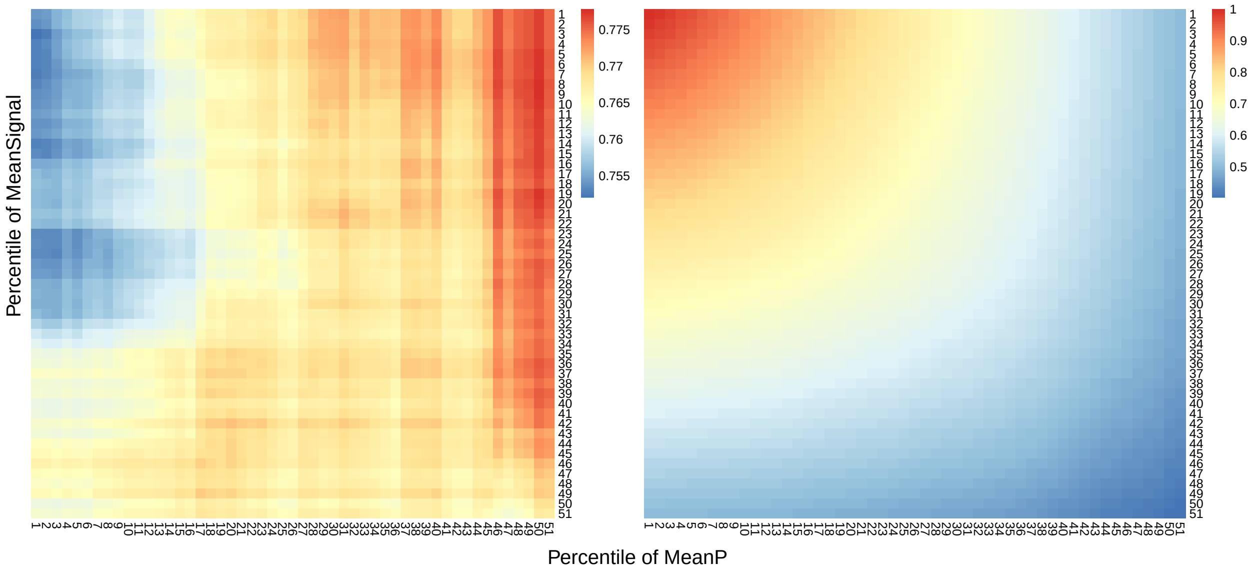

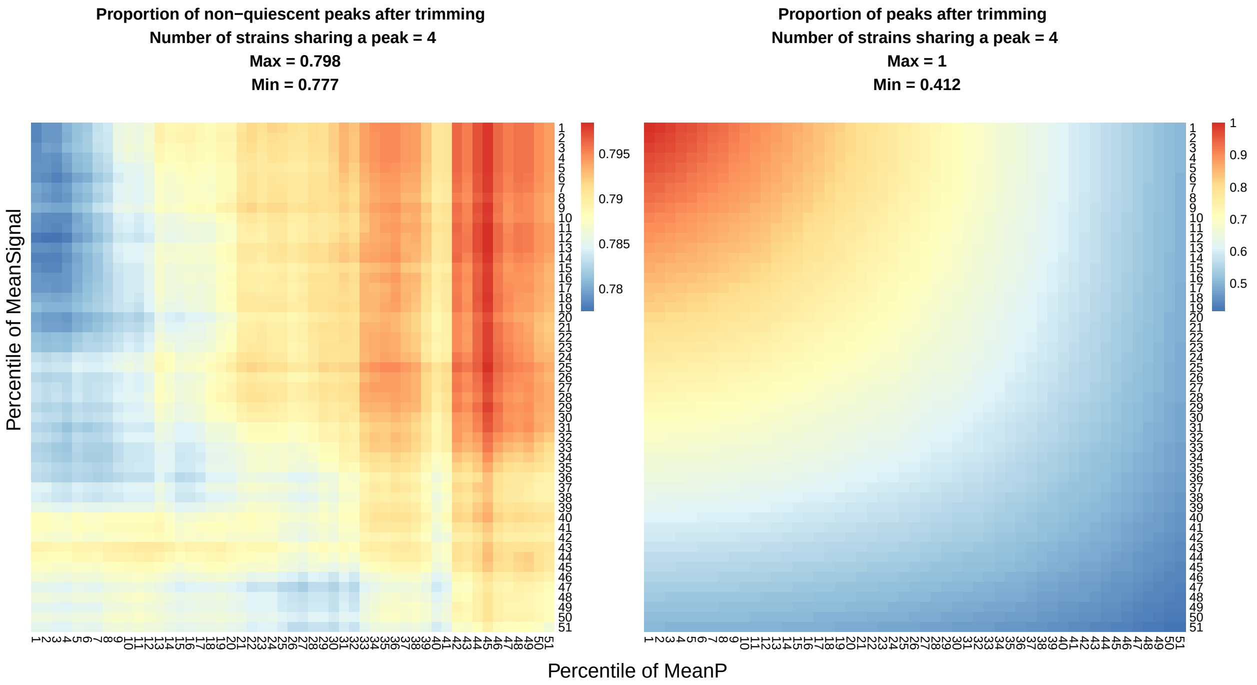
Proportion of non−quiescent peaks after trimming
Number of strains sharing a peak = 4
Max = 0.798
Min = 0.777
0.795
0.79
0.785
0.78
Proportion of peaks after trimming
Number of strains sharing a peak = 4
Max = 1
Min = 0.412
1
0.9
0.8
0.7
0.6
0.5
Percentile of MeanSignal
Percentile of MeanP

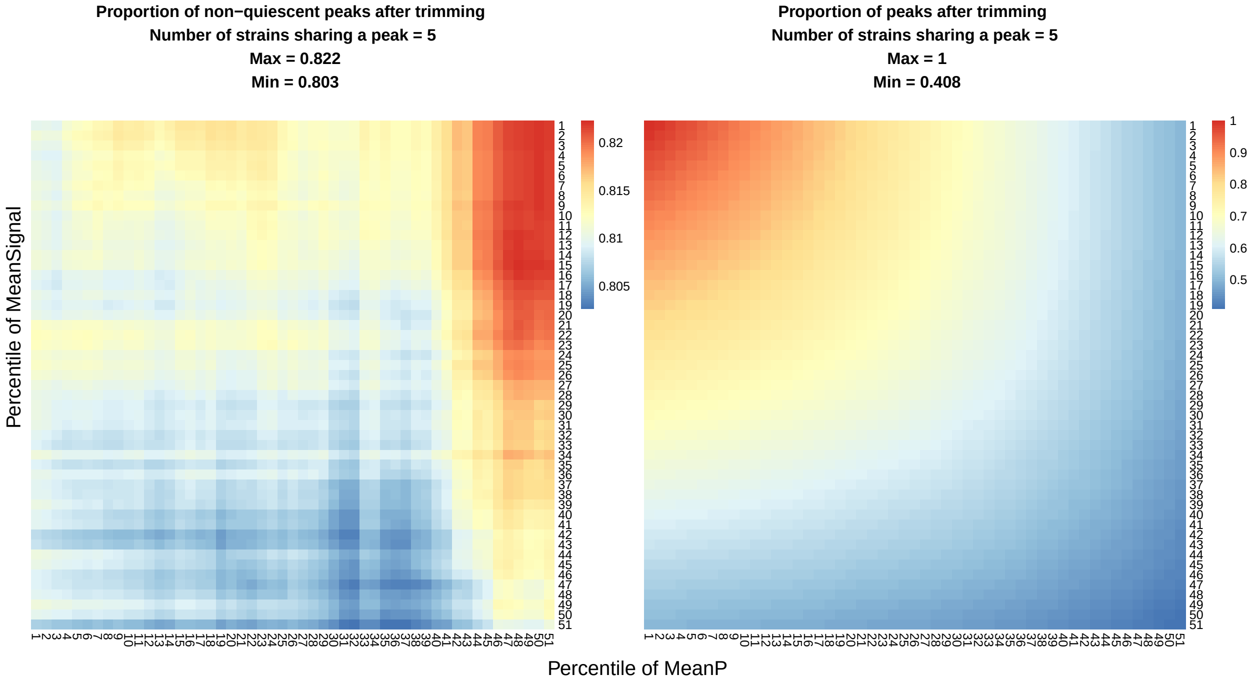
Proportion of non−quiescent peaks after trimming
Number of strains sharing a peak = 5
Max = 0.822
Min = 0.803
0.82
0.815
0.81
0.805
Proportion of peaks after trimming
Number of strains sharing a peak = 5
Max = 1
Min = 0.408
1
0.9
0.8
0.7
0.6
0.5
Percentile of MeanSignal
Percentile of MeanP

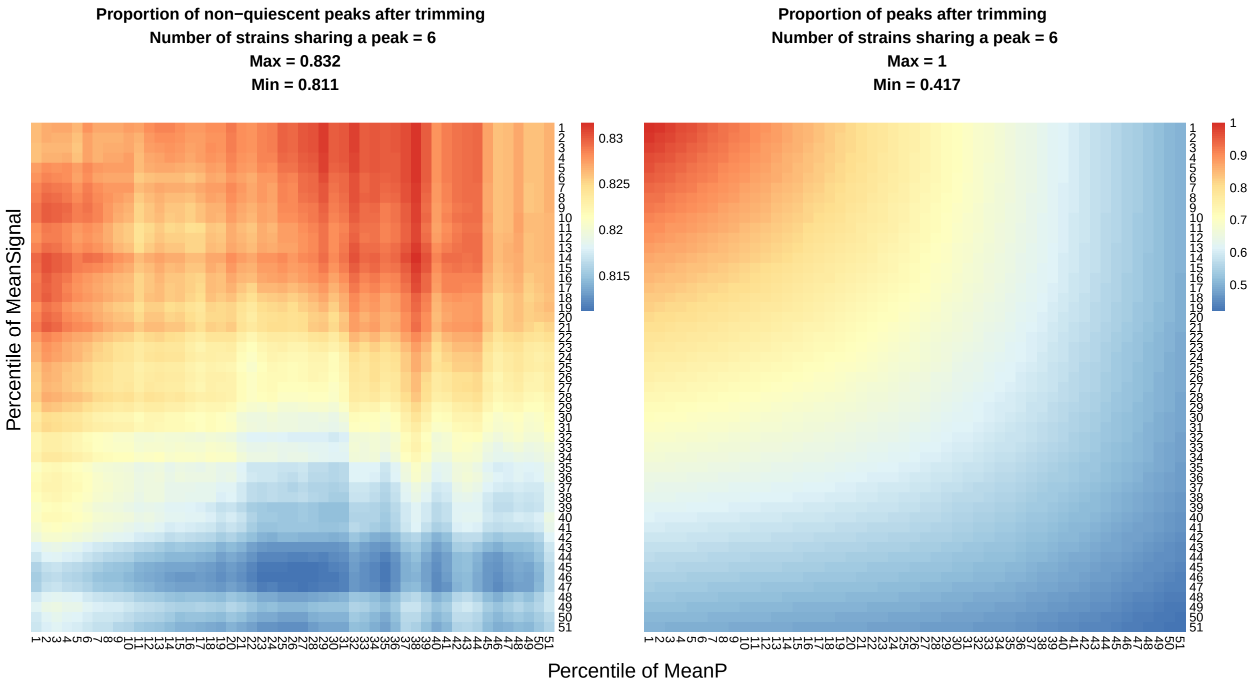
Proportion of non−quiescent peaks after trimming
Number of strains sharing a peak = 6
Max = 0.832
Min = 0.811
0.83
0.825
0.82
0.815
Percentile of MeanSignal
Proportion of peaks after trimming
Number of strains sharing a peak = 6
Max = 1
Min = 0.417
1
0.9
0.8
0.7
0.6
0.5
Percentile of MeanP

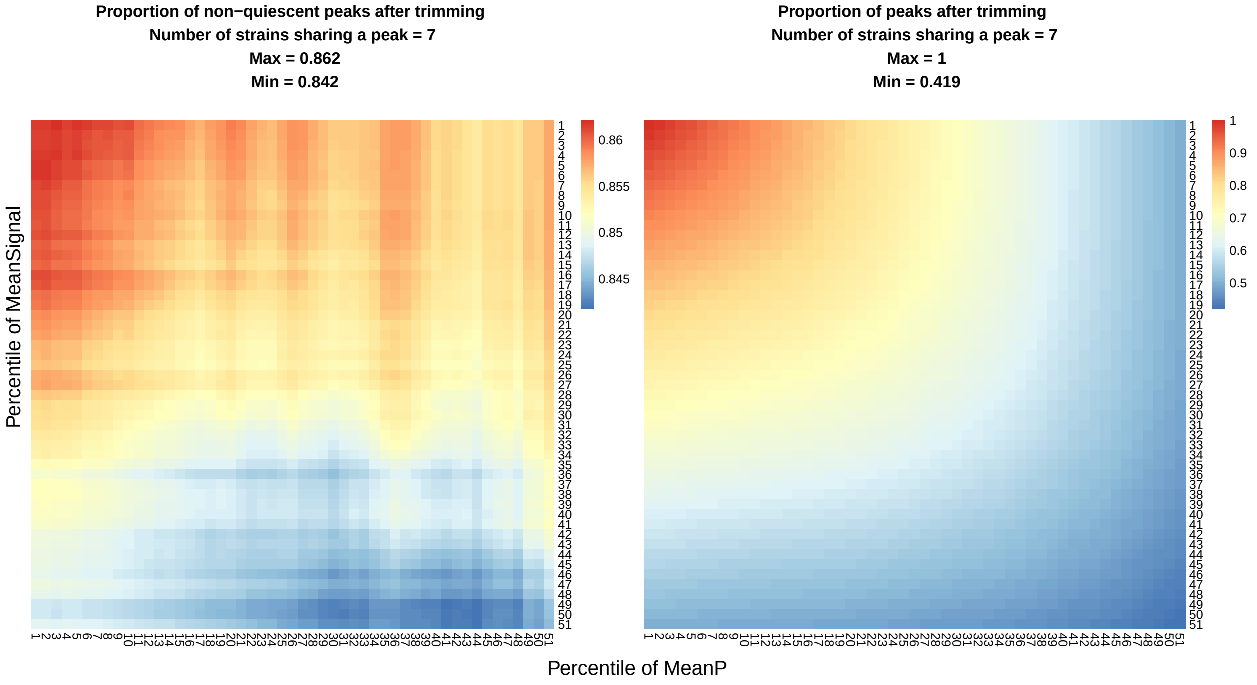
Proportion of non−quiescent peaks after trimming
Number of strains sharing a peak = 7
Max = 0.862
Min = 0.842
0.86
0.855
0.85
0.845
Percentile of MeanSignal
Proportion of peaks after trimming
Number of strains sharing a peak = 7
Max = 1
Min = 0.419
1
0.9
0.8
0.7
0.6
0.5
Percentile of MeanP

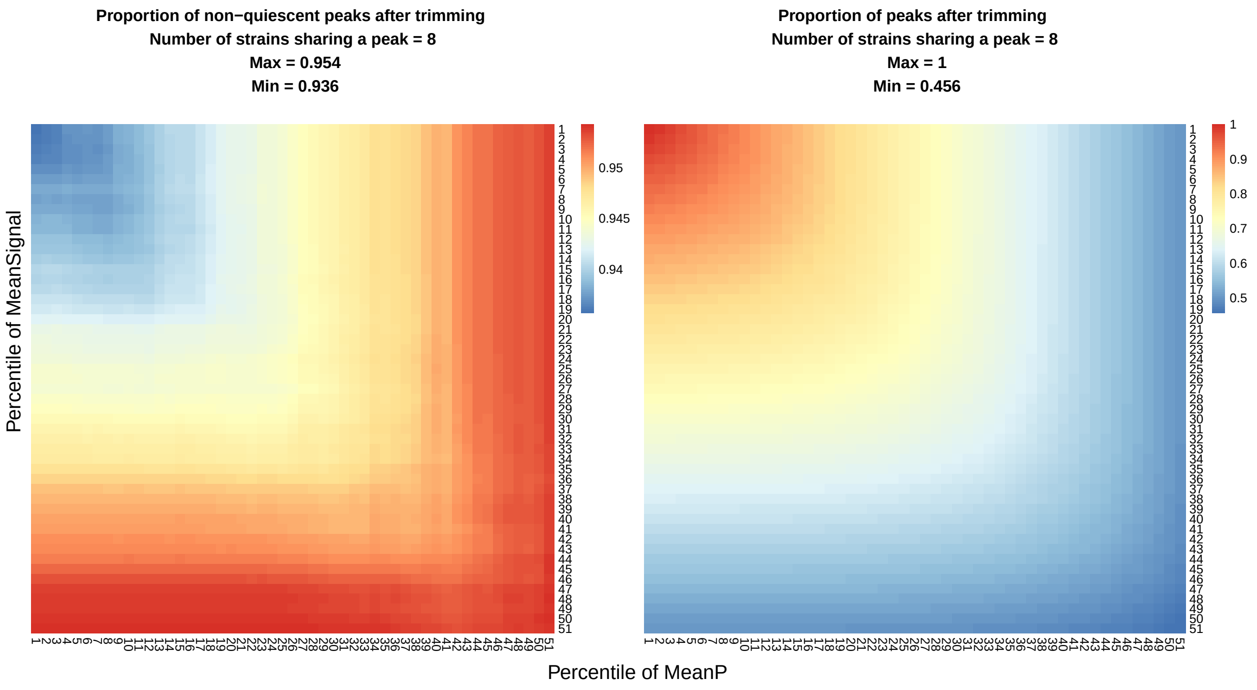

Fig. S5: Trimming of the master peak list. Heatmaps depict the proportion of retained peaks (Right columns) and the proportion of peaks in non-quiescent regions of the genome pooled across a set of tissues (Left columns) as a function of the two trimming parameters for each value of the number of strains sharing the peaks. The final percentiles for the tuning parameters (MeanP, MeanSignal) were set as: (51, 45), (23, 6), (46, 1), (45, 0), (47, 0), (2, 14), (0, 0), (0,0) for the number of strains sharing the peaks = 1, 2, 3, 4, 5, 6, 7, 8, respectively. Here, ”(MeanP, MeanSignal) = (a, b)” corresponds to removing peaks of the master peak list with the lowest (a-1)% of ‘MeanP’ and/or the lowest (b-1)% of ‘MeanSignal’.

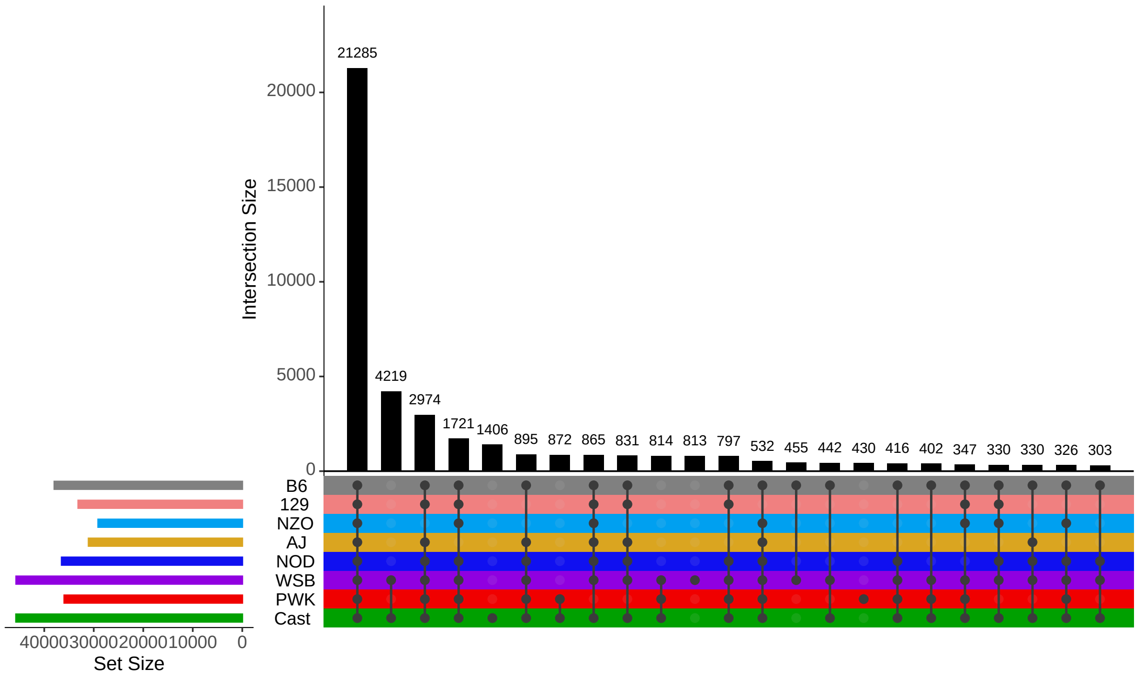

Fig. S6: The UpSet plot for 51,014 ATAC-seq master peaks after quality trimming. Bars represent the numbers of peaks that originate from different combinations of strains.

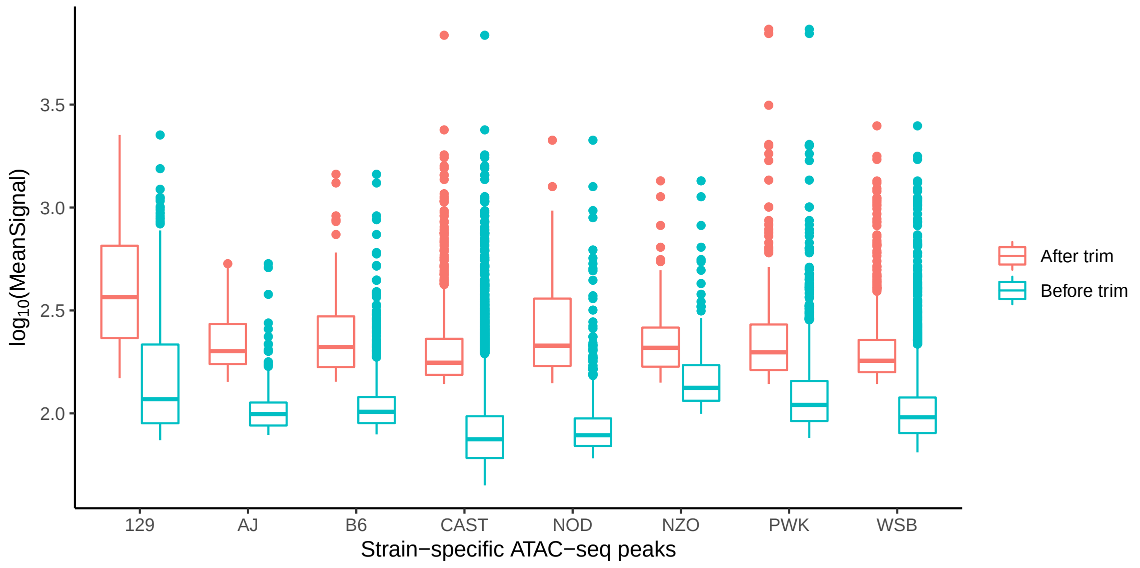

Fig. S7: ATAC-seq signal of strain-specific peaks. "Before trim" and "After trim" denote ATAC-seq signal of the master peaks before and after trimming, respectively.

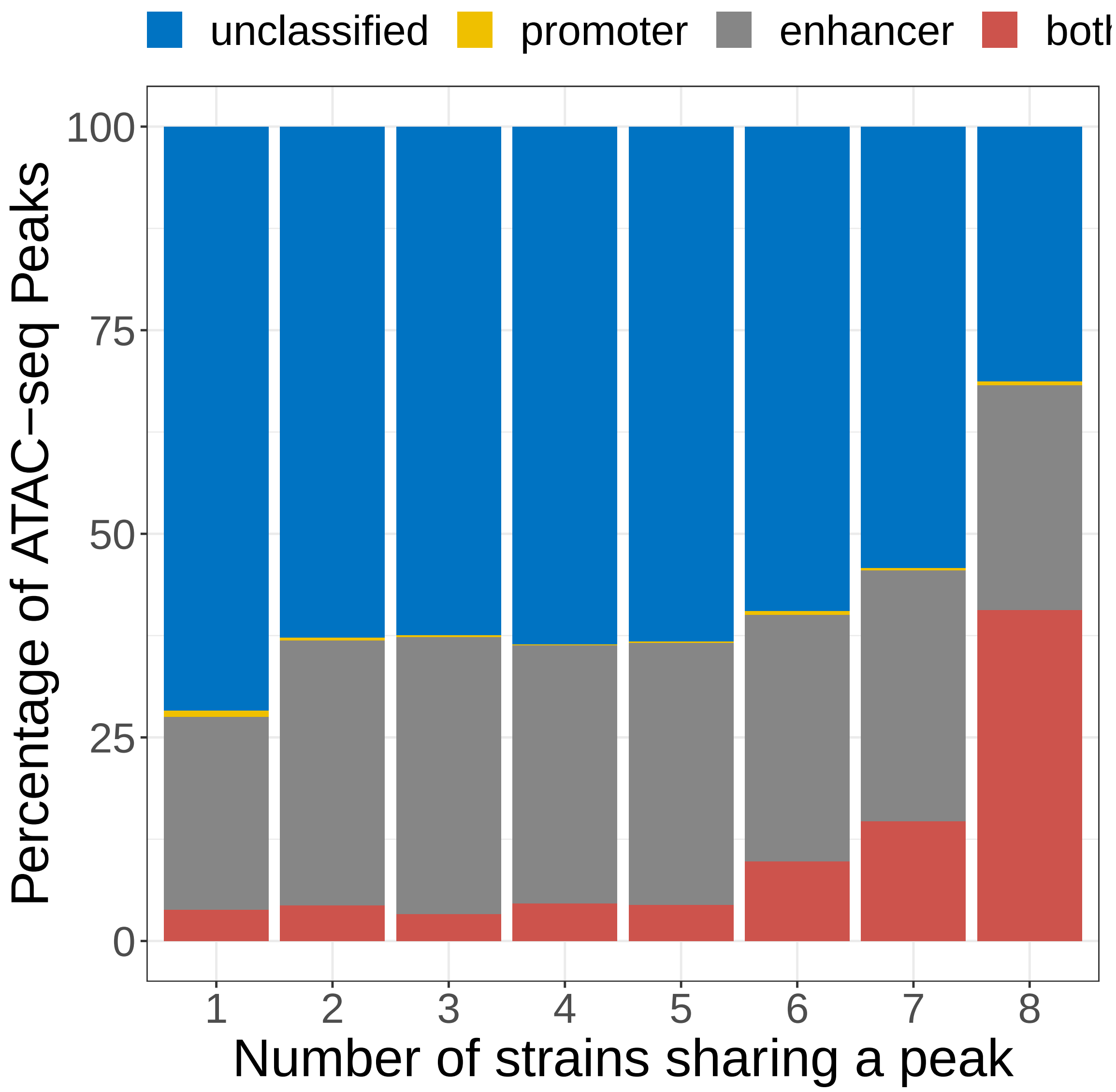

Fig. S8: Promoter/enhancer annotation of the ATAC-seq master peak list stratified by the numbers of strains sharing a peak (see URLs; Supplementary Notes).

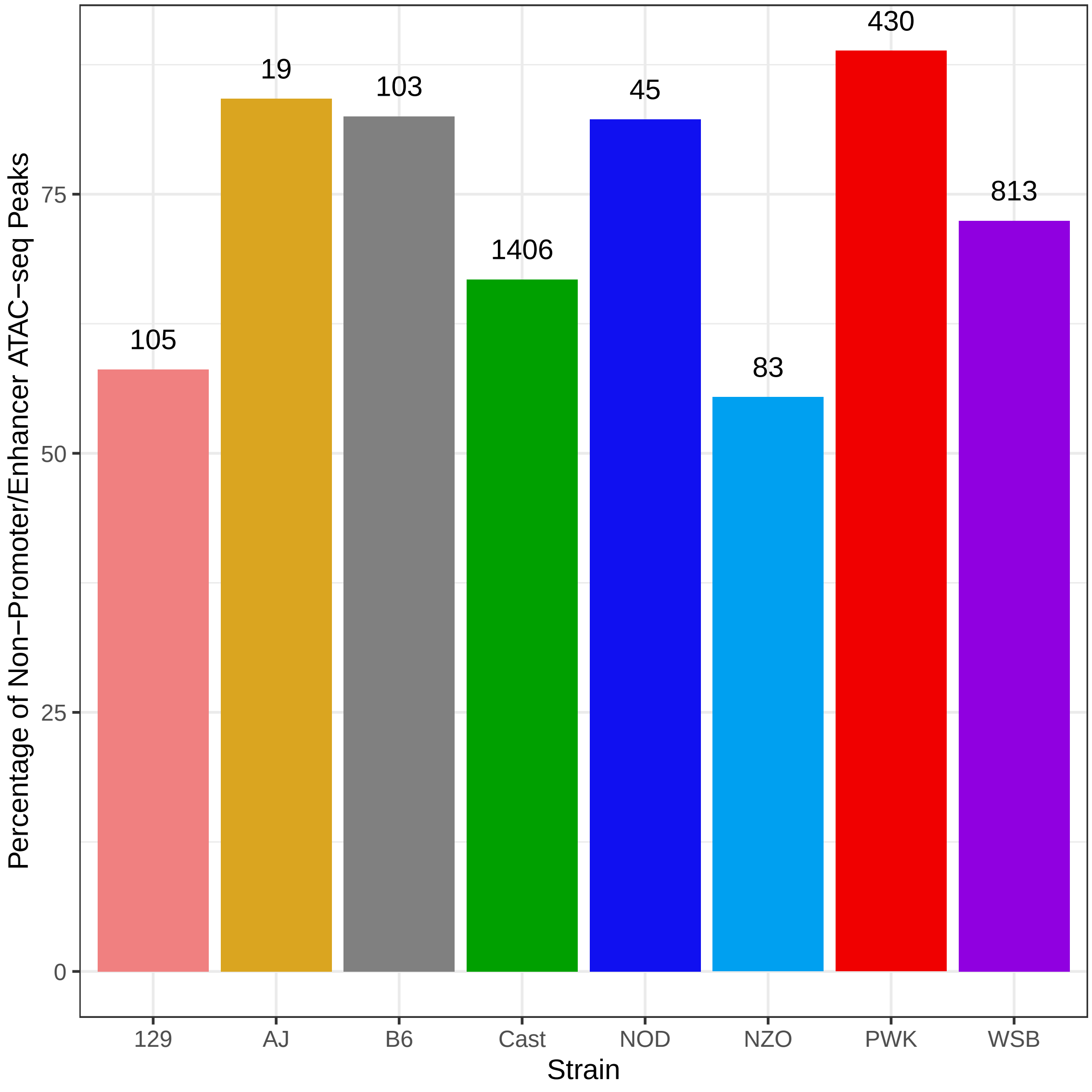

Fig. S9: Percentage of strain-specific ATAC-seq peaks not categorized as promoter or enhancer. The total numbers of strain-specifc ATAC-seq peaks are displayed on top of each bar.

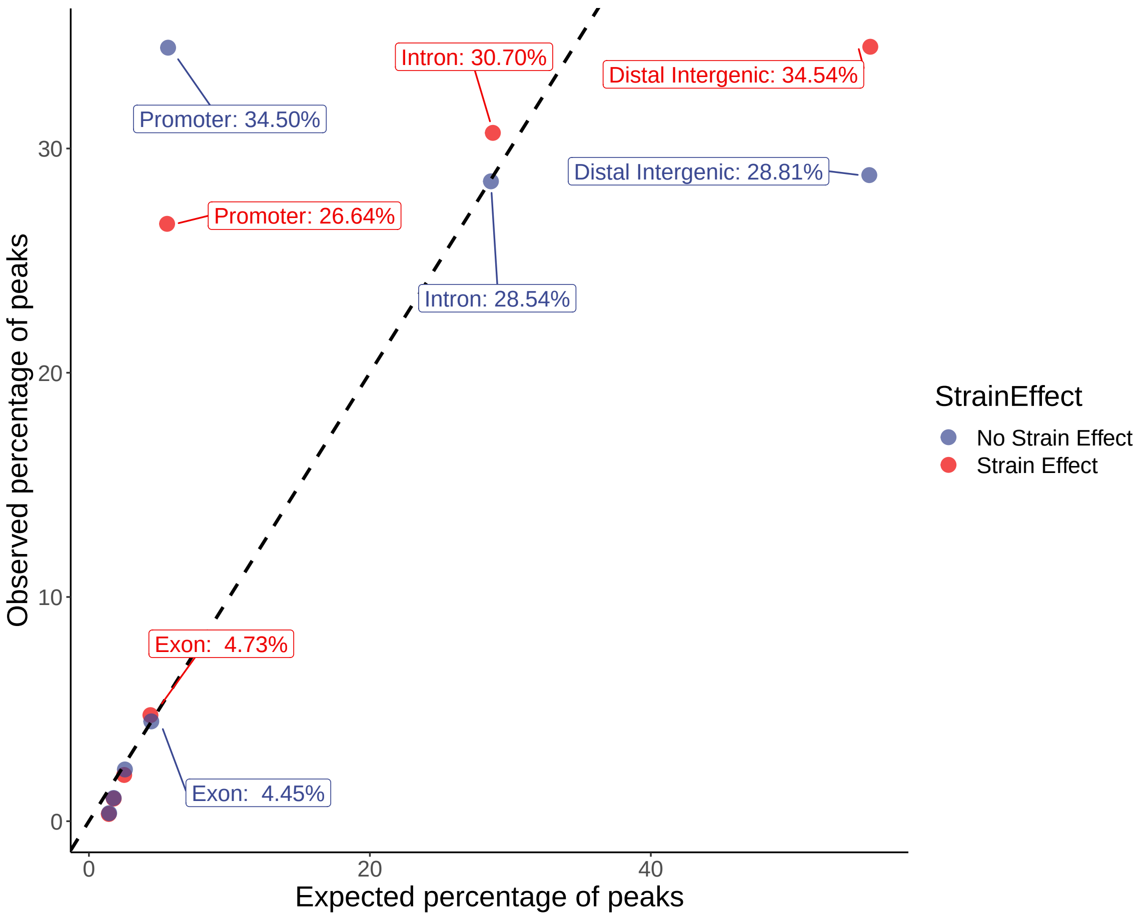

Fig. S10: Genomic location analysis of the ATAC-seq peaks with or without strain effect. Expected percentages are computed with the regioneR package using genomic location annotations from ChIPseeker package.

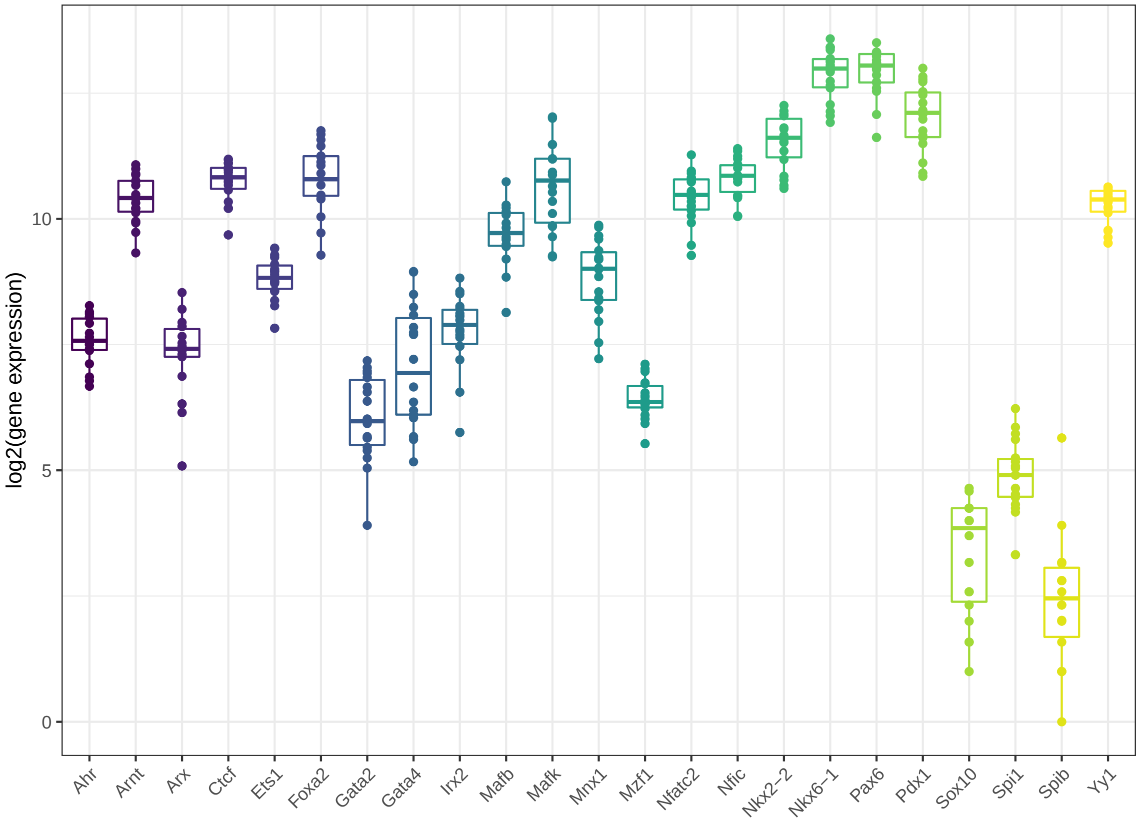

Fig. S11: Founder islet expressions of TFs in Fig. 3a. TFs that are not shown did not have detectable levels of expression.

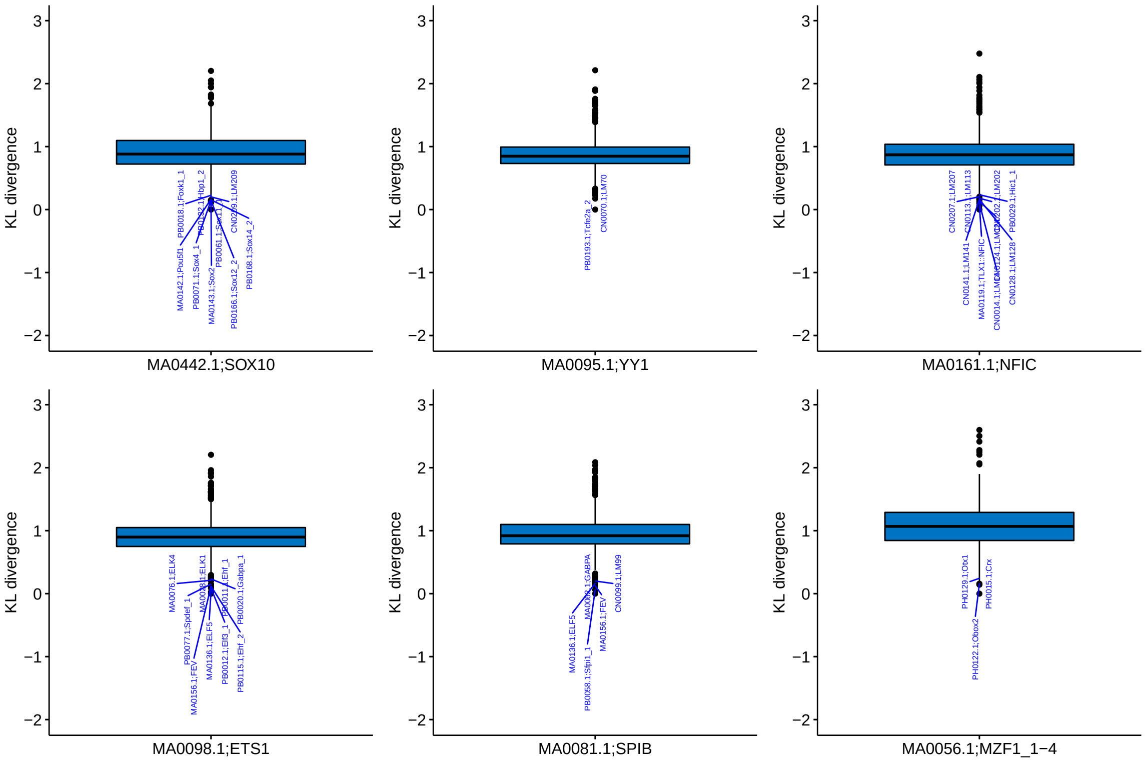

Fig. S12: Kullback-Leibler (KL) divergence of highlighted TF motifs with footprints in Fig. 3a to all other 744 TF motifs queried. Similarities between motifs were computed based on both Kullback-Leibler divergence and Pearson correlation coefficient. After taking intersection among top 20 with respect to the two similarity metrics, top 10 motifs with KL less than 0.25 are labelled.

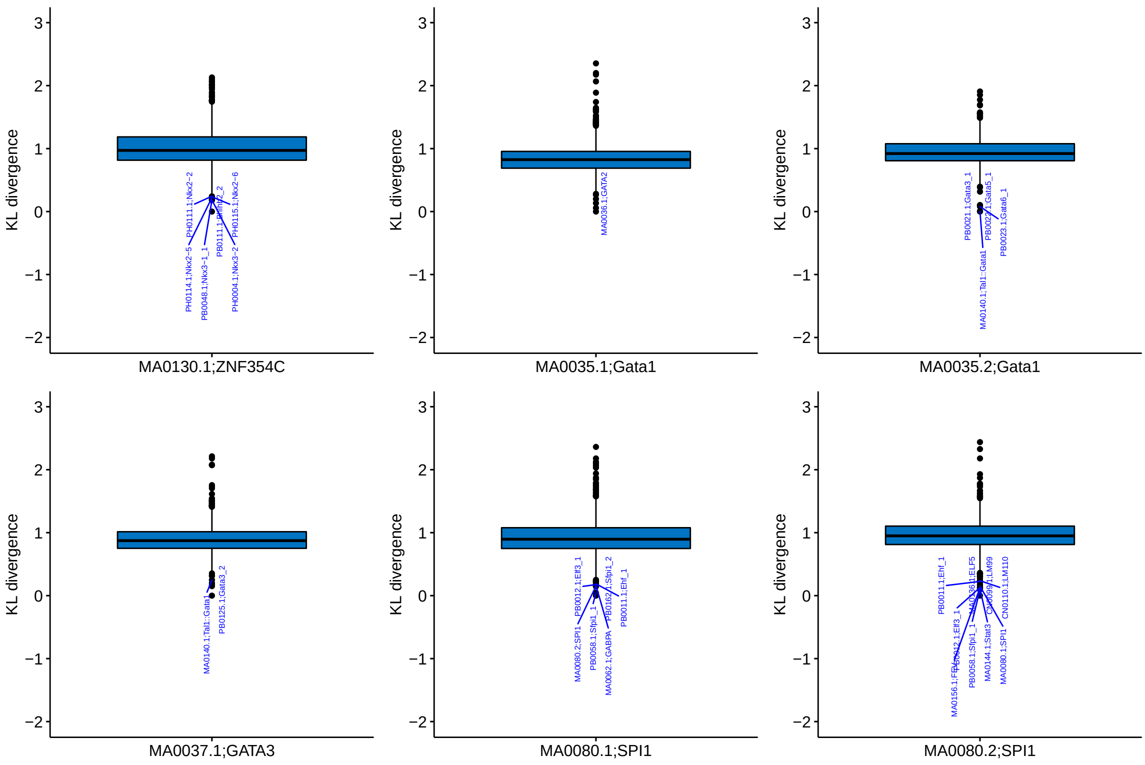

Fig. S13: Kullback-Leibler (KL) divergence of highlighted TF motifs with footprints in Fig. 3a to all other 744 TF motifs queried. Similarities between motifs were computed based on both Kullback-Leibler divergence and Pearson correlation coefficient. After taking intersection among top 20 with respect to the two similarity metrics, top 10 motifs with KL less than 0.25 are labelled.

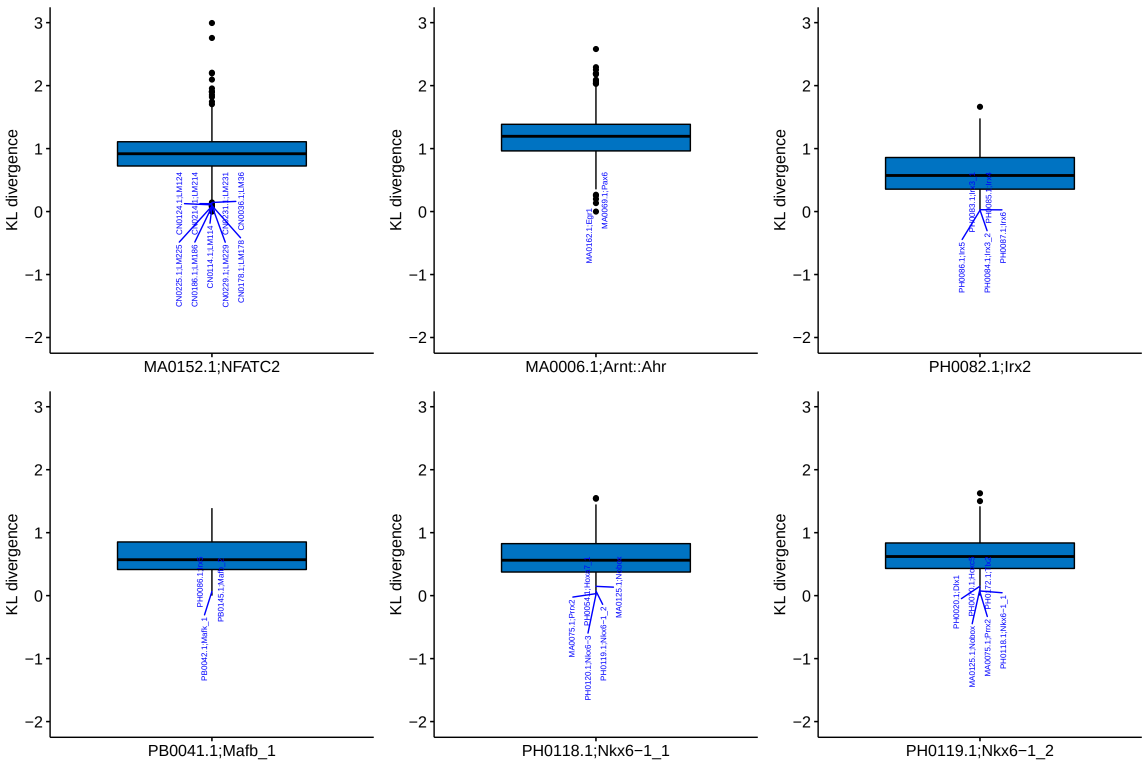

Fig. S14: Kullback-Leibler (KL) divergence of highlighted TF motifs with footprints in Fig. 3a to all other 744 TF motifs queried. Similarities between motifs were computed based on both Kullback-Leibler divergence and Pearson correlation coefficient. After taking intersection among top 20 with respect to the two similarity metrics, top 10 motifs with KL less than 0.25 are labelled.

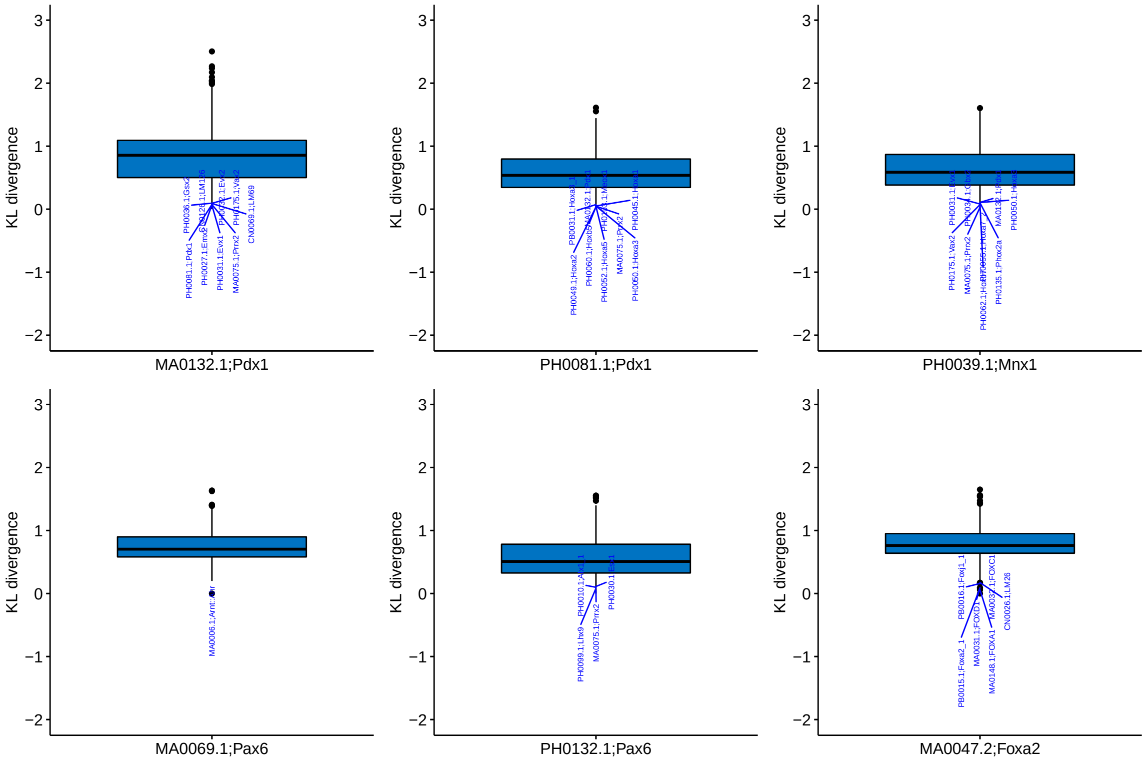

Fig. S15: Kullback-Leibler (KL) divergence of highlighted TF motifs with footprints in Fig. 3a to all other 744 TF motifs queried. Similarities between motifs were computed based on both Kullback-Leibler divergence and Pearson correlation coefficient. After taking intersection among top 20 with respect to the two similarity metrics, top 10 motifs with KL less than 0.25 are labelled.

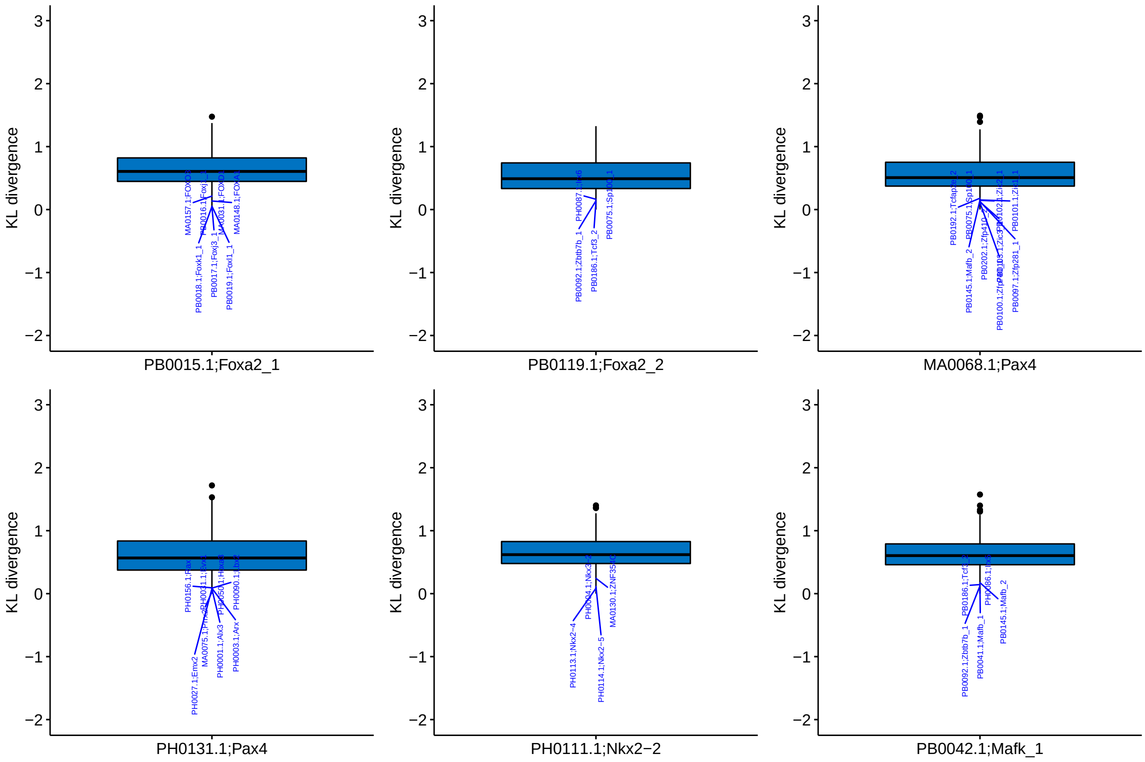

Fig. S16: Kullback-Leibler (KL) divergence of highlighted TF motifs with footprints in Fig. 3a to all other 744 TF motifs queried. Similarities between motifs were computed based on both Kullback-Leibler divergence and Pearson correlation coefficient. After taking intersection among top 20 with respect to the two similarity metrics, top 10 motifs with KL less than 0.25 are labelled.

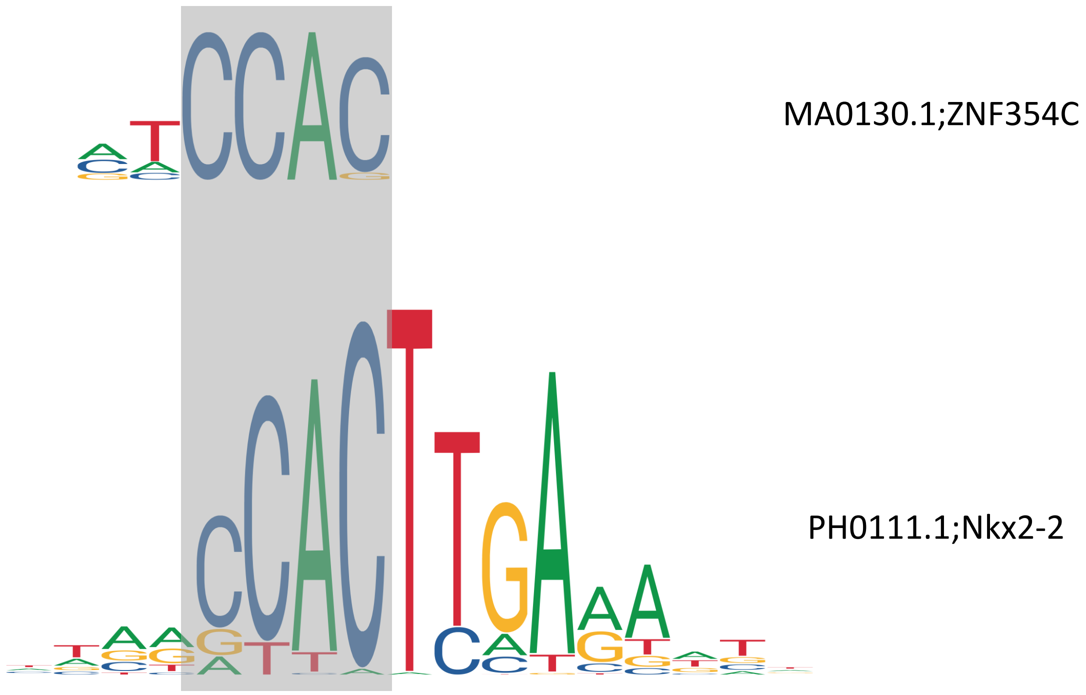

Fig. S17: Sequence logos of ZNF354C and Nkx2-2.

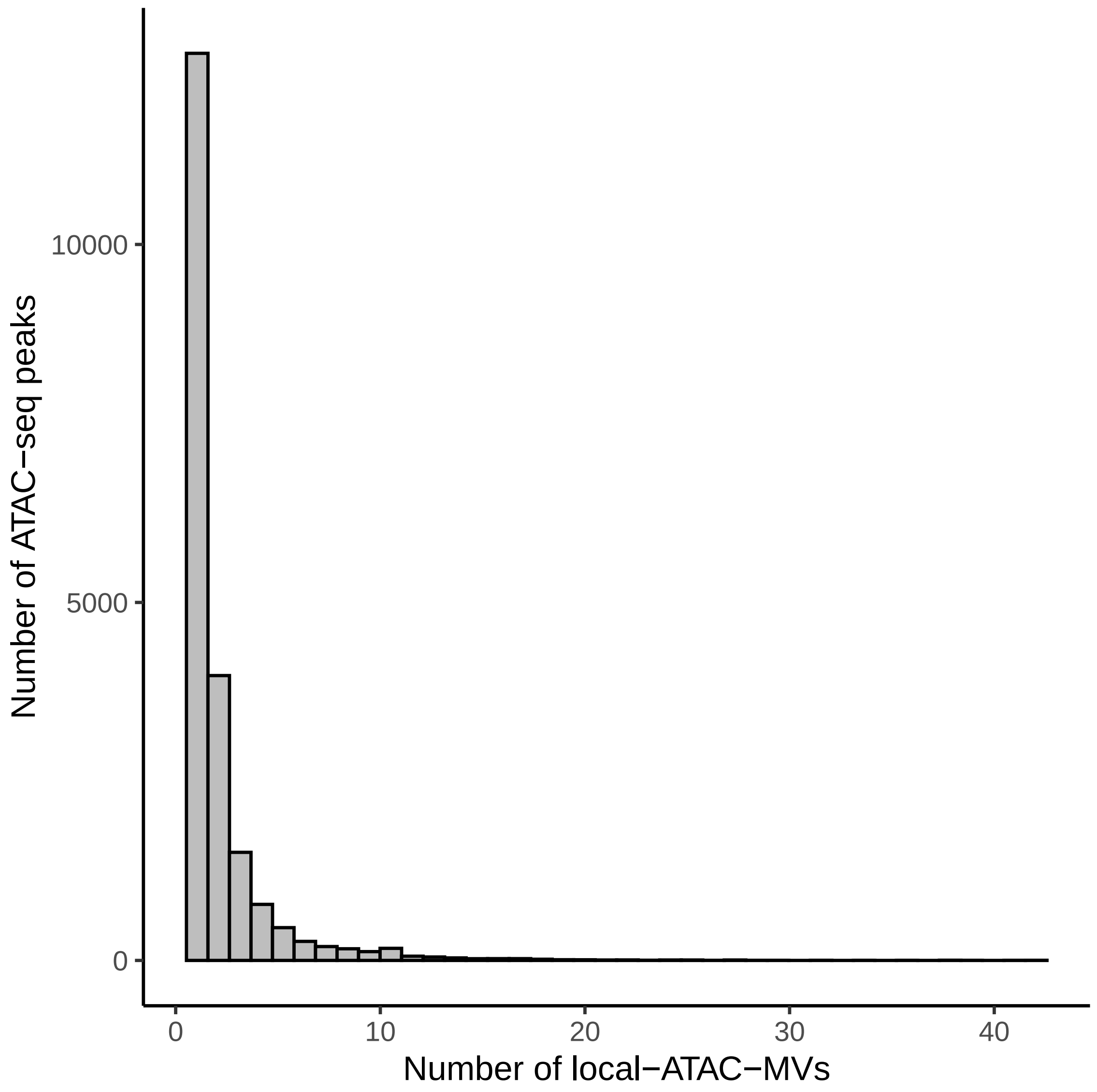

Fig. S18: Histogram of numbers of local-ATAC-MVs within a SNP harboring differential ATAC-seq master peak.

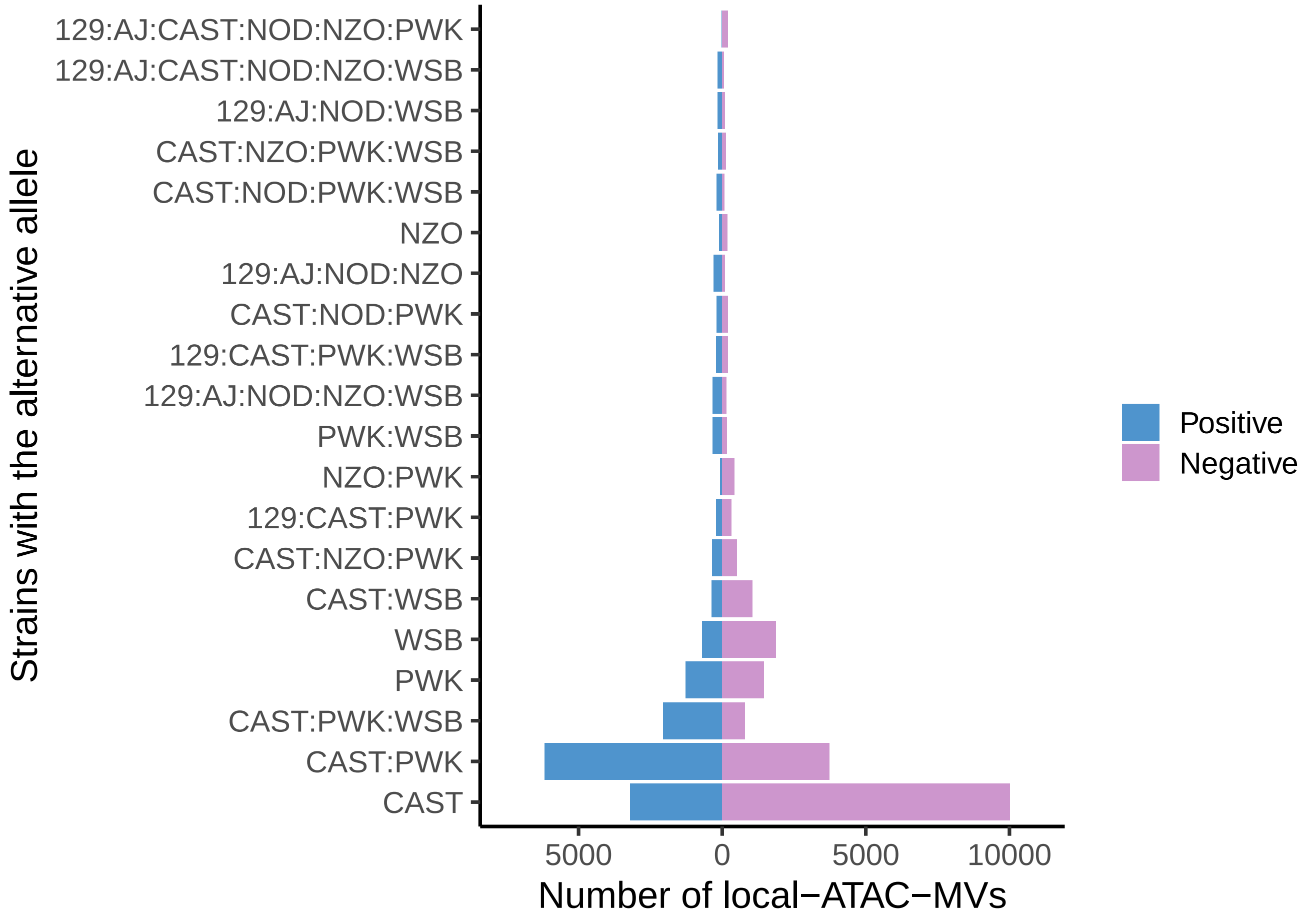

Fig. S19: Genotype decomposition of local-ATAC-MVs. Groups of strains with the alternative allele compared to reference B6 allele are displayed in the y-axis. Positive category: alternative allele is associated with an increase in chromatin accessibility. Negative category: alternative allele is associated with a decrease in chromatin accessibility.

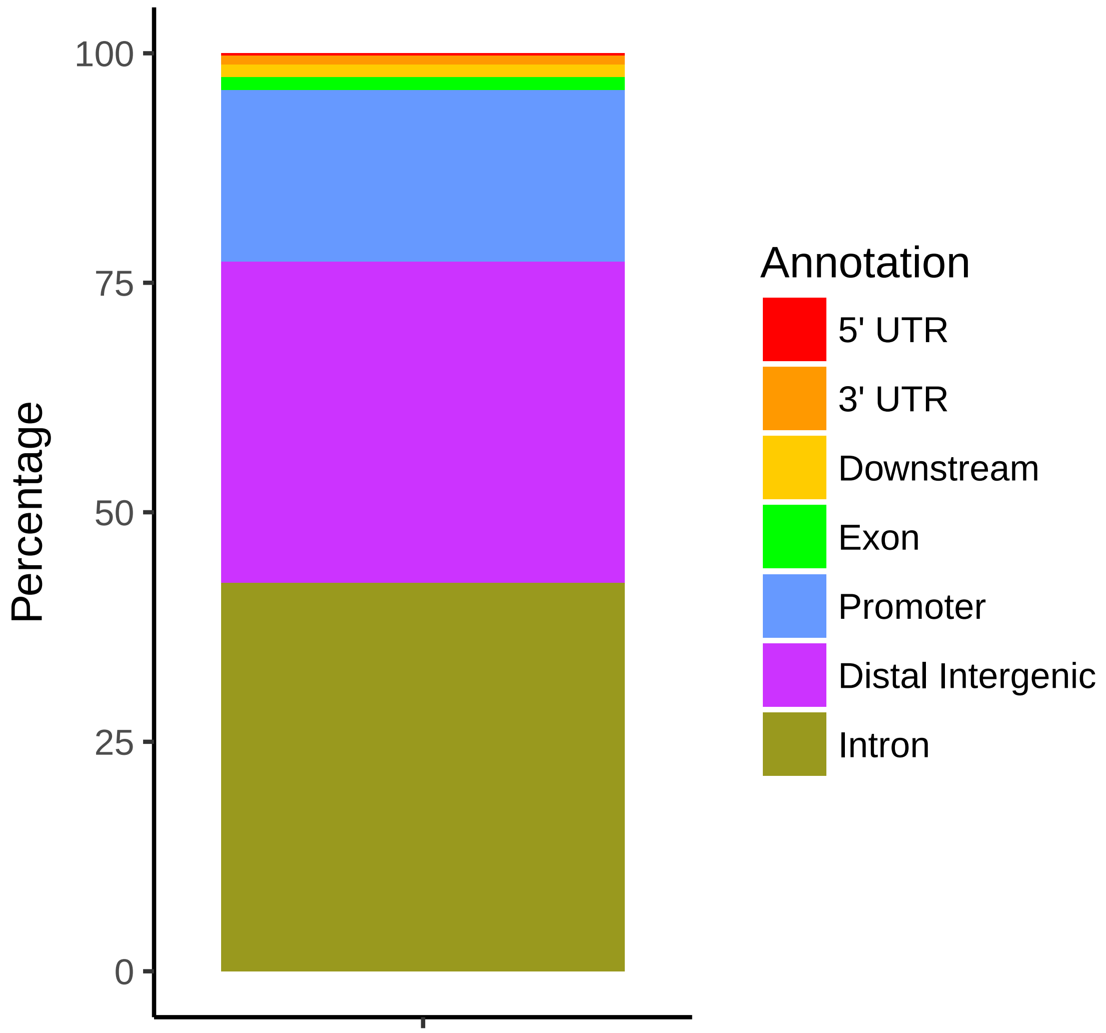

Fig. S20: Genomic location annotation of local-ATAC-MVs. The annotation categories and coordinates are from the ChIPseeker R package

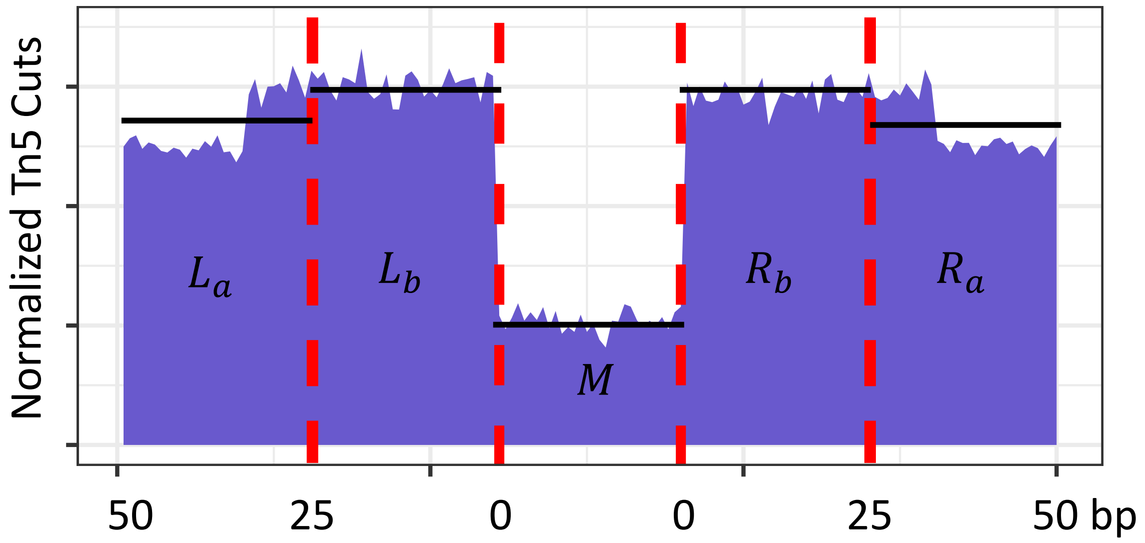

Fig. S21: Illustration of the footprint depth (FPD) calculation. y-axis represents normalized Tn5 cuts and x-axis represents distance to TF binding site in bp. $L_a$, $L_b$, $M$, $R_b$, and $R_a$ represent the average signal within regions of Left 26 - 50 bp, Left 0 - 25 bp, Middle, Right 0 - 25 bp, Right 26 - 50 bp with respect to the centered TF binding site, respectively. The black lines depict the average signal values within each of these regions. The footprint depth is defined as FPD $= 1 - \frac{M}{\max(L_a, L_b, R_b, R_a)}$. FPD values at individual motif locations are compared across strains with the reference and alternative alleles of the SNP for each SNP-motif combination.

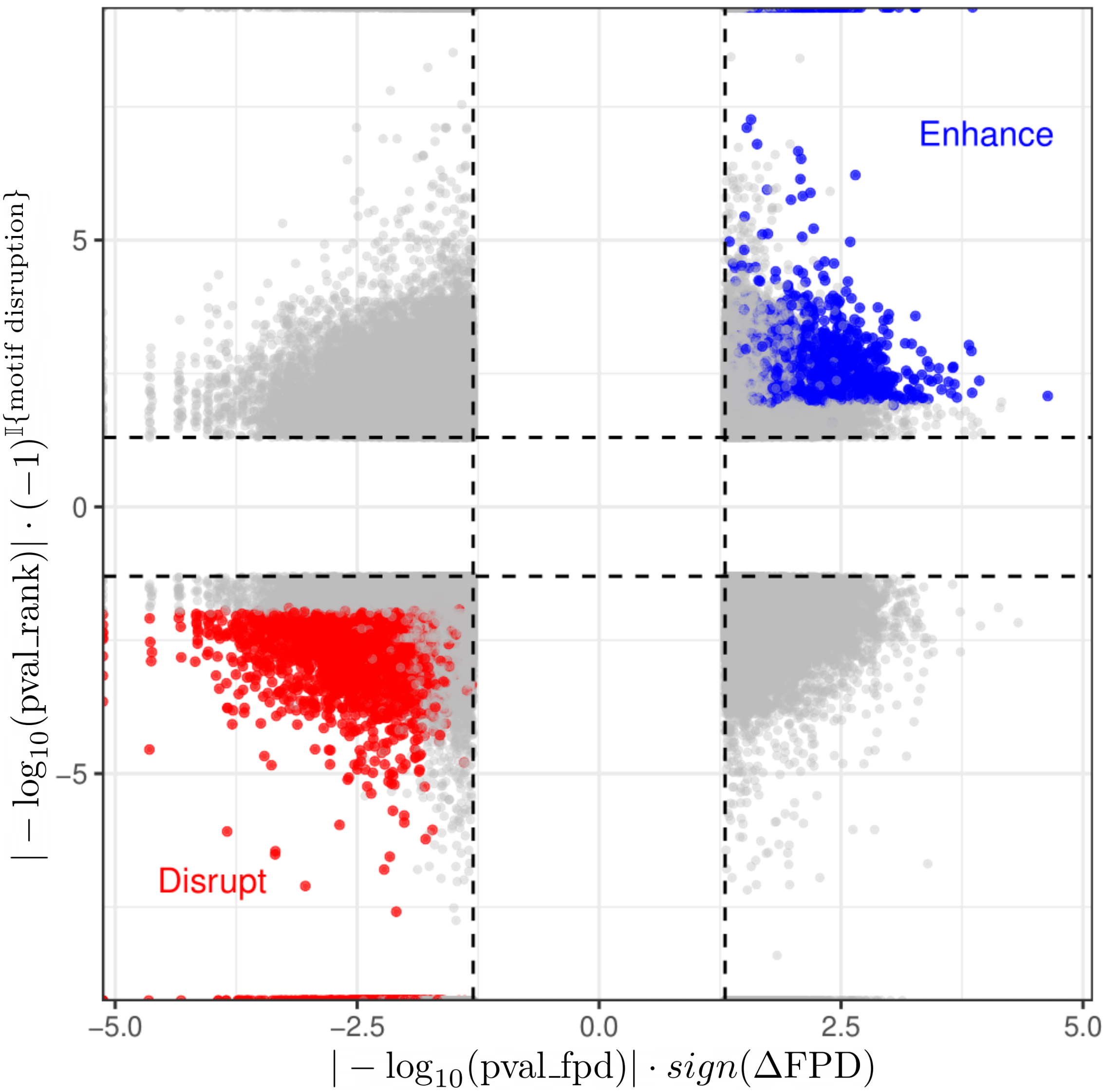

Fig. S22: Summary of the atSNP and FPD analysis for individual footprints. Only the footprints with `pval_rank` $< 0.05$ and `pval_fpd` $< 0.05$ are displayed in the figure (grey dashed lines). The sign of x-axis: negative means $\Delta$FPD $< 0$; positive means $\Delta$FPD $> 0$. The sign of y-axis: negative means motif disruption; positive means motif enhancing. Blue and red colored points are the concordant footprints passing FDR of 0.05.

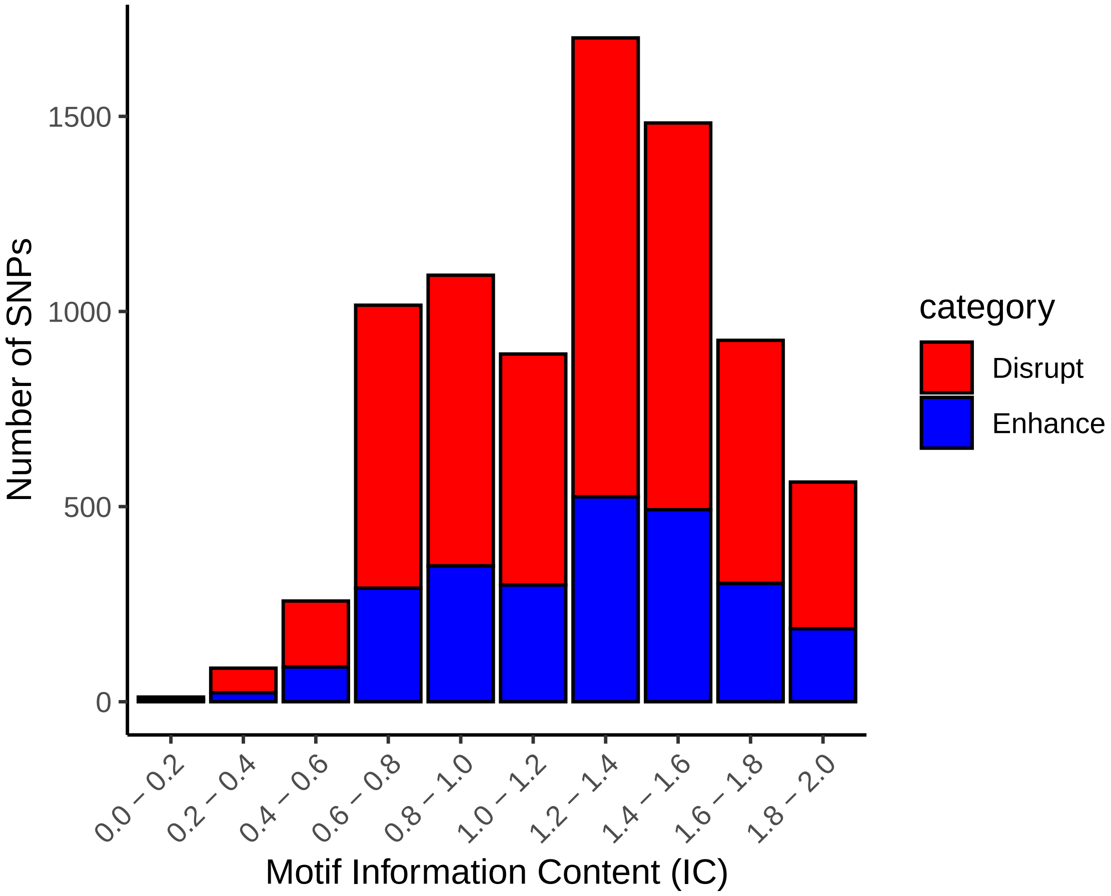

Fig. S23: Summary of the atSNP results for individual footprints binned by motif information content.

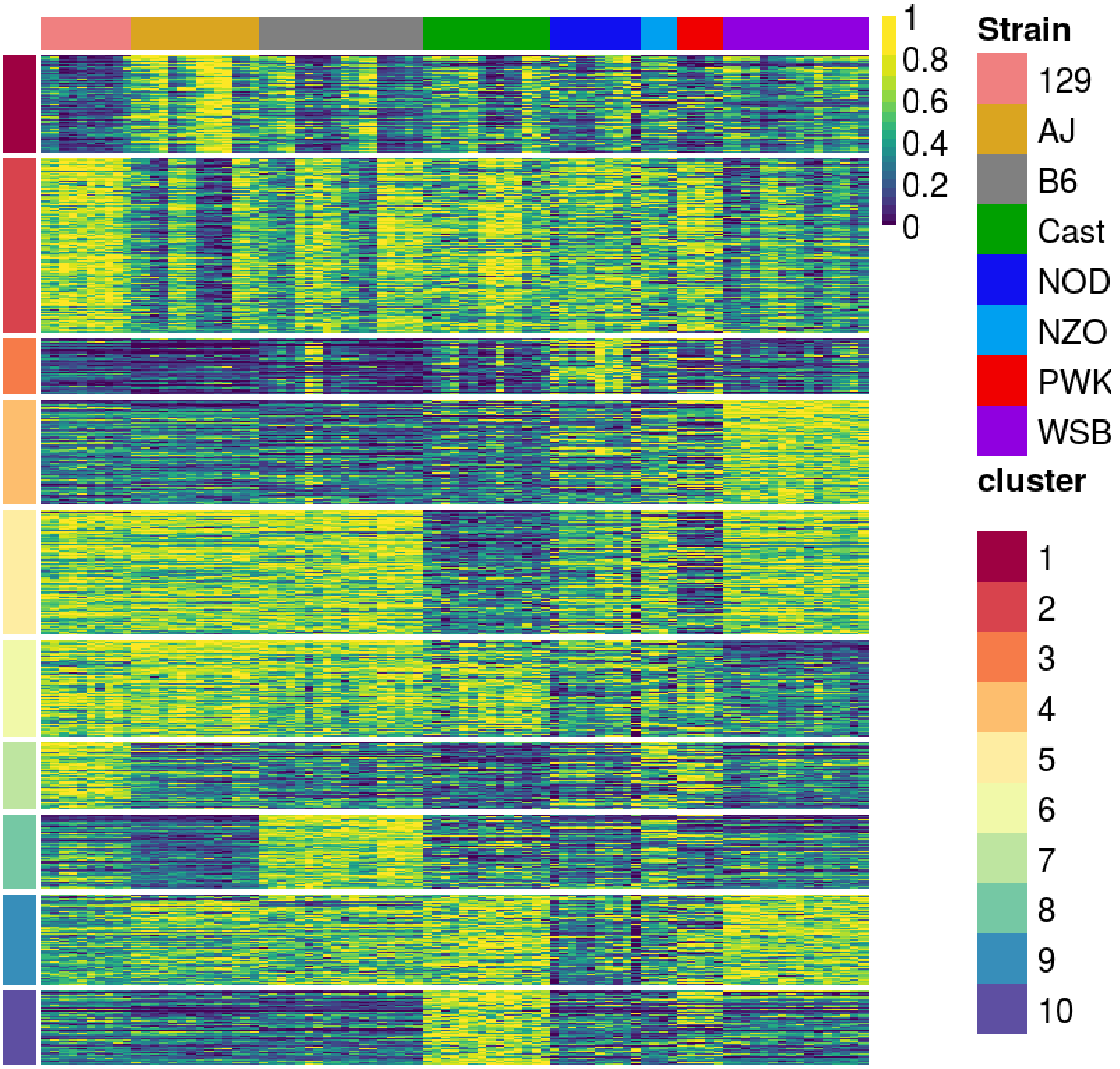

Fig. S24: Heatmap of islet RNA-seq data from founder DO strains: row and columns represent genes and samples, respectively. Transcripts of genes (rows) across samples are standardized to [0, 1] and are clustered by using k-means with k = 10. Columns are not clustered.

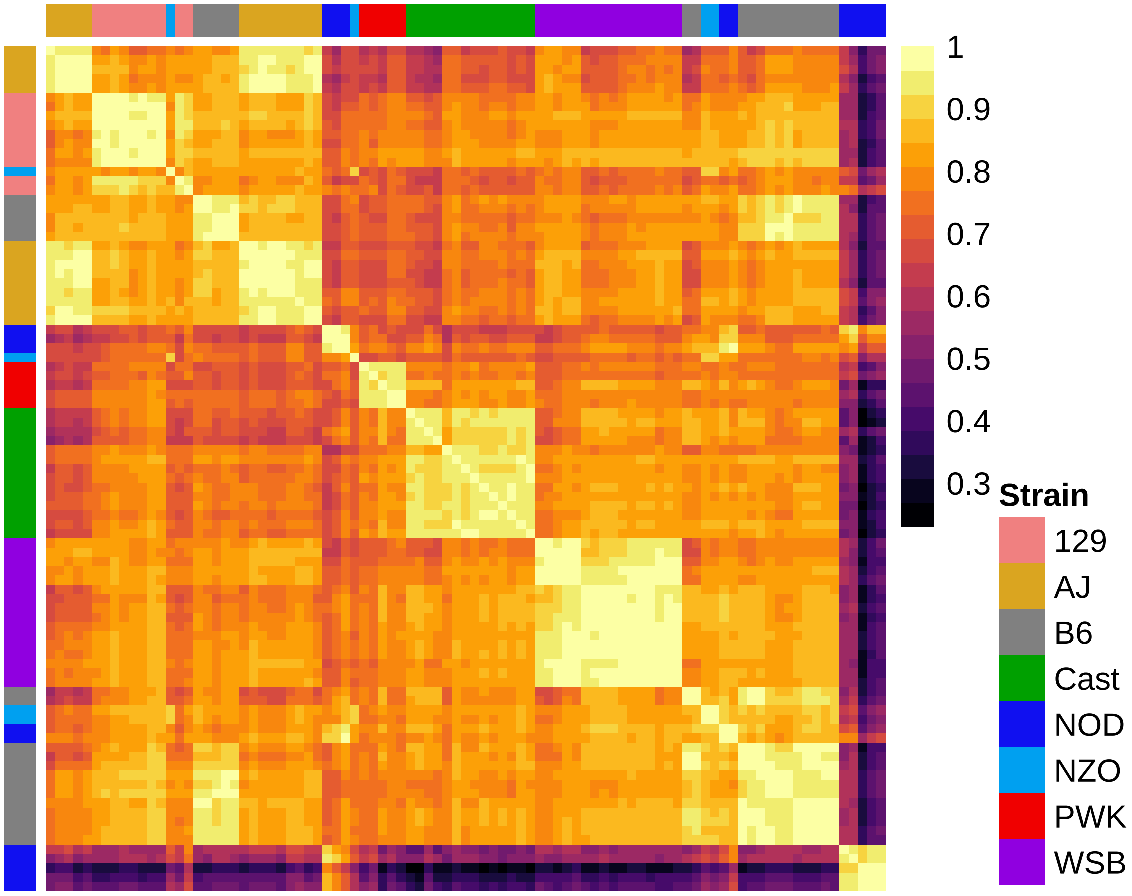

Fig. S25: Heatmap of pairwise Pearson correlations between 91 islet RNA-seq samples from DO founder strains. Rows and columns are organized hierarchically.

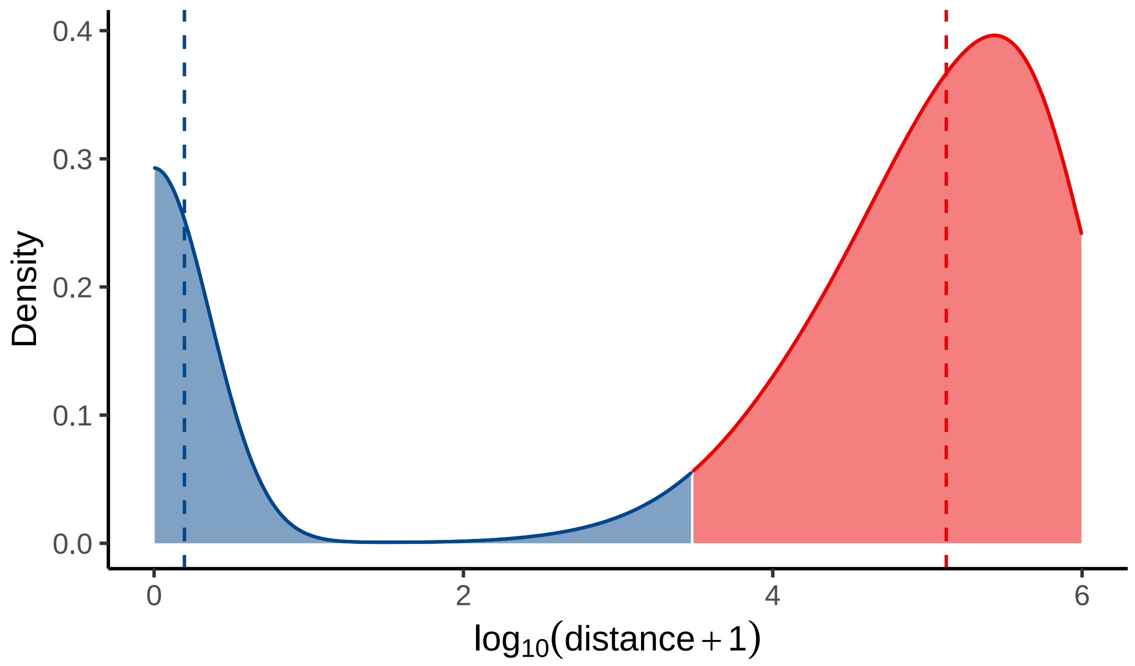

Fig. S26: The density of the genomic distances between the local-ATAC-MVs and the transcription start sites (TSSs) of the genes that they associate with. Blue and red shading denote promoter-proximal (distance $\leq$ 3 Kb) and promoter-distal ATAC-seq peaks harboring the local-ATAC-MVs (distance $>$ 3 Kb), respectively. The dashed lines represent the means for different groups.

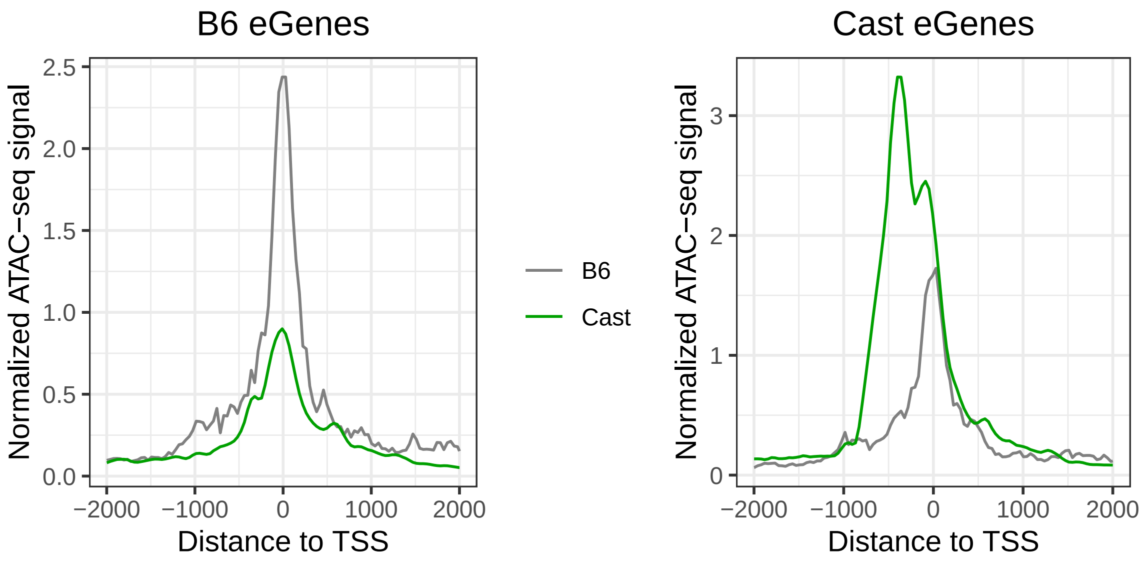

Fig. S27: `B6` and `CAST` normalized ATAC-seq signals at the TSSs of eGenes that are differentially expressed between the two strains. Left and right panels depict eGenes for `B6` (expressed more in `B6`) and `CAST` (expressed more in `CAST`), respectively.

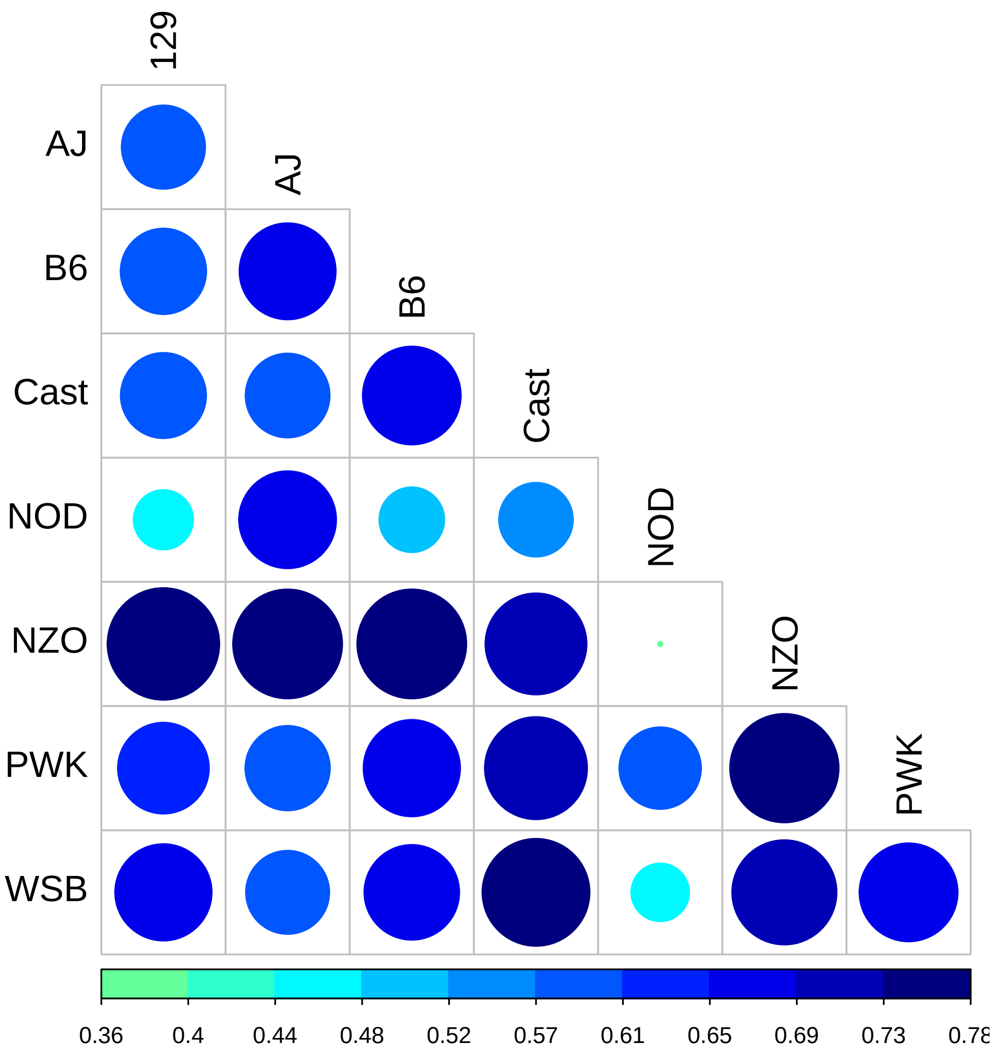

Fig. S28: Proportion of concordant eGenes across strains, i.e., differentially expressed genes with agreement between the direction of higher expression and higher TSS accessibility.

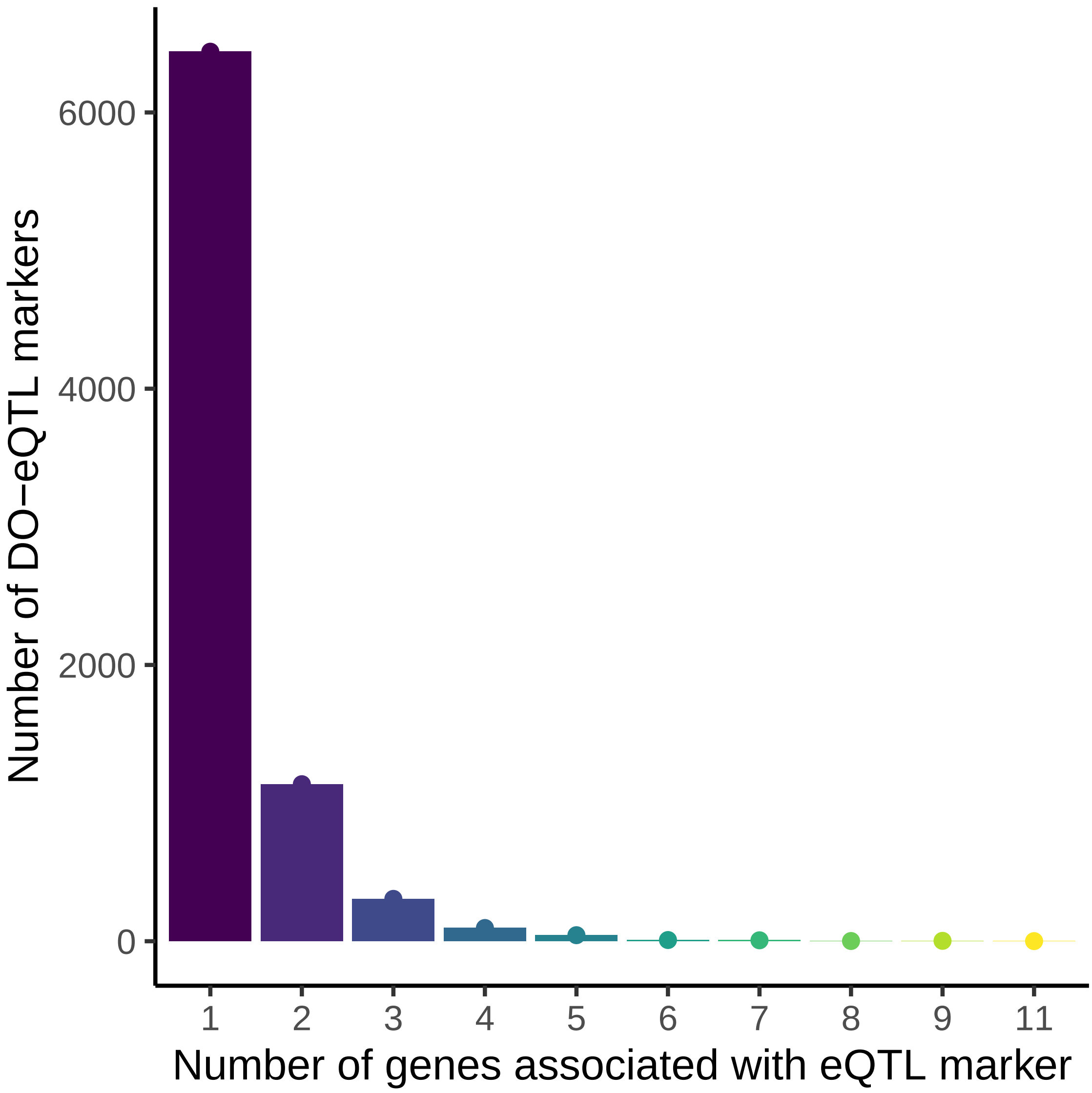

Fig. S29: Number of DO-eQTL markers (y-axis) vs. the number of DO genes a marker is associated with (x-axis). Most eQTL markers are associated with only one gene.

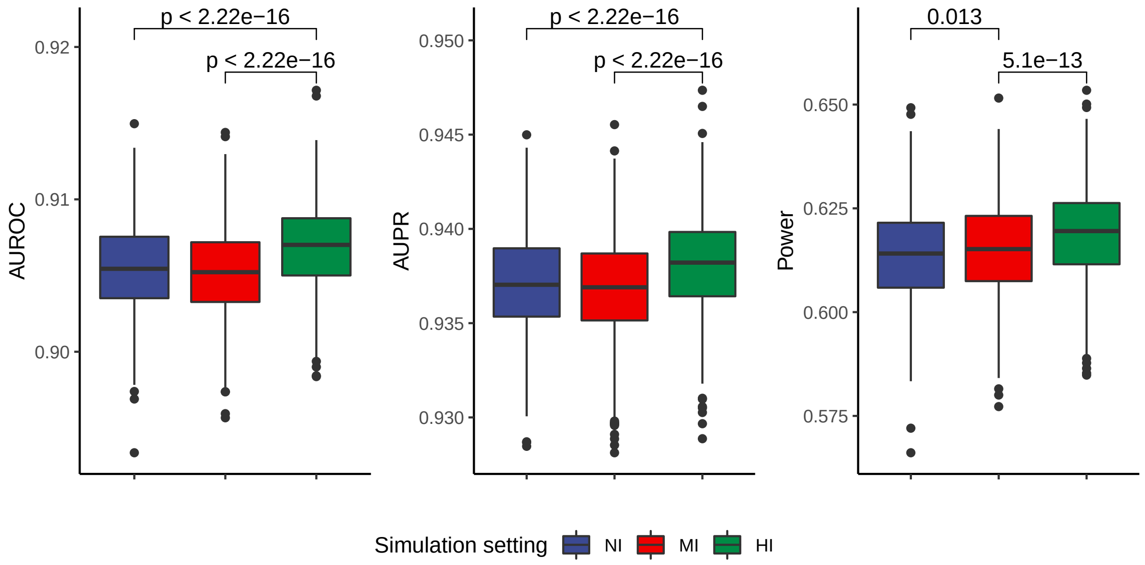

Fig. S30: INFIMA model performance quantified by inferred posterior probabilities of having a causal SNP across all the genes, i.e., $\hat{V}_g, \cdots, g = 1, \cdots, G$. Statistically significant Wilcoxon rank sum tests are highlighted at the top.

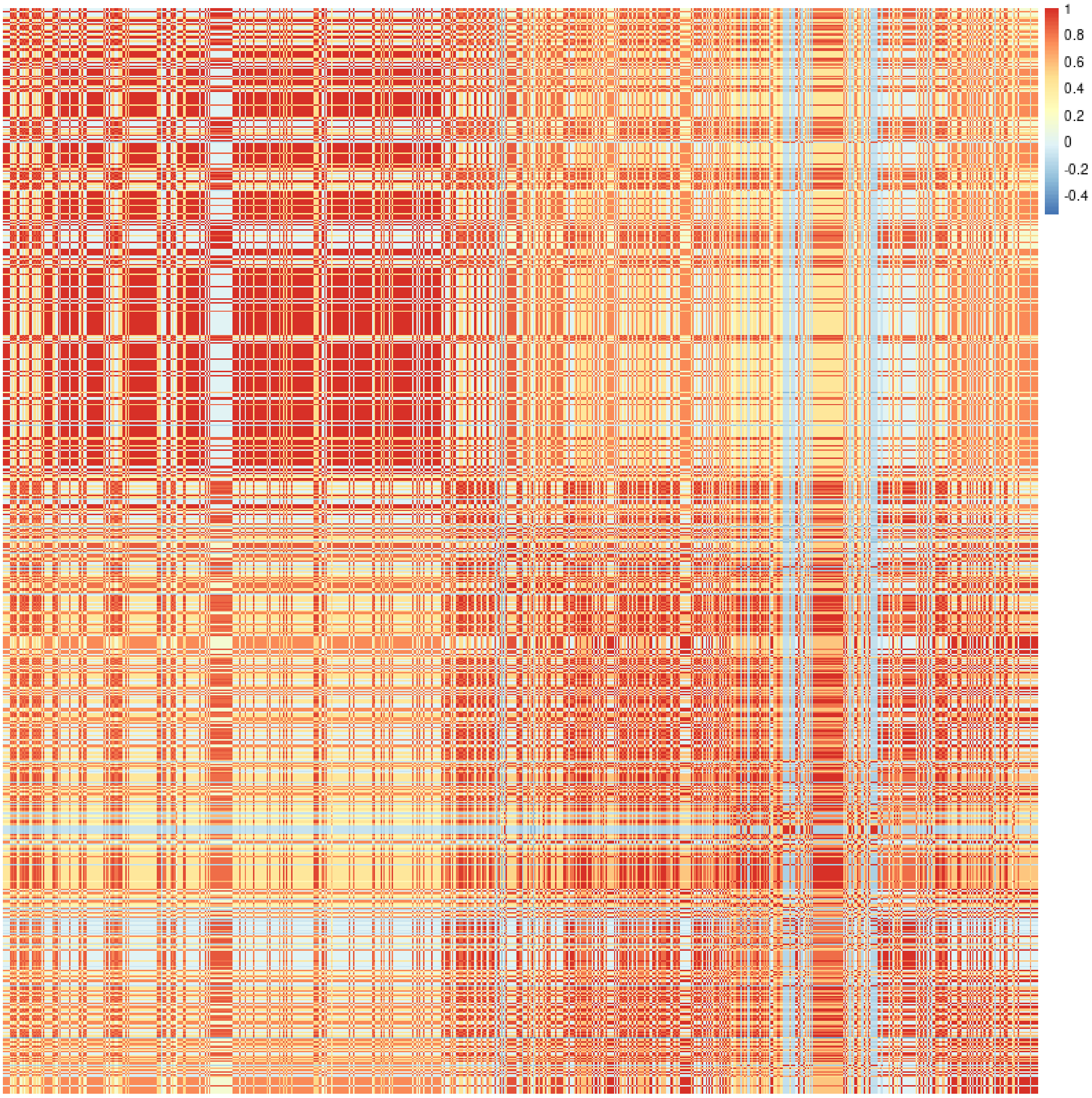

Fig. S31: The DO mice LD matrix of the imputed SNPs at the *Adcy5* locus. Row and columns depict the 4,616 SNPs within the 200 Kb of the eQTL marker and are ordered with respect to their genomic coordinates. Red color indicates a perfect LD.

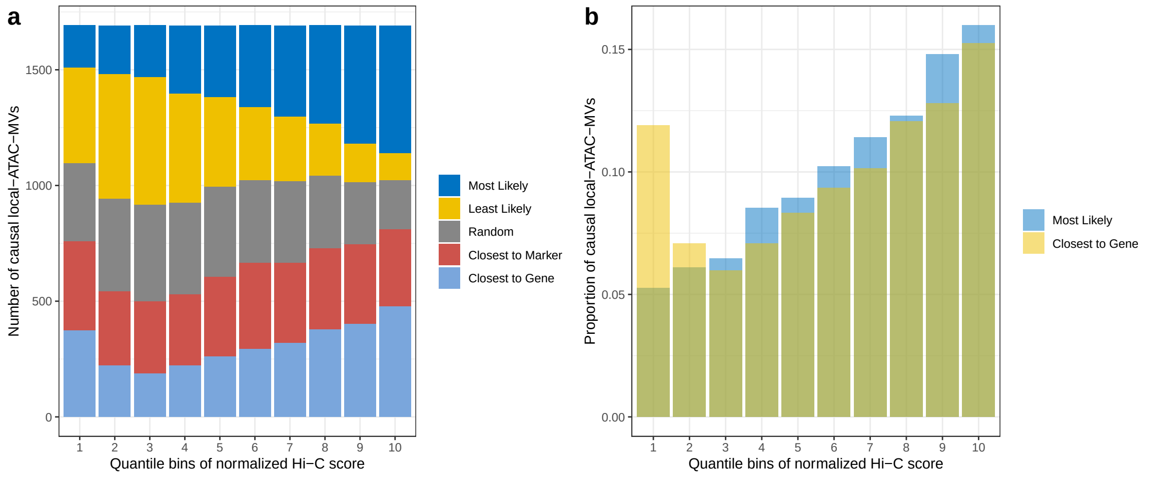

Fig. S32: **a.** Evaluation of fine-mapping strategies by counting the number of causal local-ATAC-MVs in each Hi-C quantile bin. **b.** A direct comparison between INFIMA ”Most Likely” and ”Closest to Gene”.

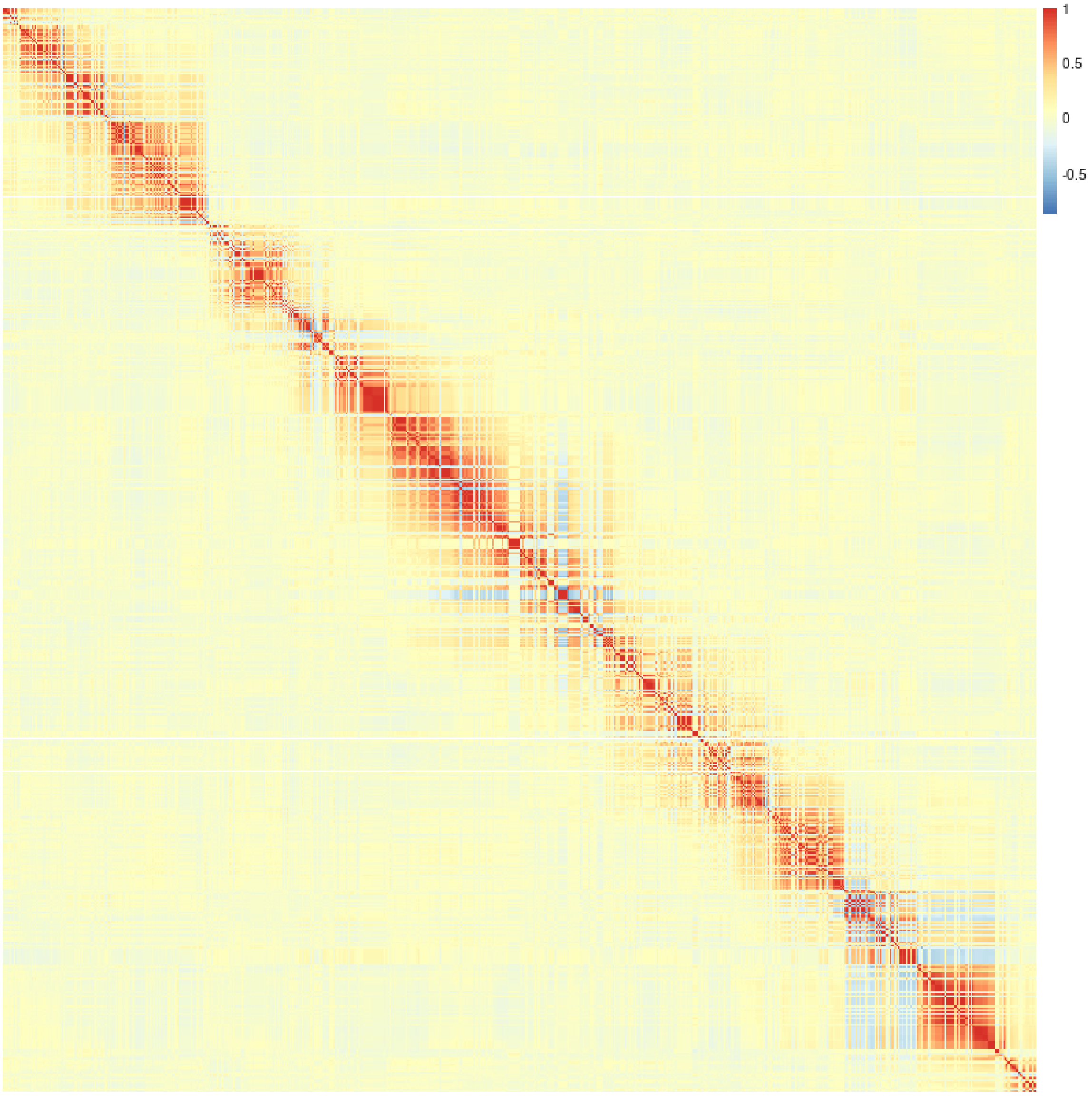

Fig. S33: The DO mice LD matrix for the 3,671 local-ATAC-MVs from chromosome 1. Rows and columns depict local-ATAC-MVs and are ordered with respect to their genomic coordinates. Red color indicates a perfect LD.

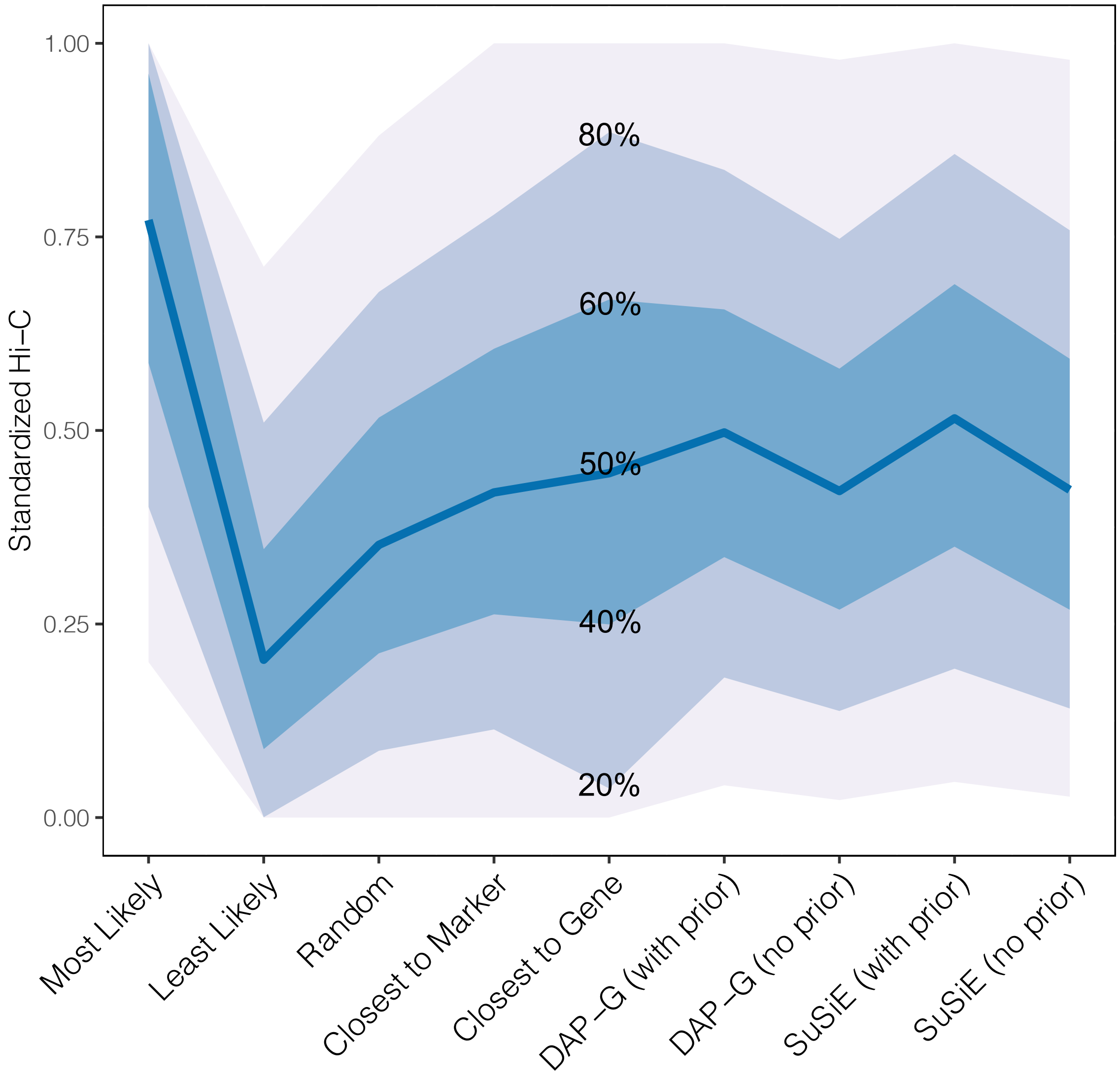

Fig. S34: Standardized Hi-C scores for the inferred causal local-ATAC-MVs by INFIMA and the competing approaches. Shaded areas mark different percentiles of normalized scores: Light blue: 20% and 80% percentiles; Blue: 40% and 60% percentiles; Dark blue: median.

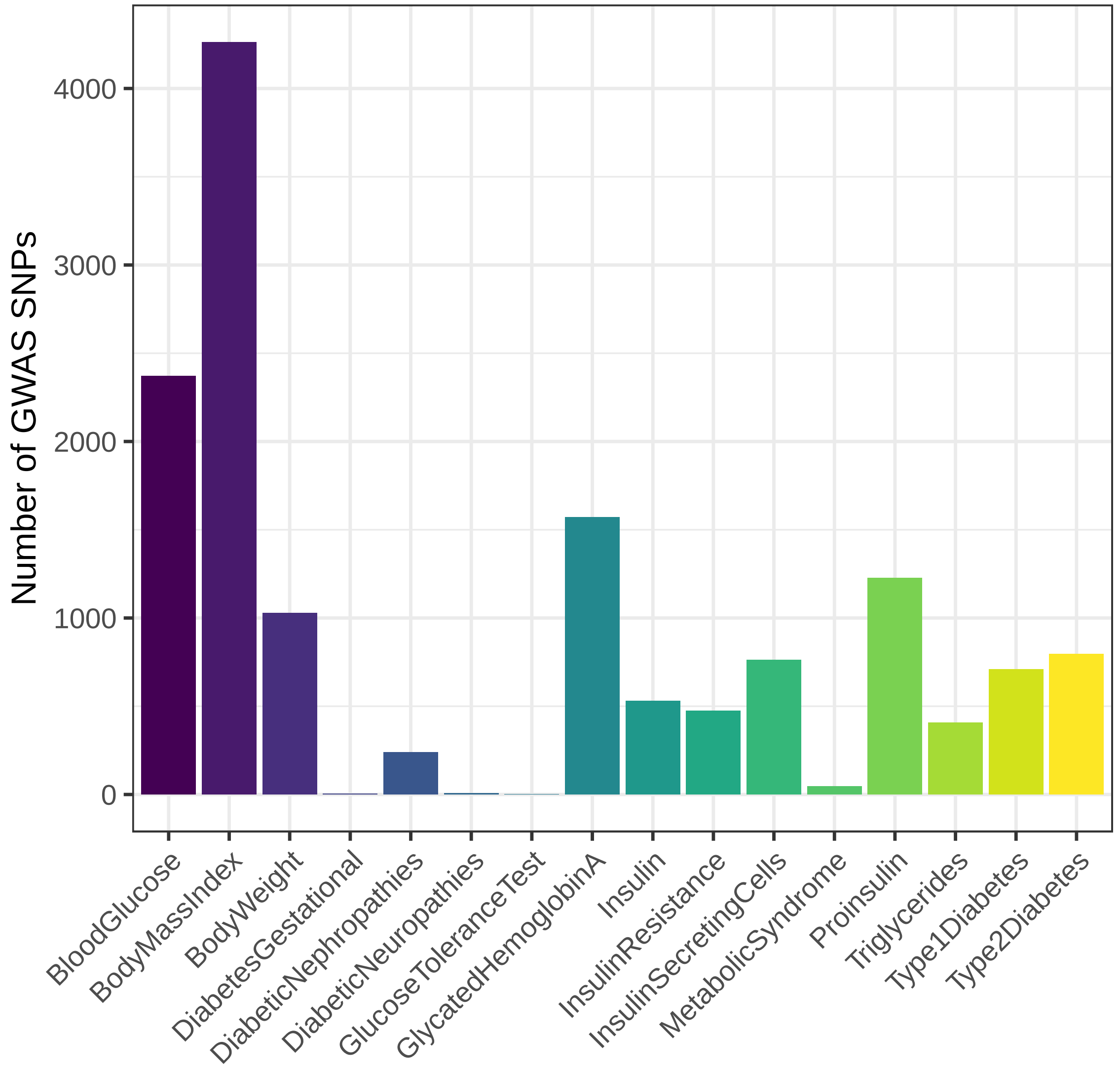

Fig. S35: An overview of the number of GWAS SNPs for 16 diabetic traits.

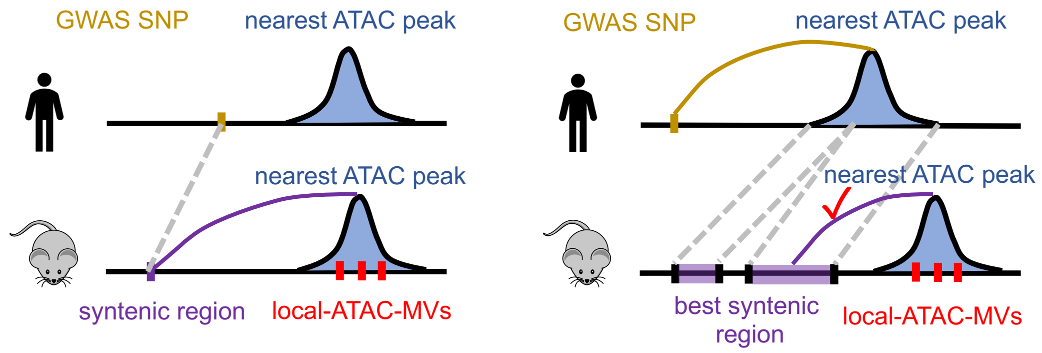

Fig. S36: Mapping scheme between human and mouse: direct lift-over (left); ATAC-seq peak-based lift-over (right).

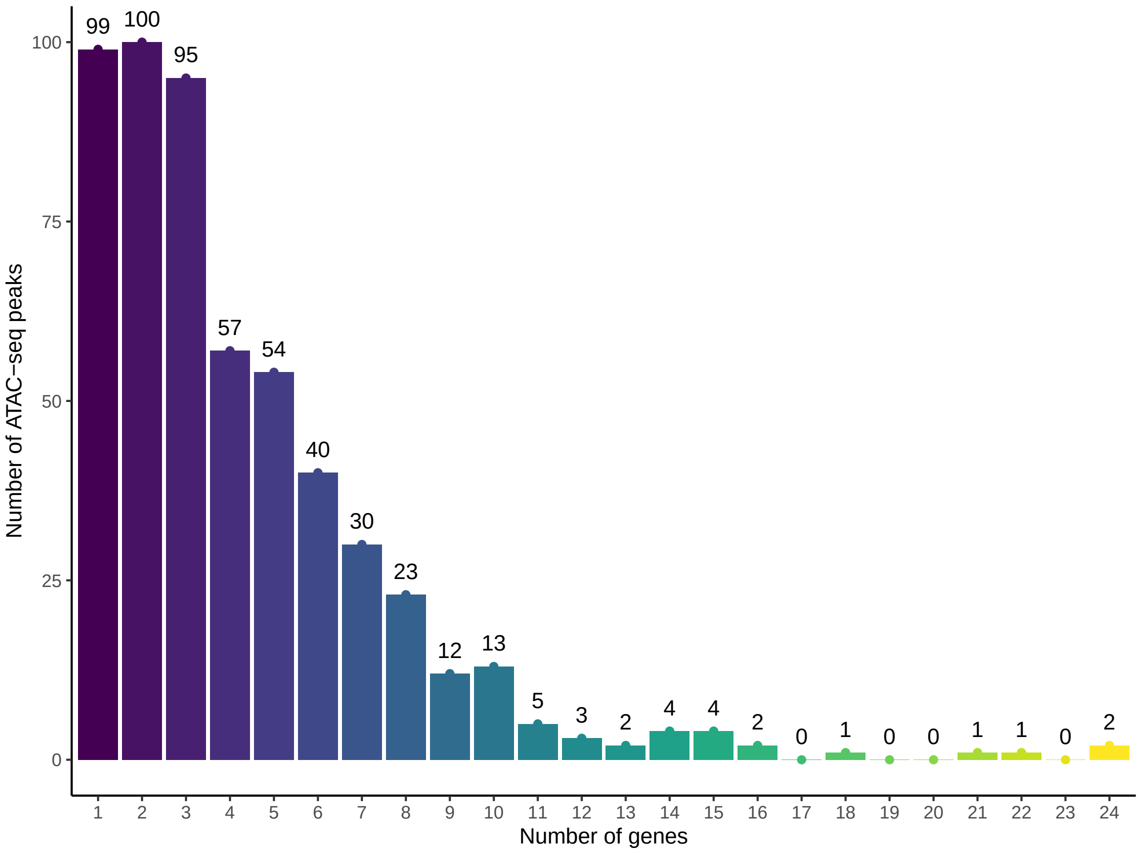

Fig. S37: Number of DO-eQTL genes an ATAC-seq peak is linked to by INFIMA.

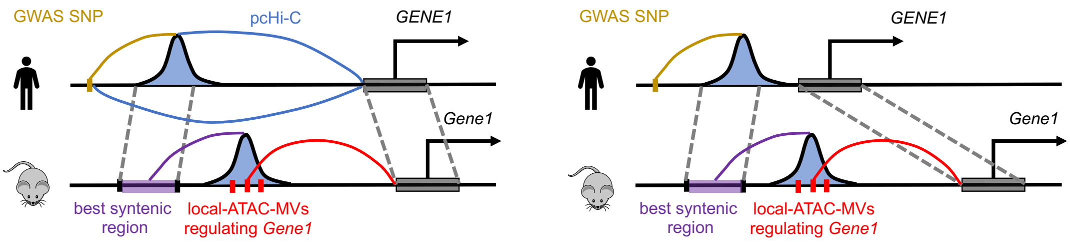

Fig. S38: Validation scheme with promoter capture Hi-C (pcHi-C) data. Left: GWAS SNP and/or human ATAC-seq peak at enhancer region in contact with orthologous gene promoter. Right: GWAS SNP and/or human ATAC-seq peak at promoter region of the orthologous gene. pcHi-C assays excludes short range interactions between a gene and its own promoter (Right panel).

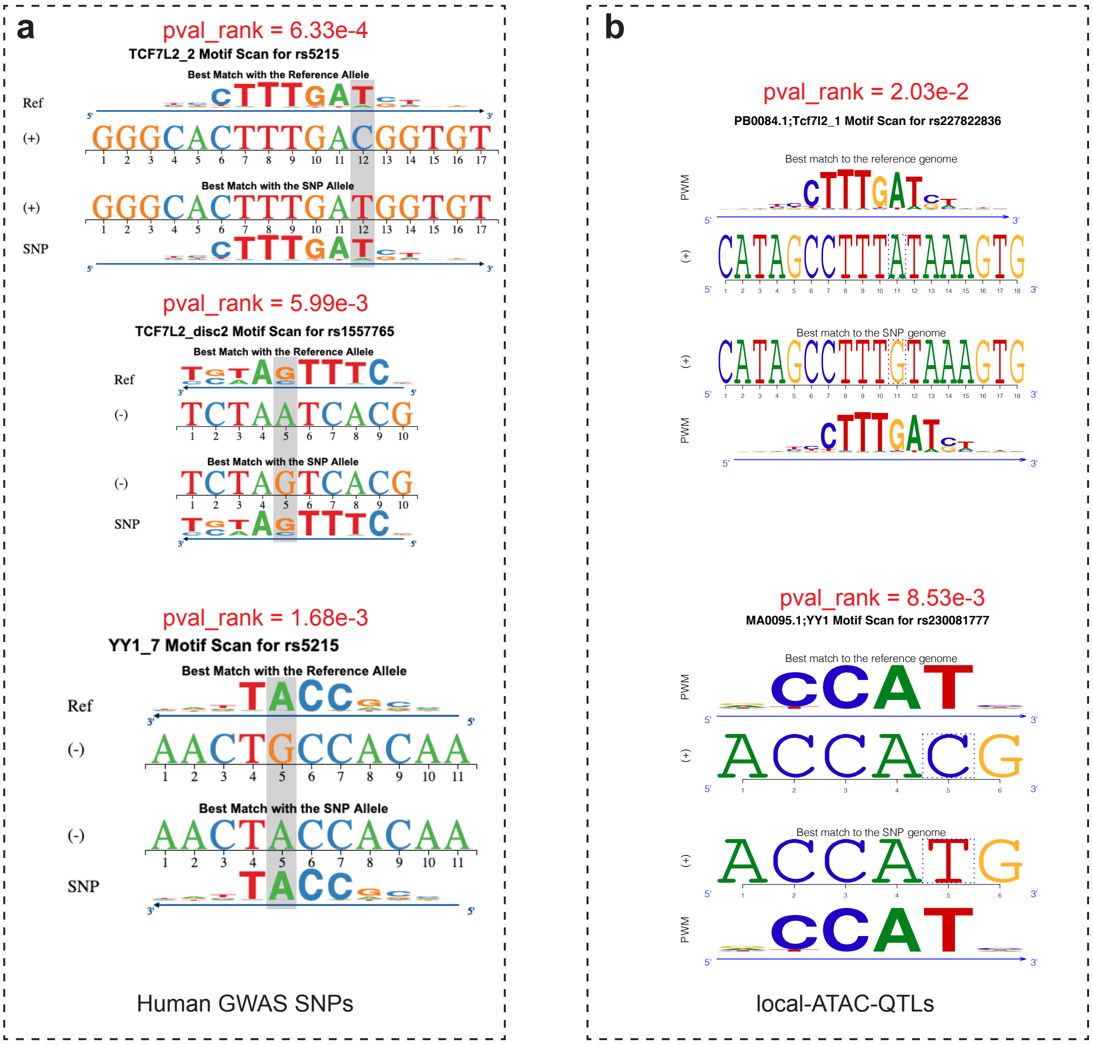

Fig. S39: *ABCC8* orthologous atSNP results. Left: human GWAS SNPs. Right: mouse local-ATAC-MVs.

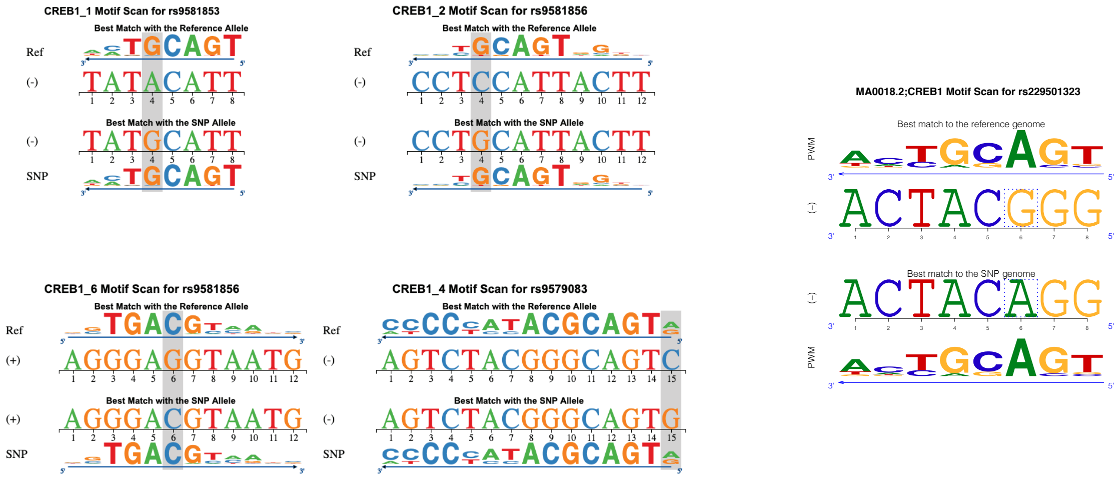

Fig. S40: *PDX1* orthologous atSNP results. Left: human GWAS SNPs. Right: mouse local-ATAC-MVs.

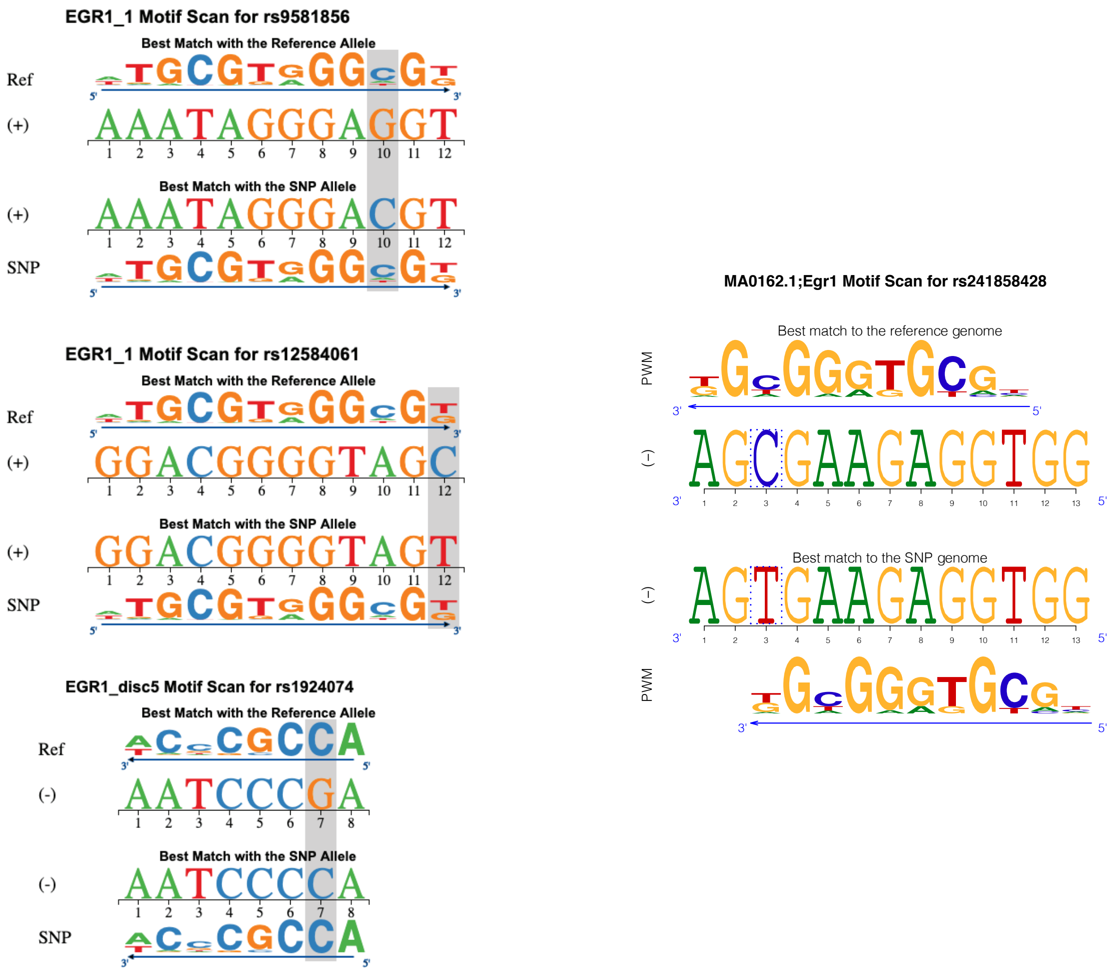

Fig. S41: *PDX1* orthologous atSNP results. Left: human GWAS SNPs. Right: mouse local-ATAC-MVs.

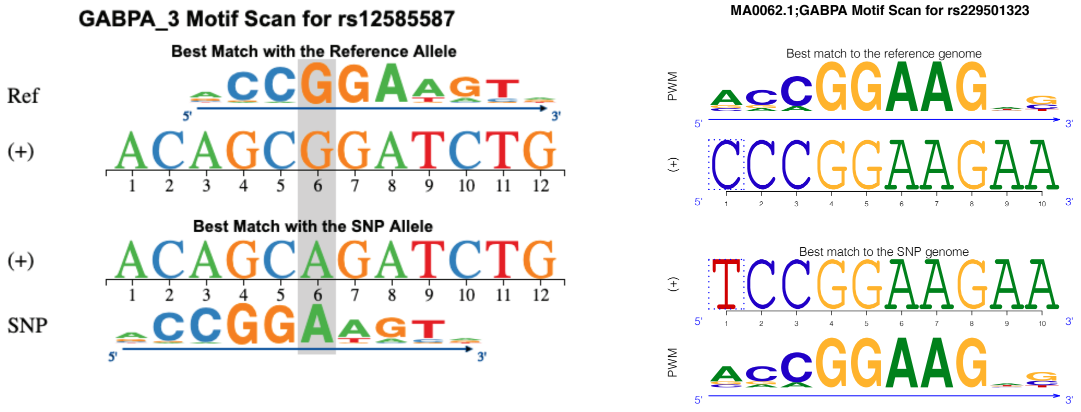

Fig. S42: *PDX1* orthologous atSNP results. Left: human GWAS SNPs. Right: mouse local-ATAC-MVs.

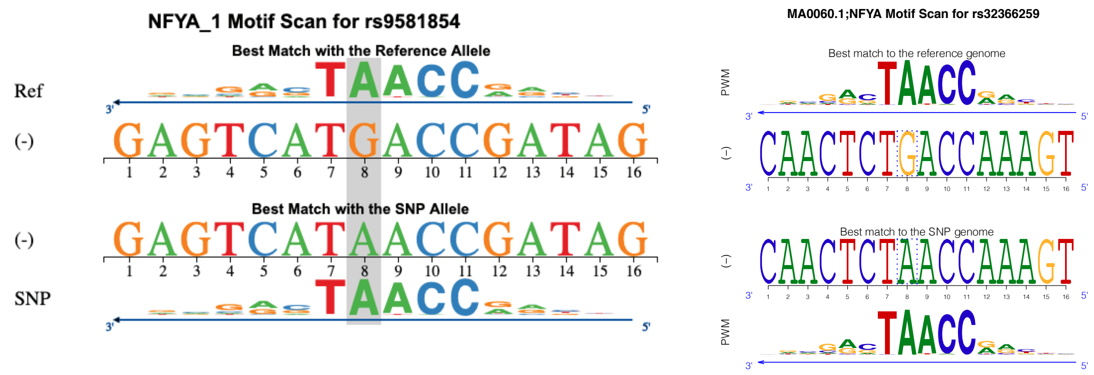

Fig. S43: *PDX1* orthologous atSNP results. Left: human GWAS SNPs. Right: mouse local-ATAC-MVs.

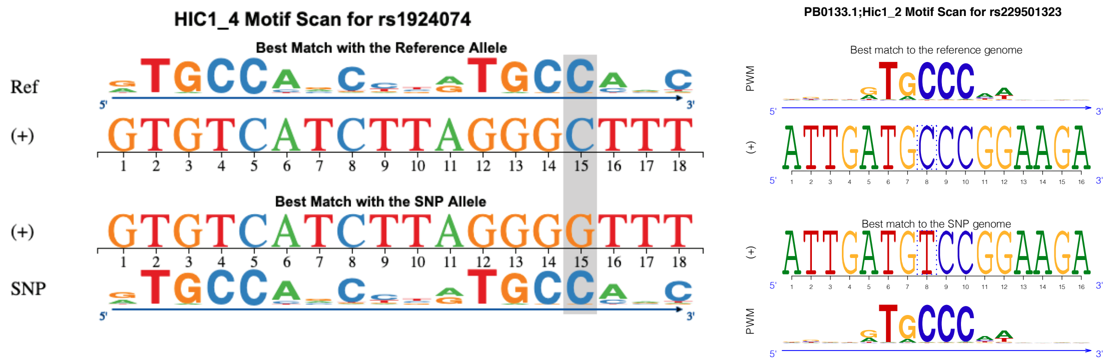

Fig. S44: *PDX1* orthologous atSNP results. Left: human GWAS SNPs. Right: mouse local-ATAC-MVs.

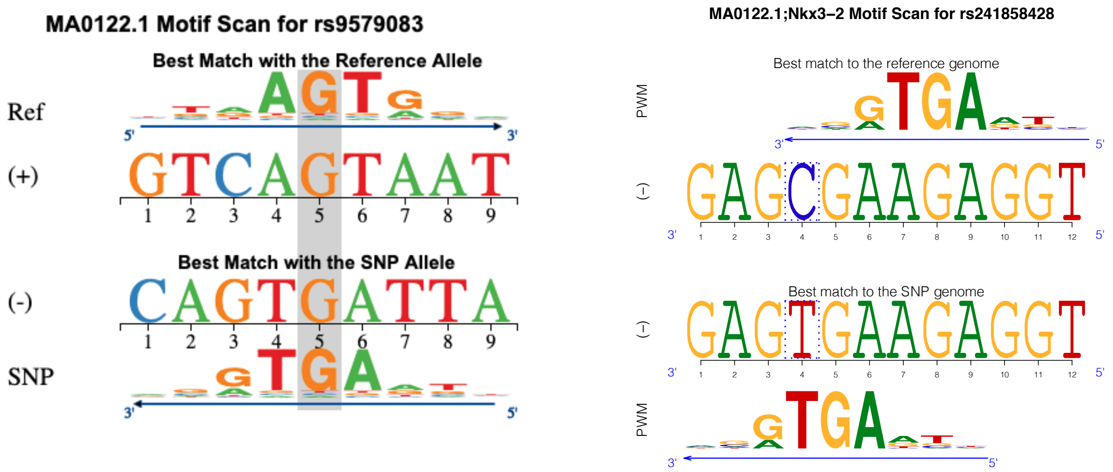

Fig. S45: *PDX1* orthologous atSNP results. Left: human GWAS SNPs. Right: mouse local-ATAC-MVs.

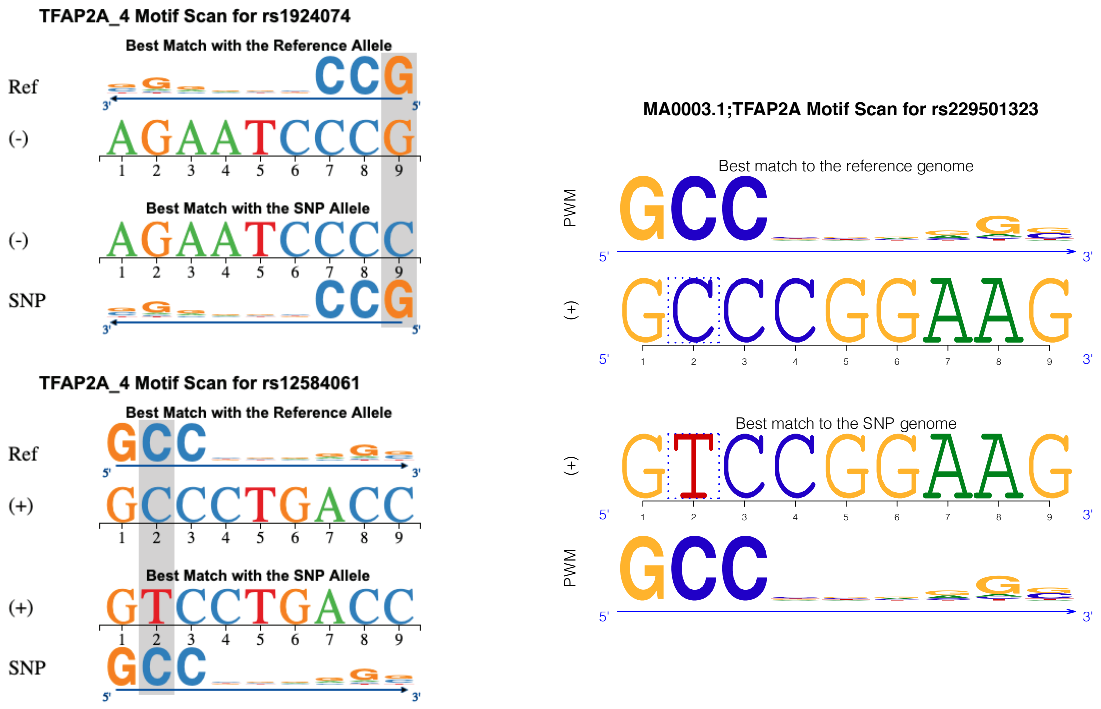

Fig. S46: *PDX1* orthologous atSNP results. Left: human GWAS SNPs. Right: mouse local-ATAC-MVs.

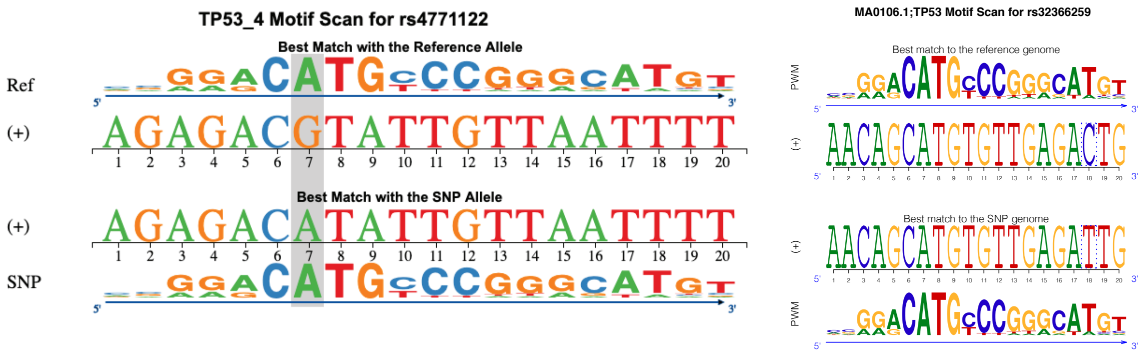

Fig. S47: *PDX1* orthologous atSNP results. Left: human GWAS SNPs. Right: mouse local-ATAC-MVs.

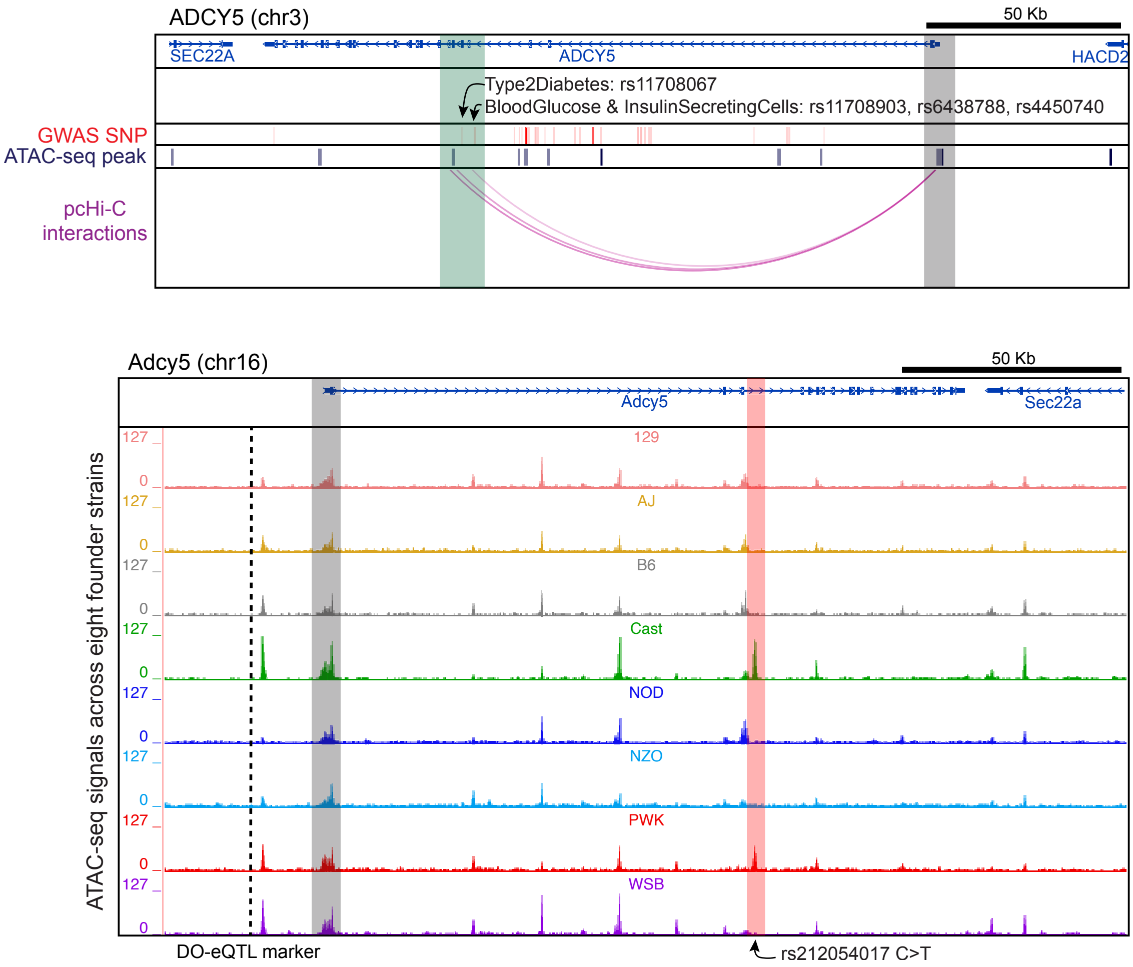

Fig. S48: Promoter capture Hi-C links *ADCY5* promoter to 4 GWAS SNPs which map to 1 mouse local-ATAC-MV with INFIMA predicted susceptibility gene *Adcy5*. Top: human genome depictions of intronic GWAS SNPs (translucent green) interactions with the *ADCY5* promoter (translucent gray), together with the human ATAC-seq peaks. Bottom: mouse genome depictions of ATAC-seq signal for local-ATAC-MVs (translucent red) where INFIMA fine-maps DO-eQTL marker (dashed line) linked to *Adcy5* (promoters highlighted in translucent gray).

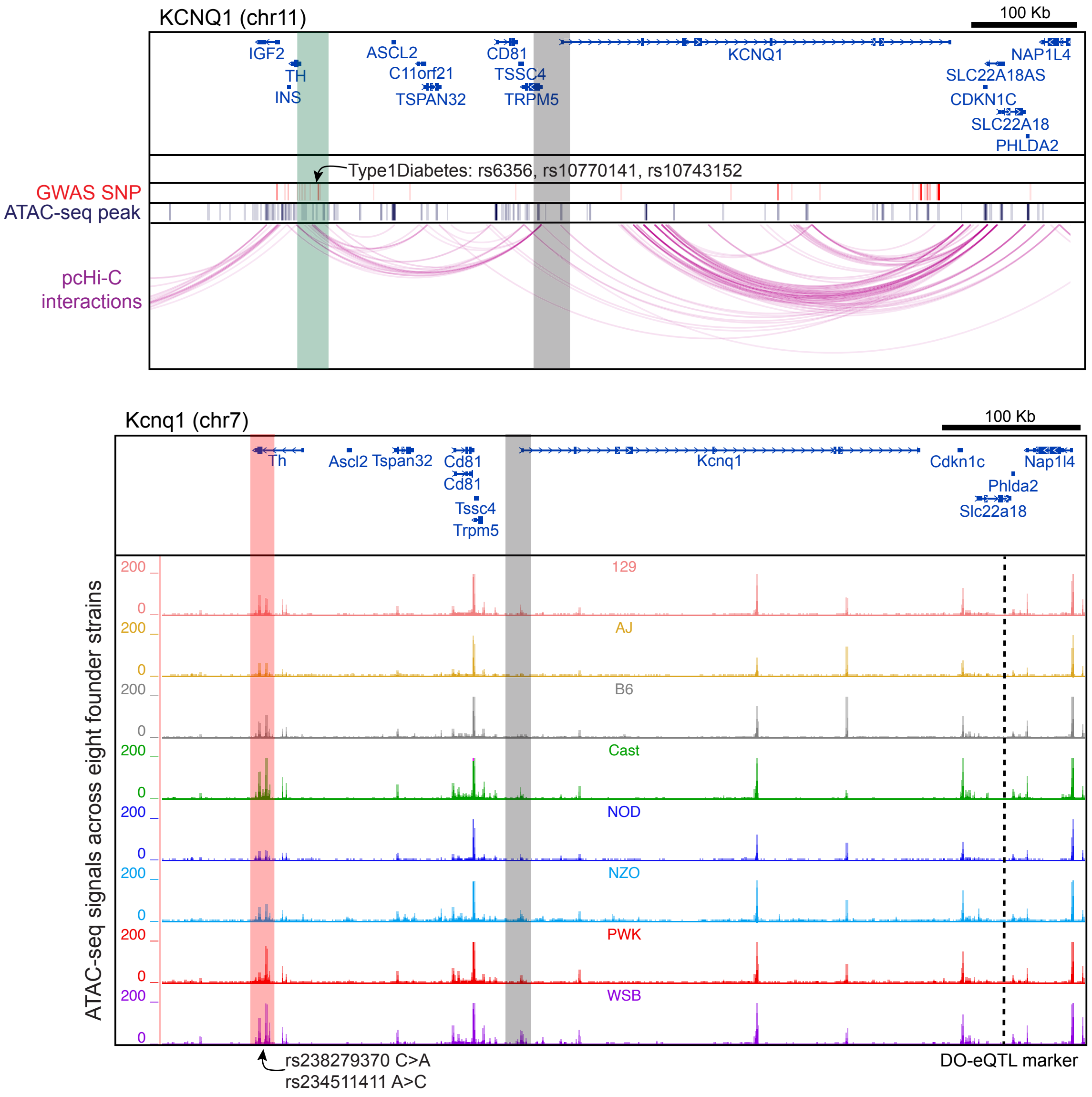

Fig. S49: Promoter capture Hi-C links *KCNQ1* promoter to 3 GWAS SNPs which map to 2 mouse local-ATAC-MV with INFIMA predicted susceptibility gene *Kcnq1*. Top: human genome depictions of *TH* promoter or distal GWAS SNPs (translucent green) interactions with the *KCNQ1* promoter (translucent gray), together with the human ATAC-seq peaks. Bottom: mouse genome depictions of ATAC-seq signal for local-ATAC-MVs (translucent red) where INFIMA fine-maps DO-eQTL marker (dashed line) linked to *Kcnq1* (promoters highlighted in translucent gray).

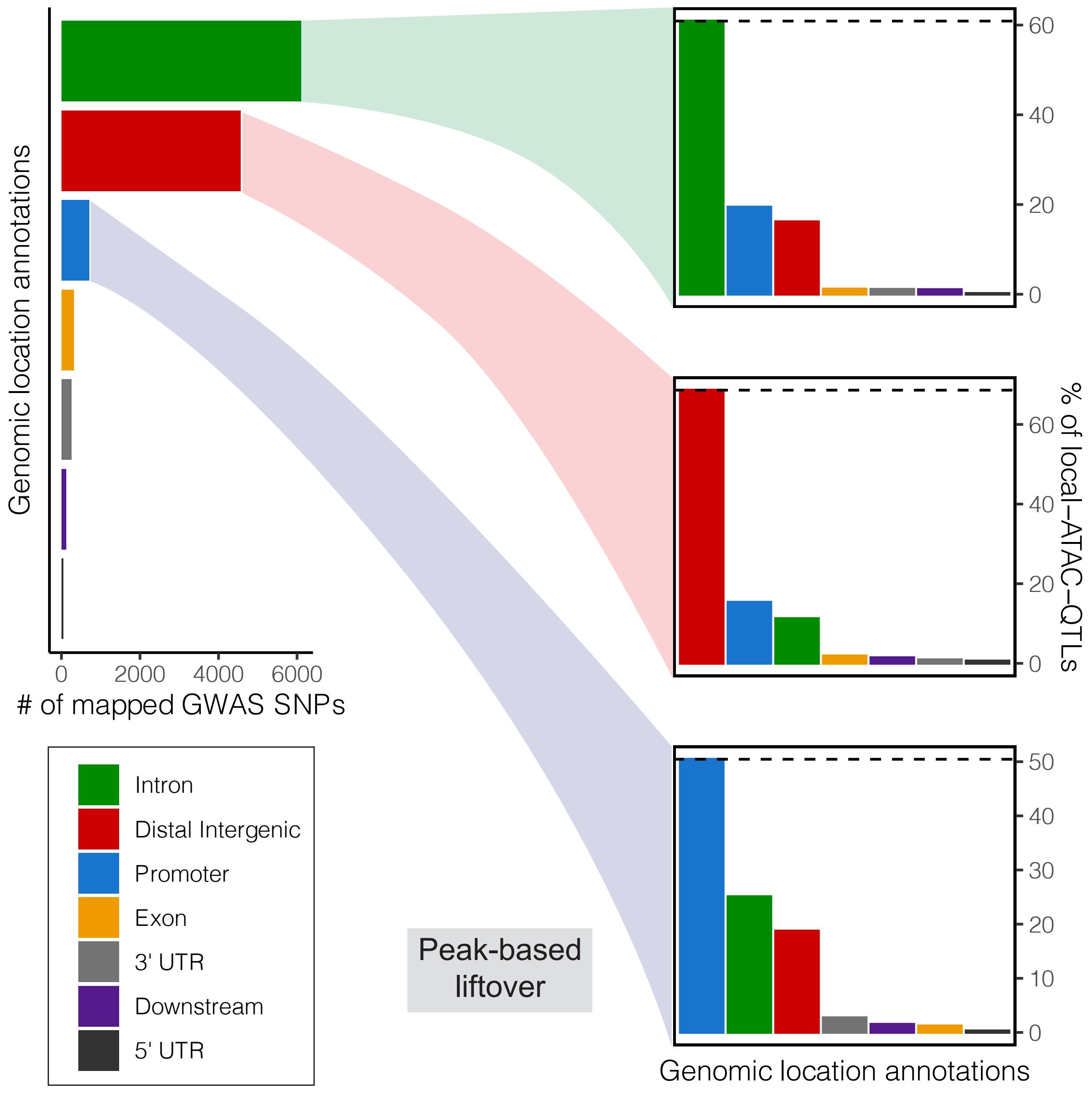

Fig. S50: Comparison of genomic location annotations between human GWAS SNPs and the orthologous mouse genetic variants. Left: numbers of mapped GWAS SNPs within intronic, distal, promoter, exonic, UTR and downstream genomic locations. Right: Barplots of the local-ATAC-MVs mapping to the intronic, distal and promoter groups of GWAS SNPs, highlighting marked conservation of genomic location types. Note that the mapping here is without the 10 Kb distance constraint in Fig. 8a. The level of genomic compartment conservation declines when removing the distance constraint.
